# Production of 130 diterpenoids by combinatorial biosynthesis in yeast

**DOI:** 10.1101/2023.03.24.534067

**Authors:** Ulschan Bathe, Jürgen Schmidt, Andrej Frolov, Alena Soboleva, Oliver Frank, Corinna Dawid, Alain Tissier

**Author notes:** Supporting information for this manuscript is provided at the end of the main text.

## Abstract

Diterpenoids form a diverse group of natural products, many of which are or could become pharmaceuticals or industrial chemicals. However, low concentration, presence in complex mixtures and challenging synthesis often limit their exploitation. The modular character of diterpene biosynthesis and the substrate flexibility of the enzymes involved make combinatorial biosynthesis a promising approach. Here, we report on the assembly in yeast of 130 diterpenoids by pairwise combinations of ten diterpene synthases producing (+)-copalyl diphosphate-derived backbones and four cytochrome P450 enzymes (CYPs); 80 of these diterpenoids have not yet been reported. The CYPs accepted the majority of substrates they were given but remained regioselective. Our results bode well for the systematic exploration of diterpenoid chemical space using combinatorial assembly in yeast.

## Introduction

Terpenoids are the largest class of natural products and include various important pharmaceuticals such as the anticancer Taxol and the antimalarial artemisinin^[1]^. Among them, diterpenoids are C-20 compounds made from four isoprene units and occur in plants, insects, bacteria and fungi^[2]^. It is estimated that there are >18,000 known diterpene structures and that they derive from 126 different backbones^[3]^. Despite this tremendous chemical diversity, the potential of diterpenoids is not fully exploited due to low concentrations in their natural hosts and/or the difficulty to access or cultivate the organisms that produce them. Furthermore, their complex structures make chemical synthesis highly challenging. Recent progress in the elucidation of diterpenoid biosynthesis pathways offers the opportunity to use metabolic engineering in appropriate hosts to produce these compounds^[4]^. Diterpene biosynthesis starts with geranylgeranyl diphosphate (GGPP) as general precursor. First, diterpene synthases (diTPS) convert GGPP to the diverse diterpene skeletons^[5]^. There are two major classes of diTPS. Class II diTPS initiate cyclization by protonation at the C14-C15 double bond of GGPP and typically generate bicyclic diterpenyl-diphosphates such as *ent*-copalyl diphosphate (*ent*-CPP), the precursor of gibberellins ^[4a, 6]^. Class I diTPS cleave the diphosphate ester bond of GGPP, or of diterpenyl-diphosphates produced by class II diTPS, generating a carbocation that initiates a cyclization reaction^[6]^. In addition, bifunctional diTPS have both class I and class II activity, and can perform two cyclizations in succession ^[4a, 6]^. The final products of diTPS are single or multiple diterpene olefins or alcohols of linear, macrocyclic or polycyclic structures. Downstream of diterpene skeletons, cytochrome P450 oxygenases (CYPs) play a crucial role by introducing oxygen groups and in some cases triggering backbone rearrangements, thereby greatly increasing the diversity of diterpenoids^[7]^. Many of the CYPs involved in diterpenoid biosynthesis are promiscuous, accepting not only their natural substrate but also structurally related compounds ^[7b]^. Modularity, multi-product formation by diTPS, and promiscuous CYPs are features that make diterpenoid biosynthesis highly amenable to combinatorial biosynthesis, which is a powerful approach to generate libraries of natural products, either existing or new-to-nature; the synthesized products can then be screened for biological activity. In the natural product field, there are successful demonstrations of combinatorial biosynthesis of polyketides, which include compounds with antitumor or antimicrobial activities^[8]^. For diterpenoids, the promiscuity of class I diTPS was harnessed by generating class II/class I combinations for the production of new-to-nature backbones^[9]^. Furthermore, specific diterpene backbones were combined with CYPs, either from the same or closely related organisms or that have a highly conserved function resulting in significant but limited extension of the chemical space^[10]^.

Here, we explore the potential of combinatorial assembly of diTPS and CYPs by combining 10 known diTPS that produce backbones derived from (*+*)-copalyl diphosphate ((*+*)-CPP; also called normal CPP) with one of four CYPs from distant plant species. Expression of all possible combinations in *Saccharomyces cerevisiae* resulted in the production of 130 different diterpenoids. 36 have already been described, 18 were determined by interpretation of mass spectra (see Fig. S13 to S36), and 2 by NMR (Fig. S1 to S12). For 80 of these diterpenoids no report could be found in the literature or in chemical databases.

## Results and Discussion

### Combinatorial biosynthesis of diterpene synthases in yeast

In recent years, more than 150 diTPS and 50 CYPs acting on diterpene backbones have been reported from plants, bacteria, and fungi ^[7b, 11]^. From this dataset, we selected ten plant diTPS that use (*+*)-CPP as substrate and/or produce diterpene skeletons of normal configuration (Fig. 1A, Table S1). Among them, four (AbCAS, SmCPSKSL1, GbLPS and PaLAS) are bifunctional, thus using GGPP as substrate, while OmTPS3, RoMiS, IrTPS2, IrKSL6, PbPIM and MsTPS1 are monofunctional class I diTPS acting on (*+*)-CPP. Using the *Golden Gate* modular cloning system we developed for yeast (Fig. 1B) ^[4d]^, we co-expressed a geranylgeranyl diphosphate synthase (GGPPS) and a (*+*)-CPP synthase from *Rosmarinus officinalis* (RoCPS). Together these constitute the core module (CM) which was assembled in a yeast expression vector with different combinations of class I diTPS (Table S2) ^[4d]^. When a bifunctional diTPS was expressed, RoCPS was replaced by a short non-coding nucleotide linker. As expected, the ten diTPS-only strains produced the diterpene skeletons as previously reported (Fig. 1, Table S1). However, in two cases we detected additional diterpenes not described before. PaLAS also synthesized sandaracopimara-8(14),15-diene, four unknown olefin diterpenes and a diterpene alcohol, and MsTPS1 produced an additional unknown olefin diterpene (Table S3).

**Figure 1.**
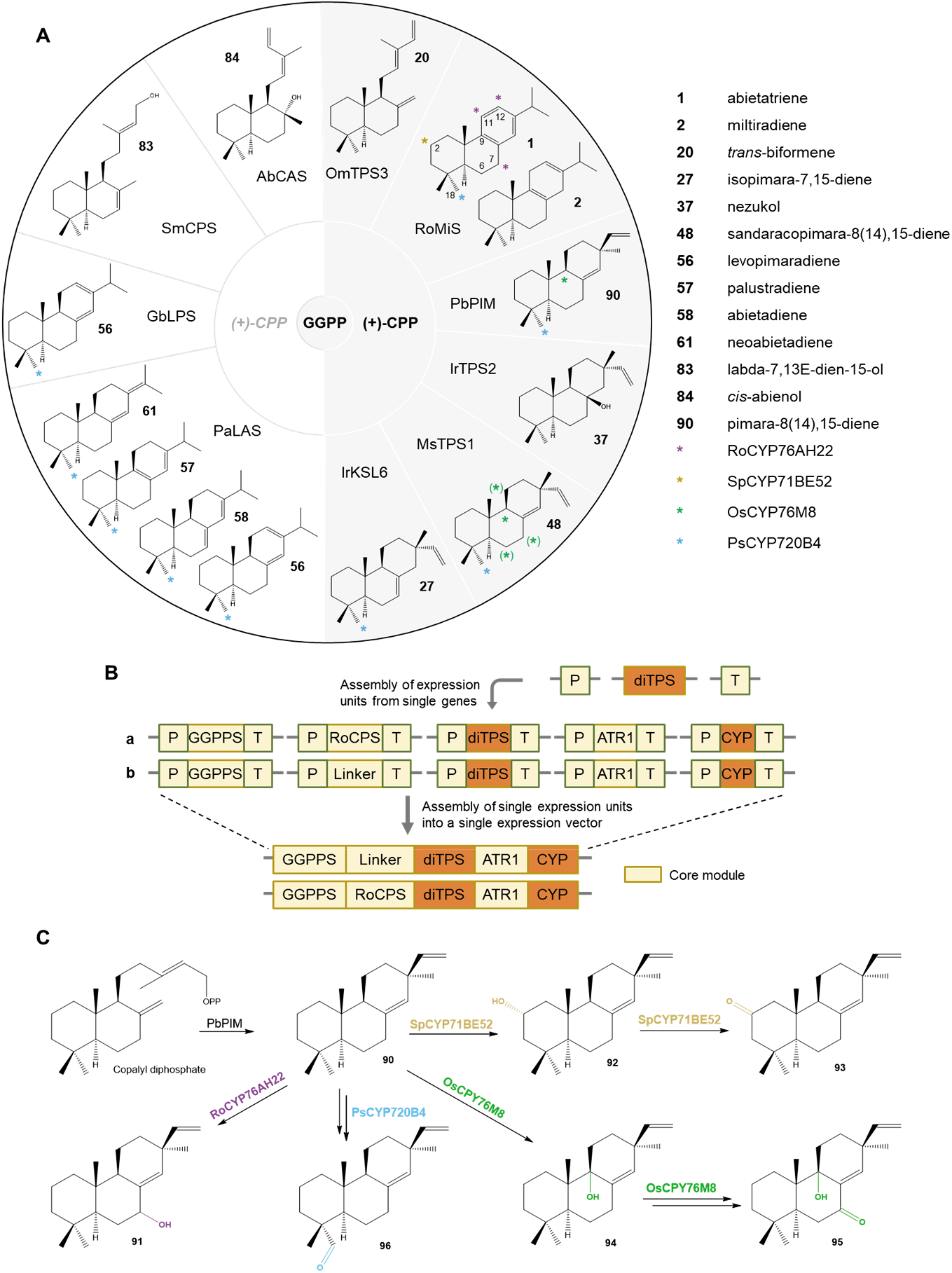
*Golden Gate* powered combinatorial biosynthesis of diterpenoids derived from (+)-CPP in yeast. A, diTPS that were selected for this study use either GGPP (left side) or (+)-CPP (right side) as substrate to produce the diterpene skeletons shown. They serve as substrates for RoCYP76AH22, OsCYP76M8, SpCYP71BE52 and PsCYP720B4 for oxidation at C2, C6, C7, C9, C11, C12 and C18 (highlighted with asterisks; asterisks in brackets refer to oxidations by OsCYP76M8 on substrates of *syn*- or *ent*-configuration). B, *Golden Gate* modular cloning developed for yeast allowed the expression of a core module (GGPPS, RoCPS or linker) and a CYP reductase (ATR1) with combinations of diTPS and CYPs (orange boxes). C, Expressed CYPs retain their regioselectivity on different substrates as exemplified for pimara-8(14),15-diene (**90**).

### Four cytochrome P450s give rise to 110 oxidized diterpenes

In order to expand the diversity of oxidized diterpenoids, we selected four CYPs from four distant plant species. Three of them, namely RoCYP76AH22, SpCYP71BE52, and PsCYP720B4, have diterpene backbones derived from (+)-CPP as natural substrates and one, OsCYP76M8, has diterpene backbones derived from *ent*- or *syn*-CPP as natural substrates but was also shown to react on diterpenes derived from (*+*)-CPP (Fig. 1A, Table S1) ^[4c, 4d, 10a, 12]^. We expected that the CYPs would accept slightly different substrates but would remain regio- and stereoselective for carbon positions C2, C6, C7, C9, C11, C12 and C18 (see Fig. 1A) compared to their known substrates^[13]^. The CM and a CYP reductase from *Arabidopsis thaliana* (ATR1) were assembled in a yeast expression vector with different combinations of diTPS and CYPs. Thus, we generated 40 strains that express both diTPS and a CYP and could therefore produce oxidized diterpenes (Table S2).

When co-expressing the four selected CYPs with diTPS that make the native CYP substrates (Fig. 1), oxidized diterpenoids were formed as described previously ^[10a, 10b, 12]^. We then analyzed the strains co-expressing each of the ten diTPS with each of the CYPs to provide the CYPs with substrates that differ from their natural ones. Of 40 enzyme combinations, 31 produced oxidized compounds, ranging from only one product up to 13 different oxidized diterpenoids (PaLAS + RoCYP76AH22) detected by high-resolution GC-MS and LC-MS (Fig. 2, Fig. 3, Table S3 and Table S4). The total number of diterpene olefins and oxidized diterpenes detected by GC-MS was 111, and 22 products were identified by LC-MS (Table 1, Table S3 and Table S4). Some combinations did not give rise to oxidized products. *trans*- biformene (**20**), nezukol (**37**) and *cis*-abienol (**84**) were not converted by some of the CYPs, while labda-7,13*E*-dien-15-ol (**83**) was not accepted by any of the CYPs (Table 1).

**Table 1.**
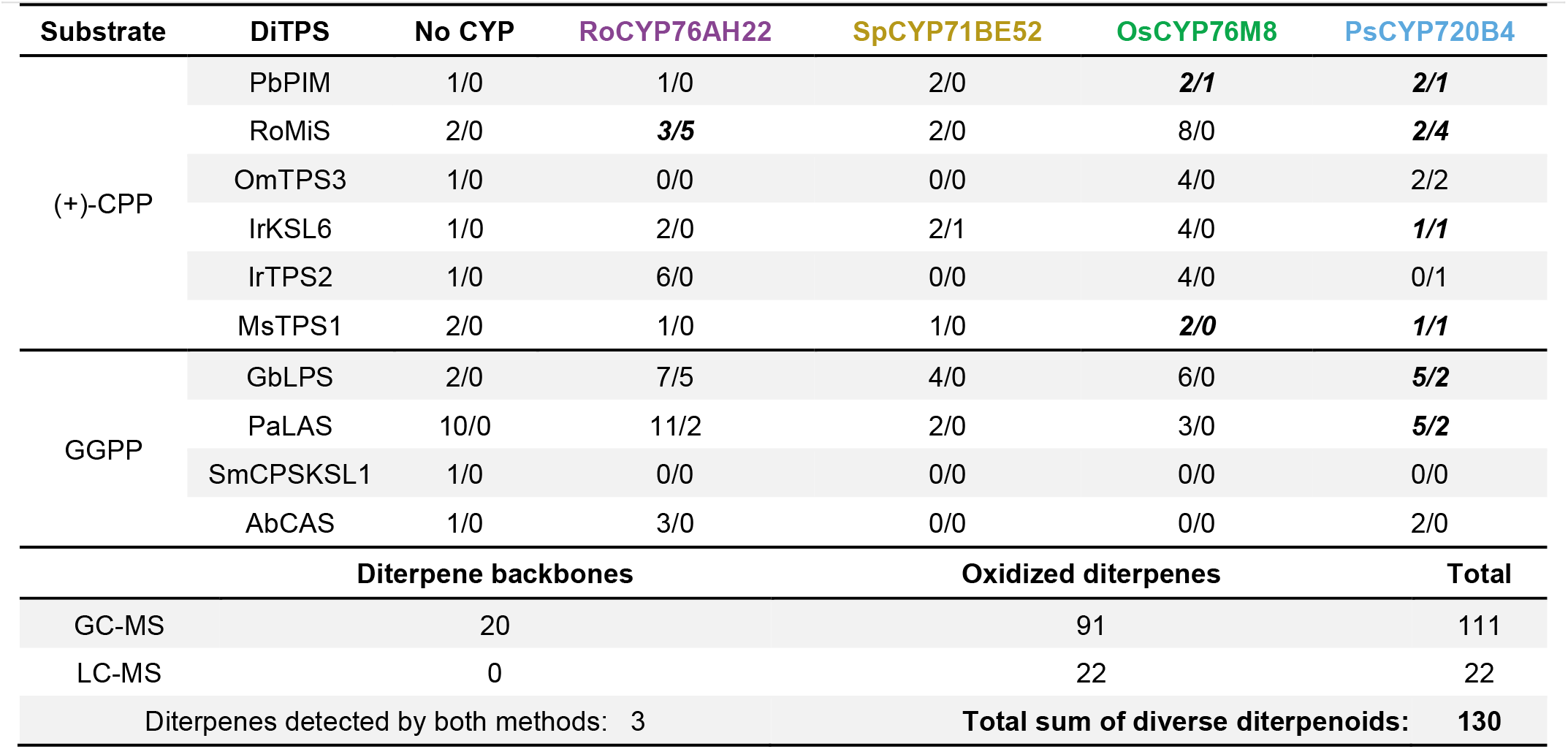
Number of diterpenes produced in yeast. Diterpene backbones formed by the activity of diTPS (“no CYP”) and oxidized diterpenoids that resulted from co-expression of diTPS and given CYP (colored columns) were detected by GC-MS or LC-MS (GC/LC). Some oxidized diterpenoids were detected by both methods. Enzyme combinations in which the diTPS provide the natural CYP substrate are given in bold-italic.

**Figure 2.**
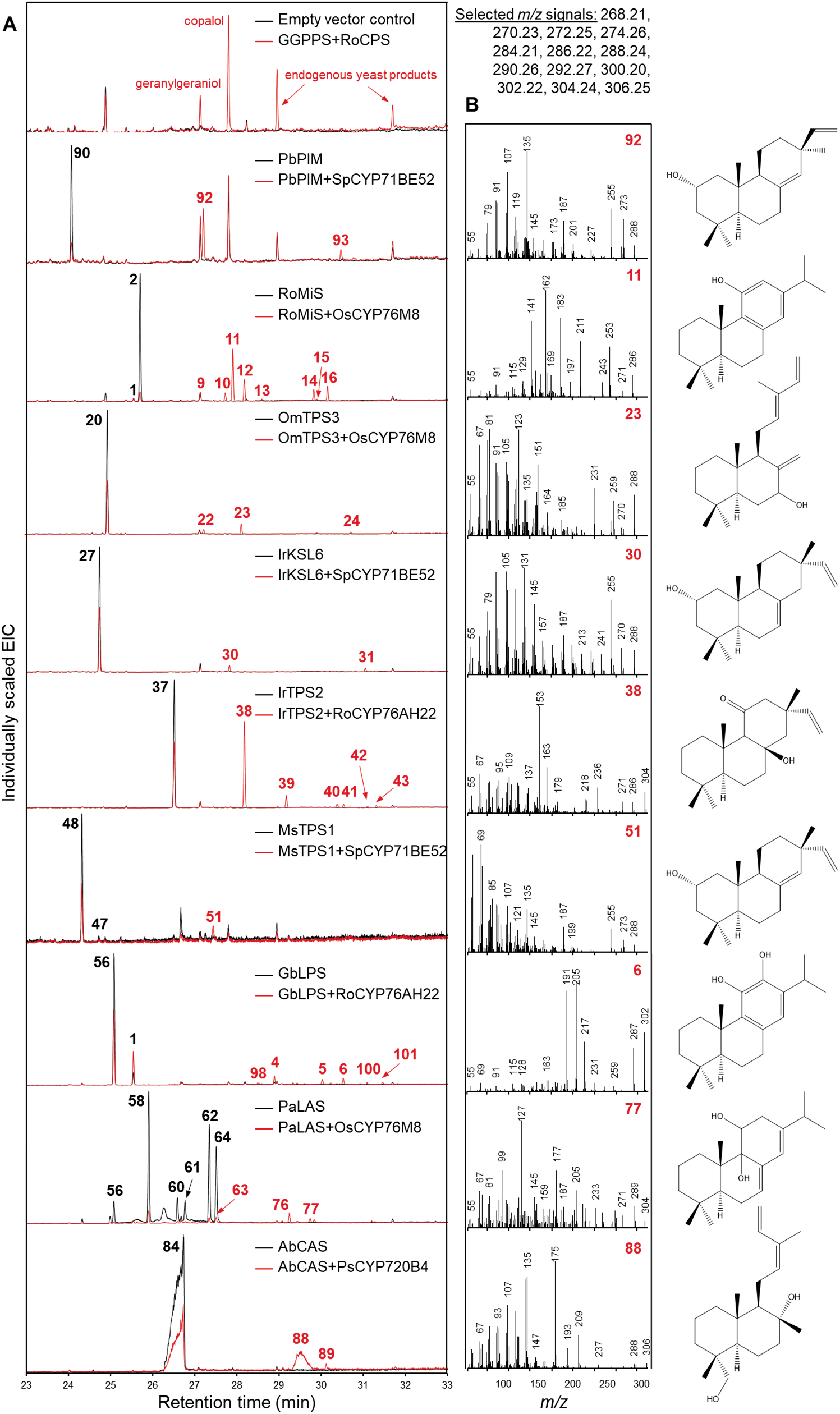
GC-MS analysis of combinatorial diterpene biosynthesis in yeast. A, Electron ionization chromatograms of selected yeast strains expressing the core module and the given enzymes. The *m/z* signals indicated were selected for the detection of diterpenoids. Diterpenoids that are produced in the control strain (black chromatogram) and in the yeast strain co-expressing a CYP (red chromatogram) are given as black numbers. Oxidized diterpenoids that derive from activity of the given CYP are indicated by red numbers. GGPP and CPP were detected as their dephosphorylated derivatives. B, Mass spectra of selected oxidized diterpenes with corresponding chemical structures.

**Figure 3.**
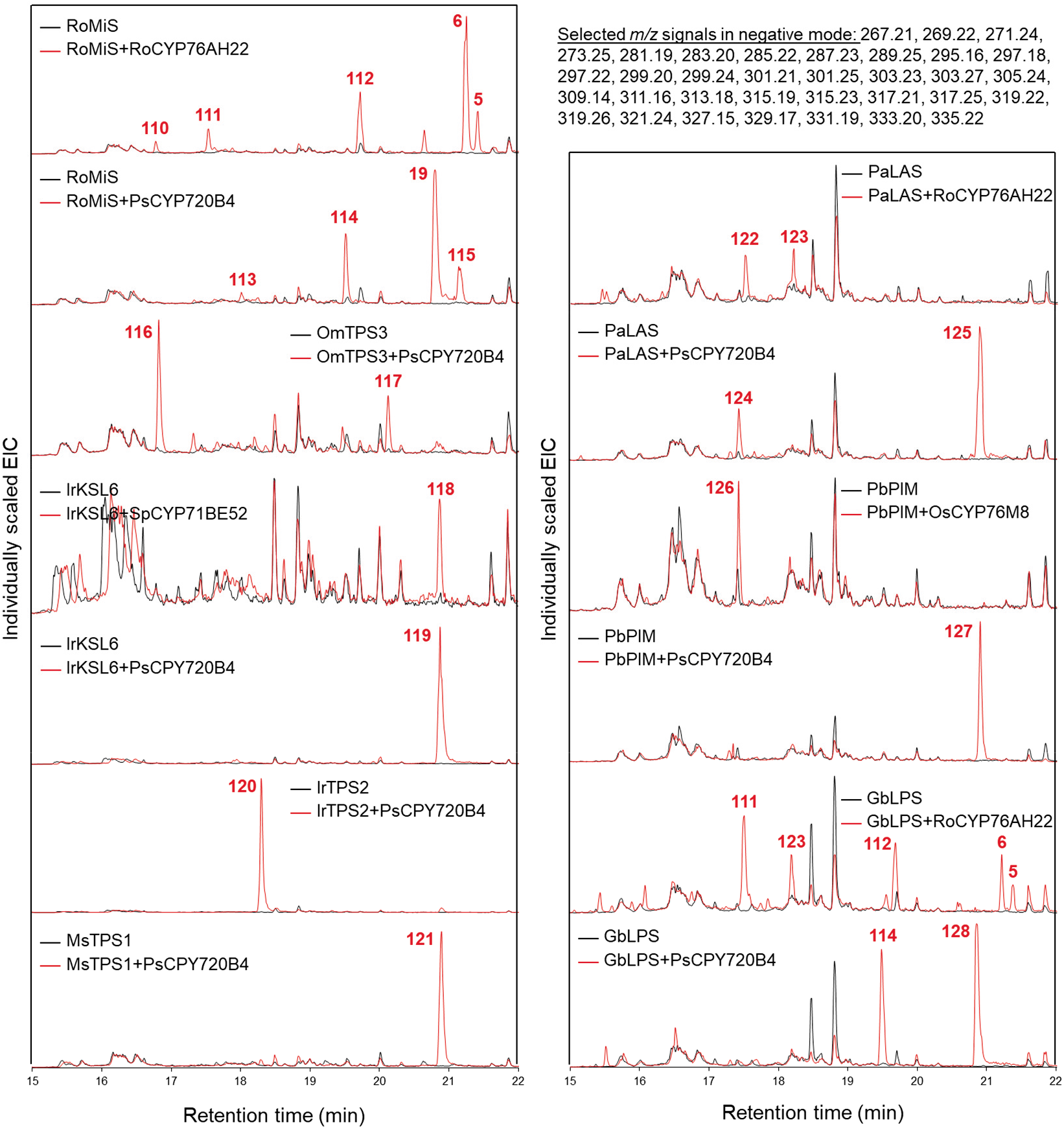
LC-MS analysis of yeast strains expressing the core module and the given enzymes. Control strains expressing no CYP are given in black. Oxidized diterpenoids that derive from CYP activity are given in red.

### Diterpene structure determination and suggestion

Mass errors of lower than 10 ppm and 5 ppm in GC-MS and LC-MS, respectively, allowed us to determine the molecular masses of the formed products with high confidence (Table S3 and Table S4). SpCYP71BE52 and PsCYP720B4 mostly introduced single hydroxyl and carboxyl groups, respectively, whereas RoCYP76AH22 and OsCYP76M8 catalyzed the formation of single or multiple hydroxyl and carbonyl groups. We used NMR analysis and comparison with databases and the literature to identify the structure of 40 of the formed diterpenoids (Table S3, Fig. S1-S12, Table S5-S8). For comparison with the literature, we mined SciFinder^n[14]^ to search for oxidized diterpene products reported before that have the same molecular mass as the compounds produced in this study. We assumed that the CYPs are regioselective not only with their natural substrate but also with slightly different diterpene backbones. We then introduced all possible chemical structures into SciFinder^n^ that derive from a diterpene skeleton produced by the diTPS in an engineered yeast strain and that is further oxidized at regioselective positions by the expressed CYP. For example, when RoMiS was co-expressed with OsCYP76M8, we searched for all reasonable oxidized diterpenes that derive from miltiradiene and abietatriene, the known products of RoMiS. As OsCYP76M8 was reported to oxidize tricyclic diterpenes at positions C6, C7, C9 and C11, we assumed theoretical miltiradiene and abietatriene derivatives oxidized at C6, C7, C9 and C11 that match the product masses determined by GC-MS or LC-MS. When a putative diterpene of *m/z* 286.22 was measured in GC-MS analysis of this yeast strain, we inserted 6-hydroxy-abietatriene, 7-hydroxy- abietatriene and 11-hydroxy- abietatriene, but also carbonyl derivatives of miltiradiene at C6, C7 and C11 into SciFinder^n^. The results were then mined for mass spectra and compared to our data.

In addition to NMR and comparison with the literature, we propose structures for 20 of the oxidized diterpenoids based on the fragmentation pattern of silylated or native extracts measured by GC-MS or LC-MS (Fig. 2, Fig. S13-37). These data confirm that the regioselectivity of the expressed CYPs on the diverse substrates they were presented with, is largely conserved. Thus, RoCYP76AH22 oxidizes positions C7, C11 and C12, SpCYP71BE52 position C2 and rarely C19, OsCYP76M8 positions C6, C7, C9 and C11 and PsCYP720B4 position C18 (Fig. 1C). In addition to oxygenation reactions, OsCYP76M8 also formed diterpenoids with an extra C-C bond, resulting in a double bond. This was observed with miltiradiene (**2**), abietatriene (**1**), levopimaradiene (**56**), *trans*-biformene (**20**) and nezukol (**37**) (Table S3). Although in the vast majority of cases no modifications other than oxidations were observed, in a few cases methylation (compound **112**) or acetylation (compounds **78**-**82**, **97**) occurred. These rarely occurring modifications are likely to be carried out by non-specific endogenous yeast enzymes. An overview of the network of products derived from each diterpene synthase is shown in Fig. S37A-H. In summary, for 80 of the 130 produced diterpenoids no information in the literature could be found, so that these compounds can be considered “new-to Nature”.

## Conclusion

Our results show that yeast is well suited for the combinatorial biosynthesis of diterpenoids. Recently, a combination of enzymatic oxidation and chemical synthesis allowed the synthesis of nine highly oxidized natural diterpenoids^[15]^. In that case, the starting diterpenoids were oxidized diterpenoids such as steviol. In contrast, our strategy does not depend on the availability of a specific diterpenoid but starts *ab initio* from known diTPS and advances further with known CYPs. All diterpene backbones produced have in common the same structure of the A-ring, because they are all derived from (*+*)-CPP, whereas the rest of structures around the B-ring is variable. Despite this common structural feature, not all substrates could be oxidized by PsCYP720B4 or SpCYP71BE52, two CYPs that specifically oxidize around the A-ring. This concerns mostly diterpenes without a C-ring and with a side chain instead, such as *trans*- biformene and *cis*-abienol (not oxidized by SpCYP71BE52), labda-7,13-*E*-dien-15-ol (oxidized by neither SpCYP71BE52 nor PsCYP720B4). This shows that although the CYPs conserve their regioselectivity, structural features that are further away from the oxidation sites can determine the ability of these CYPs to oxidize diterpene backbones. By selecting CYPs from distantly related species but that are known to act on diterpene backbones derived from (*+*)-CPP we could generate a high proportion of compounds that have not yet been reported. Our approach opens up further possibilities, either by further combining CYPs on (*+*)-CPP derived diterpenes or by applying the same strategy to other groups of diterpene skeletons, e.g. derived from other CPP isoforms, or macrocyclic ones, to generate diverse libraries of diterpenoids in high throughput using yeast.

## Supporting information

All Materials and Methods, supplemental Figures and Tables are available in the Supplemental Information file, except Table S3 and Table S4, which are available as separate files.

## Acknowledgements

The authors thank Dr. Andrea Porzel for assisting NMR analysis and Chris Richter for technical support regarding measurements at the GC-QTOF. The work was supported by the European Regional Development Fund (grant number ZS/2018/08/94166 to AT) and the Leibniz Institute of Plant Biochemistry in Halle, Germany.

## Materials and Methods

### Gene synthesis and cloning

Synthetic genes (PbPIM, RoMiS, OmTPS3, IrKSL6, IrTPS2, MsTPS1, GbLPS, PaLAS, SmCPSKLS1, AbCAS, SpCYP71BE52, OsCYP76M8 and PsCYP720B4) were synthesized by GeneArt (Thermo Fisher Scientific) as codon optimized sequences for expression in *Saccharomyces cerevisiae* (Table S1). Sequences of NtGGPPS, RoCPS, ATR1 and RoCYP76AH22 were used as described before ^[4d]^. Transit peptides of PbPIM, RoMiS and GbLPS were predicted using ChloroP (http://www.cbs.dtu.dk/services/ChloroP/) and deleted using the primers listed in Table S9.

Golden Gate compatible yeast expression vectors were created as described before ^[4d]^. Expression units consisting of synthetic galactose inducible promoters, genes (Table S1) and terminators were assembled in Golden Gate vectors of level 1. Expression units as given in Table S3 were finally assembled in yeast Golden Gate expression vectors of level M.

### Production of diterpenoids in yeast and extraction

Yeast expression vectors (Table S3) were transformed into yeast strain INVSc1 (Invitrogen) and plated out onto uracil-free selection medium (1 g/L Yeast Synthetic Drop-out Medium Supplements without uracil, 6.7 g/L Yeast Nitrogen Base Without Amino Acids, 20 g/L Micro Agar). Positively transformed colonies were inoculated into 5 mL YPD medium (20 g/L tryptone and 10 g/L yeast extract) containing 2 % of glucose and grown for 24 h with shaking at 30 °C. Protein expression was induced by resuspending the cell pellet in fresh YPD medium containing 2 % of galactose. After another 24 h of cultivation, whole cultures were extracted using 2 mL *n*-hexane.

Large volume production of diterpenoids for NMR analysis were started with 40 precultures per construct. Each preculture of 5 mL uracil-free selection medium (1 g/L Yeast Synthetic Drop-out Medium Supplements without uracil, 6.7 g/L Yeast Nitrogen Base Without Amino Acids) containing 2 % of glucose were inoculated with a single positively transformed colony and grown for 24 h with shaking at 30 °C. Cultures were centrifuged the next day and the cells were resuspended in fresh uracil-free selection medium containing 2 % of glucose. After cultivation for 24 h at 30 °C while shaking, cells of 20 precultures were transferred into 1.5 L uracil-free selection medium containing 2 % of glucose in a 5 L-flask. Cultivation was performed at 30 °C for 24 h with moderate shaking (100 rpm). Protein expression was induced by resuspending the cell pellet in fresh uracil-free selection medium containing 2 % of galactose. After another 24 h of cultivation, cells were resuspended in fresh uracil-free selection medium containing 2 % of galactose and the supernatant was kept for further extraction. This was repeated the next day. Supernatants from 2 days of expression and the final culture were extracted twice, each time with 4 L ethyl acetate or hexane. The pooled extract was dried using a rotary evaporator.

### Diterpenoid purification for NMR analysis

Dried raw yeast extracts from large-volume production contained either **38** or **120**. The extracts were dissolved in 76% acetonitrile/water and purified on a HPLC system which consisted of two PU-2080 Plus pumps, a DG-2080−53 degasser, a MD-2010 Plus type diode array detector operating at 220 nm (Jasco, Gross-Umstadt, Germany), and a Rh 7725i type Rheodyne injection valve (Rheodyne, Bensheim, Germany). Isolation of all compounds was done using a semi-preparative RP-18 column (250 × 10.0 mm, 5 μm, Hyperclone ODS C18 column, Phenomenex, Aschaffenburg, Germany) operating with a flow rate of 4.8 mL/min. For elution, aqueous 0.1 % formic acid (A) and 0.1 % formic acid in acetonitrile (B) were used. The elution conditions for **38** (injection volume 300 µL) were as follows: isocratic from 0 to 5 min at 65 % eluent B, from 5 to 30 min linear from 65 to 100 % eluent B, isocratic from 30 to 32 min at 100 % eluent B, from 32 to 35 min from 100 to 65 % eluent B and isocratic from 35 to 40 min at 65 % eluent B. **120** (injection volume 250 µL was separated using the following gradient: isocratic from 0 to 5 min at 50 % eluent B, from 5 to 30 min linear from 65 to 100 % eluent B, isocratic from 30 to 32 min at 100 % eluent B, from 32 to 35 min from 100 to 50 % eluent B and isocratic from 35 to 40 min at 50 % eluent B. Data acquisition was performed using Chrompass or Galaxy software (Jasco, Gross-Umstadt, Germany). 2.97 mg of **38** (purity > 97%) and 5.81 mg of **120** (purity > 95 %) were used for NMR analysis.

### Derivatization

100 µL pyridine and 100 µL MSTFA (Macherey-Nagel) were added to dried yeast extracts and incubated for 2 h at 70 °C.

### GC-MS analysis

Extracts of yeast strain grown in YPD medium were dried under a nitrogen stream and resuspended in 200 µl of *n*-hexane. GC-MS analysis carried out using an Agilent 7890B gas chromatograph coupled to an Agilent 7200 Accurate-Mass Quadrupole Time-of-Flight mass spectrometer with electron impact ionization. Chromatographic separation was performed on a HP-5MS UI capillary column (30 m, 0.25 mm, 0.25 µm, Agilent) using split or splitless injection with a liner temperature of 250 °C and an injection volume of 1 µl. The flow rate of helium was 1 ml/min and the GC oven temperature ramp was as follows: 50 °C for 1 min; 50 to 290 °C with 7 °C/min. Mass spectrometry was performed at 70 eV with a source temperature of 230 °C. A full scan mode with *m/z* from 50 to 450 and an acquisition rate of 6.5 spectra/sec was performed in a 4 GHz high resolution mode. Mass calibration was done before each measurement. Data analysis was carried out using the device specific software Qualitative Navigator B.08.00.

Derivatized diterpenes were analyzed using a Trace GC Ultra gas chromatograph (Thermo Scientific) coupled to an ATAS Optic 3 injector and an ISQ single quadrupole mass spectrometer (Thermo Scientific) with electron impact ionization. The compounds were separated on a ZB-5ms capillary column (30 m x 0.32 mm, Phenomenex) using splitless injection and an injection volume of 1 µL. An injection temperature gradient from 60 °C to 250 °C with 10 °C/s was used and the flow rate of helium was 1 mL/min. The GC oven temperature gradient was as follows: 50 °C for 1 min, 50 to 300 °C with 7 °C/min, 300 to 330 °C with 20 °C/min and 330 °C for 5 min. Mass spectrometry was performed at 70 eV, in a full scan mode with *m/z* from 50 to 500. Data analysis was done with the device specific software Xcalibur (Thermo Scientific).

### LC-MS^n^ experiments

The methanolic extracts (50 µL) were transferred to 2-mL vials with glass inserts, and 5 µL of each sample were loaded on a EC 150/2 Nucleoshell RP18 column (endcaped C18 phase, ID 2 mm, length 150 mm, particle size 2.7 μm, Macherey Nagel, Düren, Germany) using a Dionex Ultimate 3000 UHPLC, equipped with a 3400RS pump and 3000TRS autosampler (Thermo-Fisher Scientific, Bremen, Germany). The eluents A and B were 0.3 mmol/L ammonium formate (adjusted to pH 3.5 with formic acid) and acetonitrile, respectively. After a 2-min isocratic step (5% eluent B), analytes were eluted at a flow rate of 400 µL/min at 40° C in a 17-min linear gradient to 95% eluent B or to 45% eluent B. The column effluents were introduced on-line in an Orbitrap Elite mass spectrometer operated in negative ion mode, equipped with a heated electrospray (HESI) ion source, and controlled by Xcalibur (Thermo- Fisher Scientific, Bremen, Germany). The source and transfer capillary temperatures were set to 300°C. The spray voltage was -3.8 kV, while sheath and auxiliary gases were set to 18 and 5 psig, respectively. Analytes were annotated in preliminary data-dependent acquisition experiments designed according to the doubly play algorithm. The DDA experiments comprised a survey FT-scan with mass resolution of 30000 (*m/z* 100 – 1500) followed with dependent linear ion trap (LIT) scans for the two most abundant signals selected in each survey scan. Analytes were annotated in survey scans by their tR, m/z and isotopic patterns.

The structures of the annotated analytes were confirmed by tandem mass spectrometric (MSn) analysis. Thereby, collision induced fragmentation (CID) was performed in LIT by resonance activation in presence of He as a collision/cooling gas. The corresponding quasi-molecular ions were isolated with the width of 2 *m/z*, activation time and relative activation frequency were 10 ms and 0.250, respectively. Normalized collision energy was experiment-specific and varied in the range of 30 – 45%. For some analytes higher-energy C- trap dissociation (HCD) was performed in C-trap. In those experiments, kinetic energy was experiment-specific and varied in the range of 160 – 200 eV.

### Structure elucidation by NMR and quantification

Samples were analyzed on a Bruker Avance Neo 600 MHz system (Bruker Rheinstetten, Germany), equipped with a Z-gradient 5 mm TCI Probe at 298 K. The samples were analyzed in 5 mm × 7’’ NMR tubes (Z107374 USC tubes, Bruker, Faellanden, Switzerland). Data processing was done using the Bruker TopSpin software (version 4.0.7). Zero and first order phase correction were performed manually, and baseline correction was performed automatically using the command *absn*. For all 1D and 2D spectra standard Bruker pulse sequences were used ^1^H (*zg30*), ^13^C (*zgpg30, deptsp135*), COSY (*cosygppqf*), HSQC (*hsqcedetgpsp.3*), HMBC (*hmbcetgpl3nd*). All data for acquisition and processing can be found in the original data as well as in the spectra.

For quantitative NMR analysis (qHNMR), the ERETIC 2 procedure was applied on a Bruker AV III 400 MHz system (Bruker Rheinstetten, Germany), equipped with a Z-gradient 5 mm multinuclear observe probe (BBFO_plus_) at 298 K, as described in the literature, using *L*- tyrosine (5.21 mmol/L) as the external standard for spectrometer calibration, integrating the specific resonance signal at 7.10 ppm (m, 2H)^[16]^. For compound quantitation, stock solutions of compounds **38** and **120** were prepared and aliquots (600 *µ*L) were analyzed by means of NMR. The concentration of each compound was calculated using the ERETIC 2 software tool of TopSpin (version 3.5, Bruker, Rheinstetten, Germany). The exact concentrations of all measured compounds, determined via qHNMR, were used for the preparation of stock solutions for experiments.

---

**Fig. S1.**
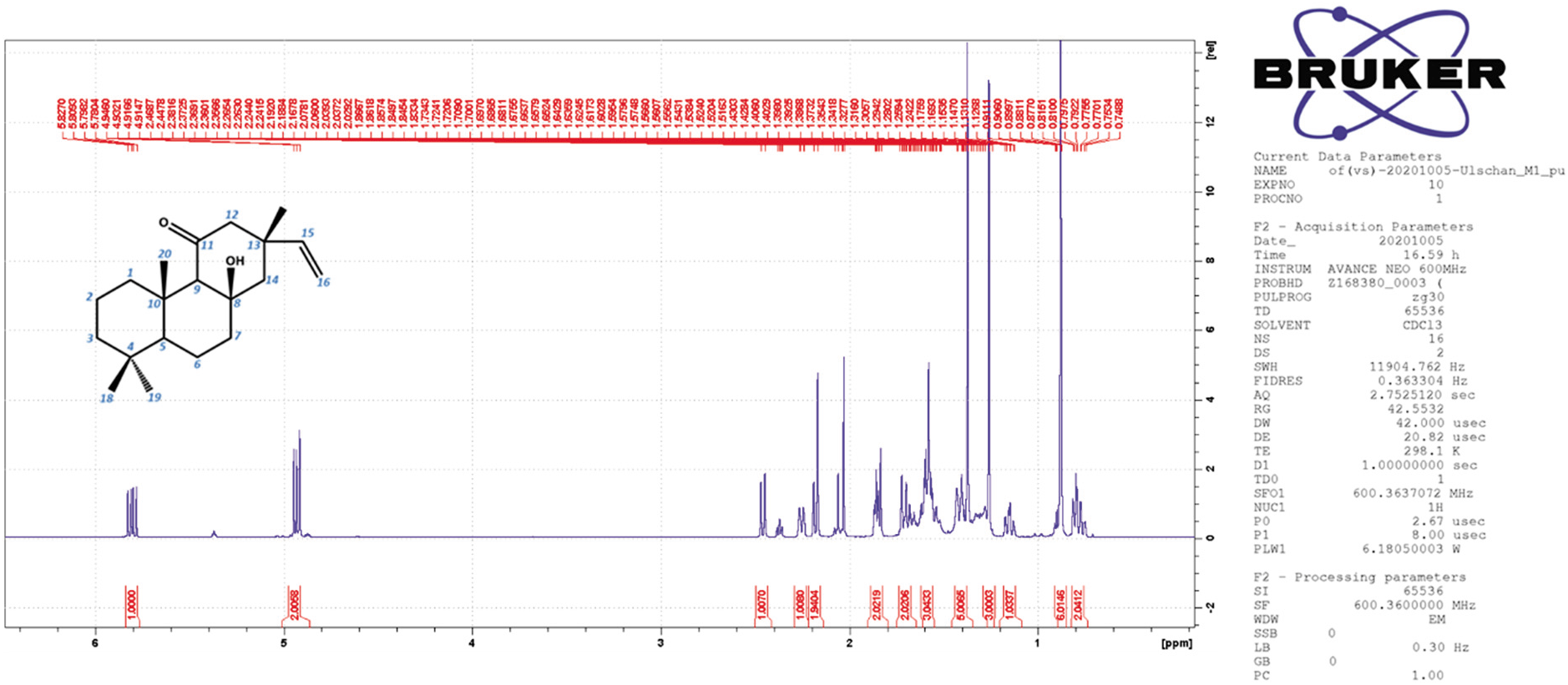
^1^H NMR spectrum (600 MHz, CDCl_3_, 298 K) of 38.

**Fig. S2.**
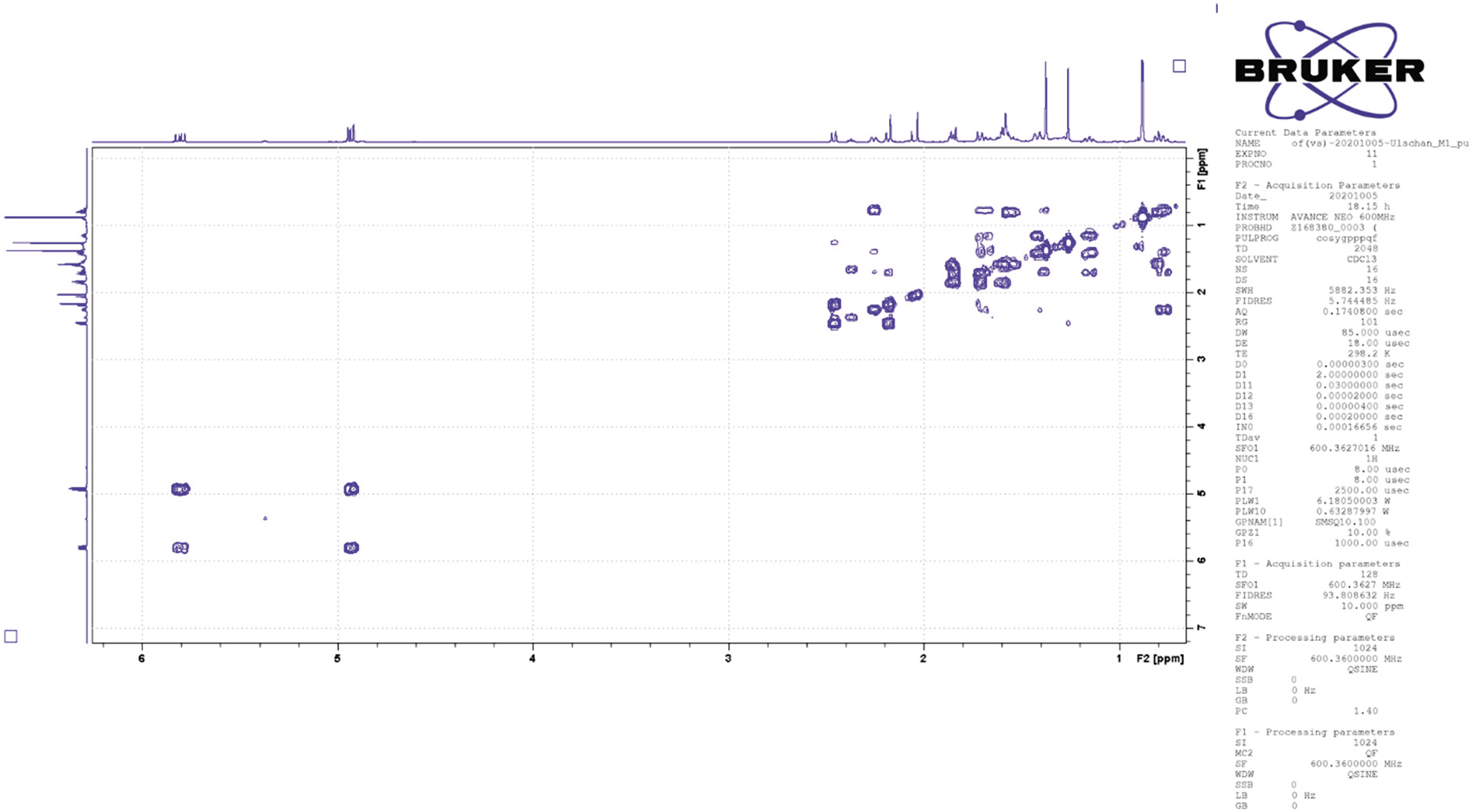
H,H COSY NMR spectrum (600 MHz, CDCl_3_, 298 K) of 38.

**Fig. S3.**
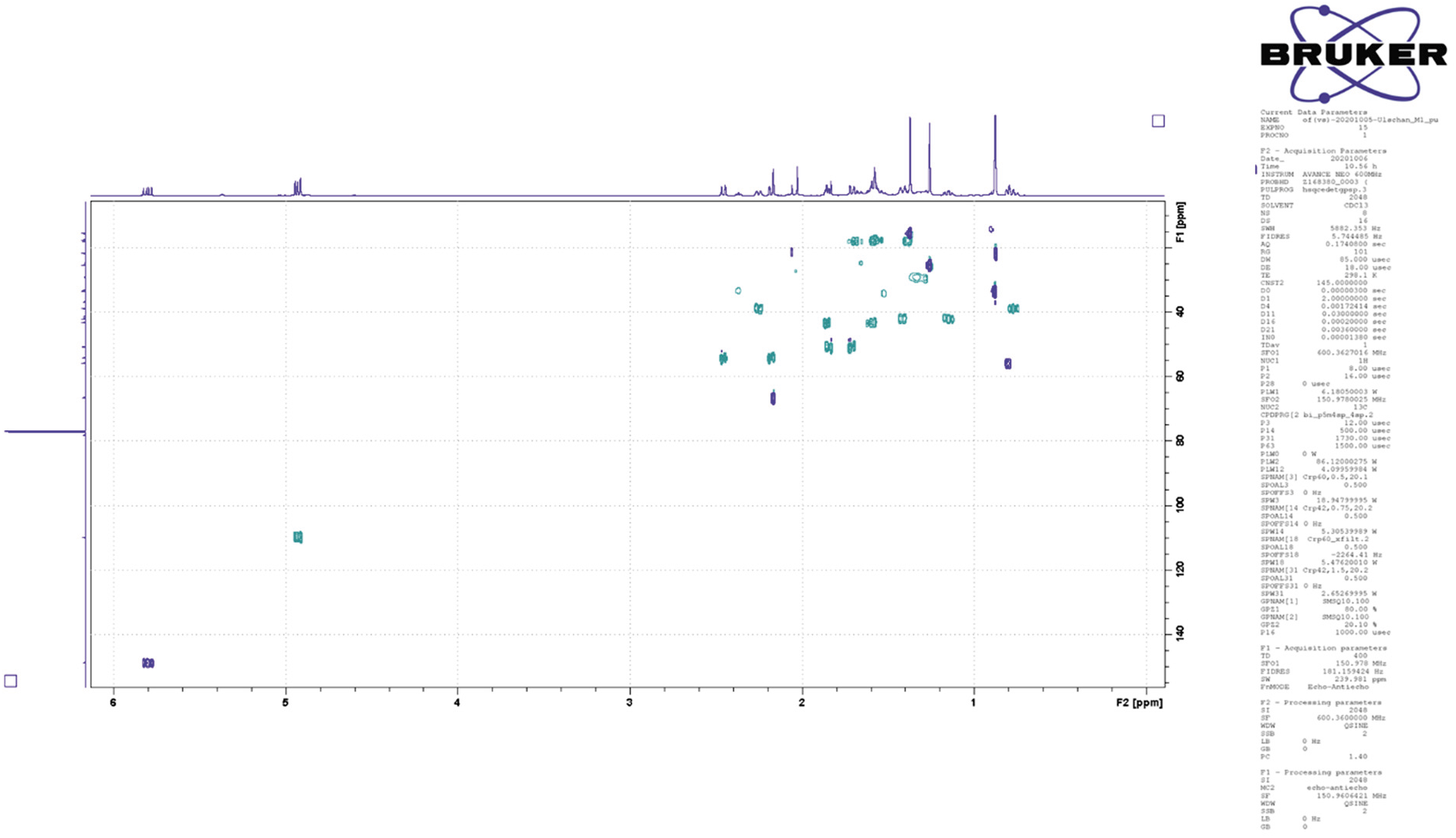
^1^H, ^13^C HSQC NMR spectrum (600 MHz, CDCl_3_, 298 K) of 38.

**Fig. S4.**
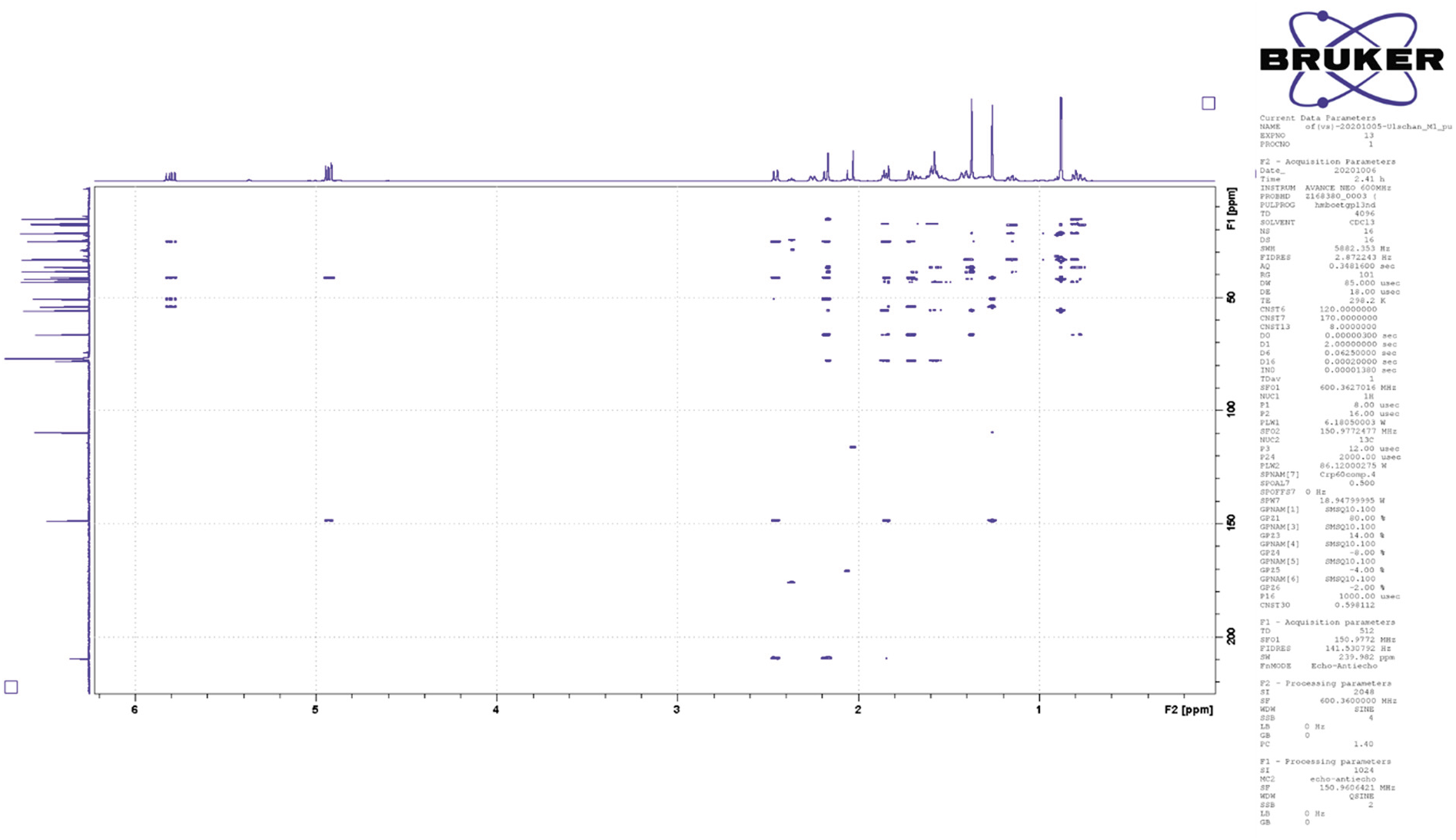
^1^H, ^13^C HMBC NMR spectrum (600 MHz, CDCl_3_, 298 K) of 38.

**Fig. S5.**
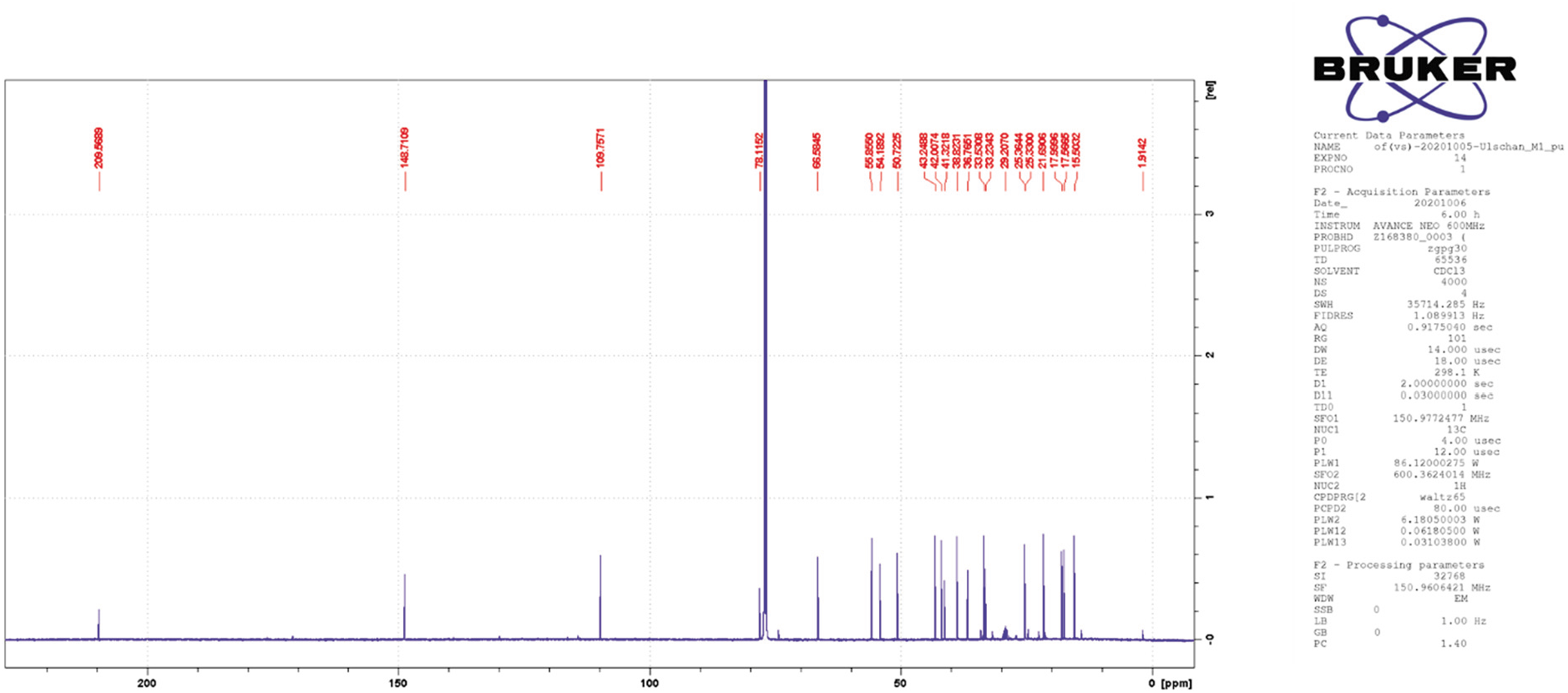
^13^C NMR spectrum (150 MHz, CDCl_3_, 298 K) of 38.

**Fig. S6.**
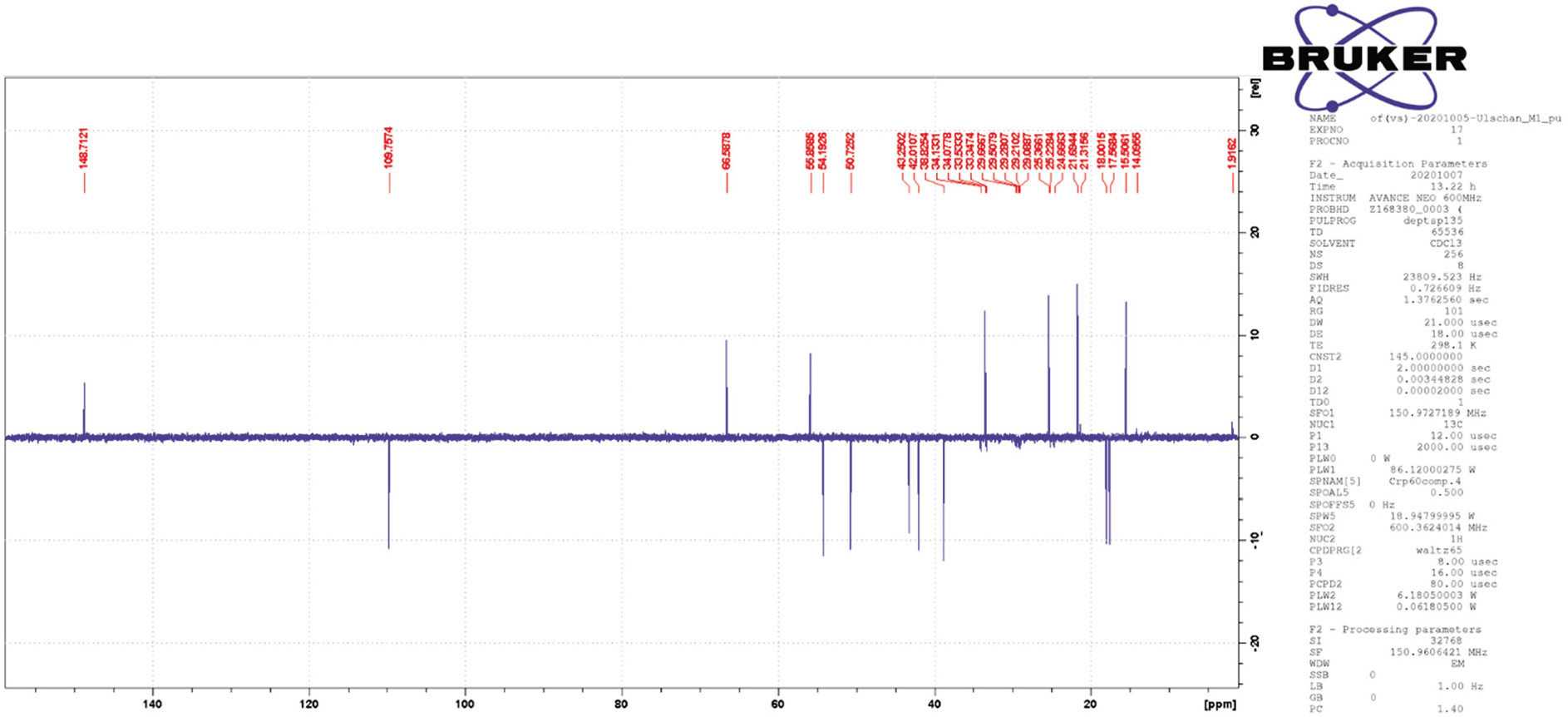
^13^C 135° DEPT NMR spectrum (150 MHz, CDCl_3_, 298 K) of 38.

**Fig. S7.**
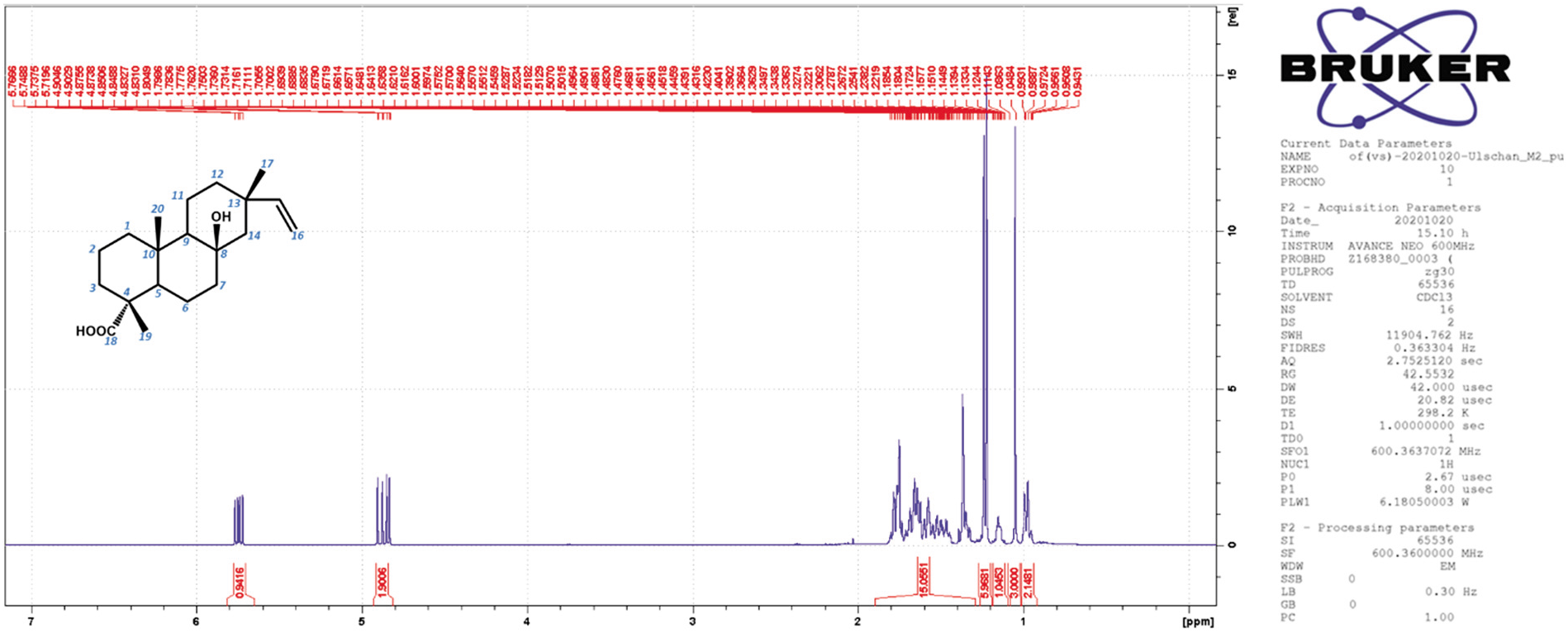
^1^H NMR spectrum (600 MHz, CDCl_3_, 298 K) of 120.

**Fig. S8.**
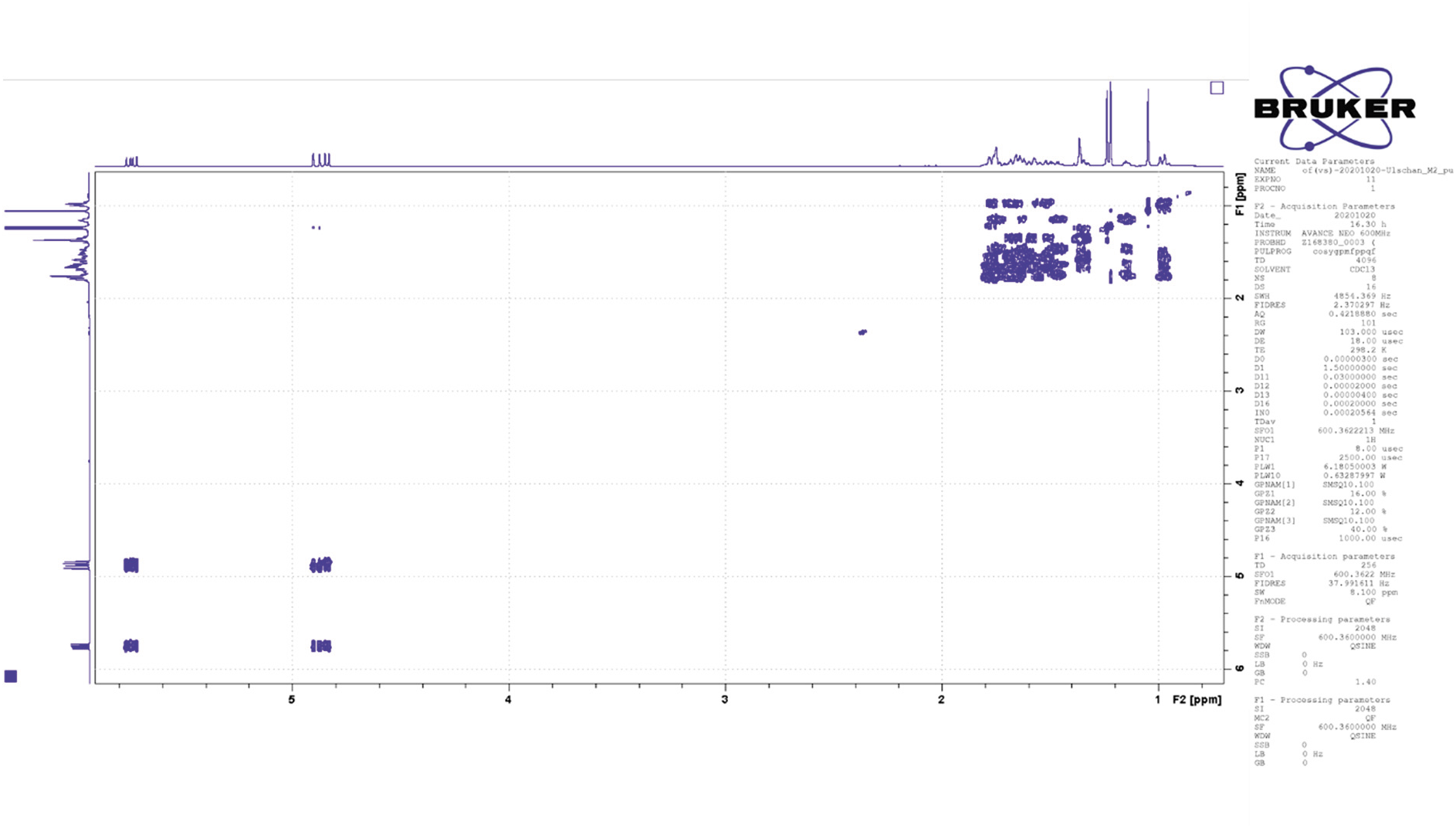
H,H COSY NMR spectrum (600 MHz, CDCl_3_, 298 K) of 120.

**Fig. S9.**
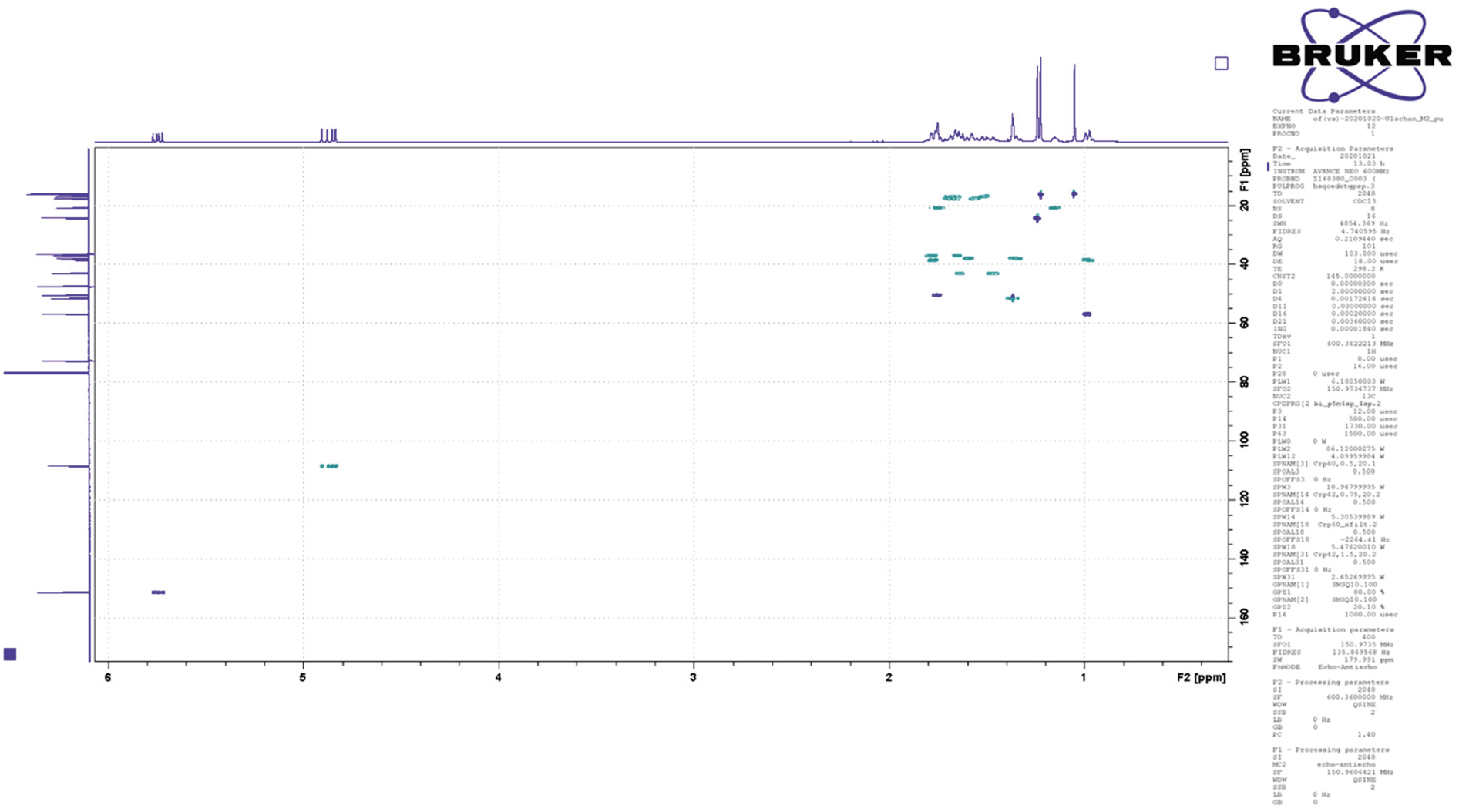
^1^H, ^13^C HSQC NMR spectrum (600/150 MHz, CDCl_3_, 298 K) of 120.

**Fig. S10.**
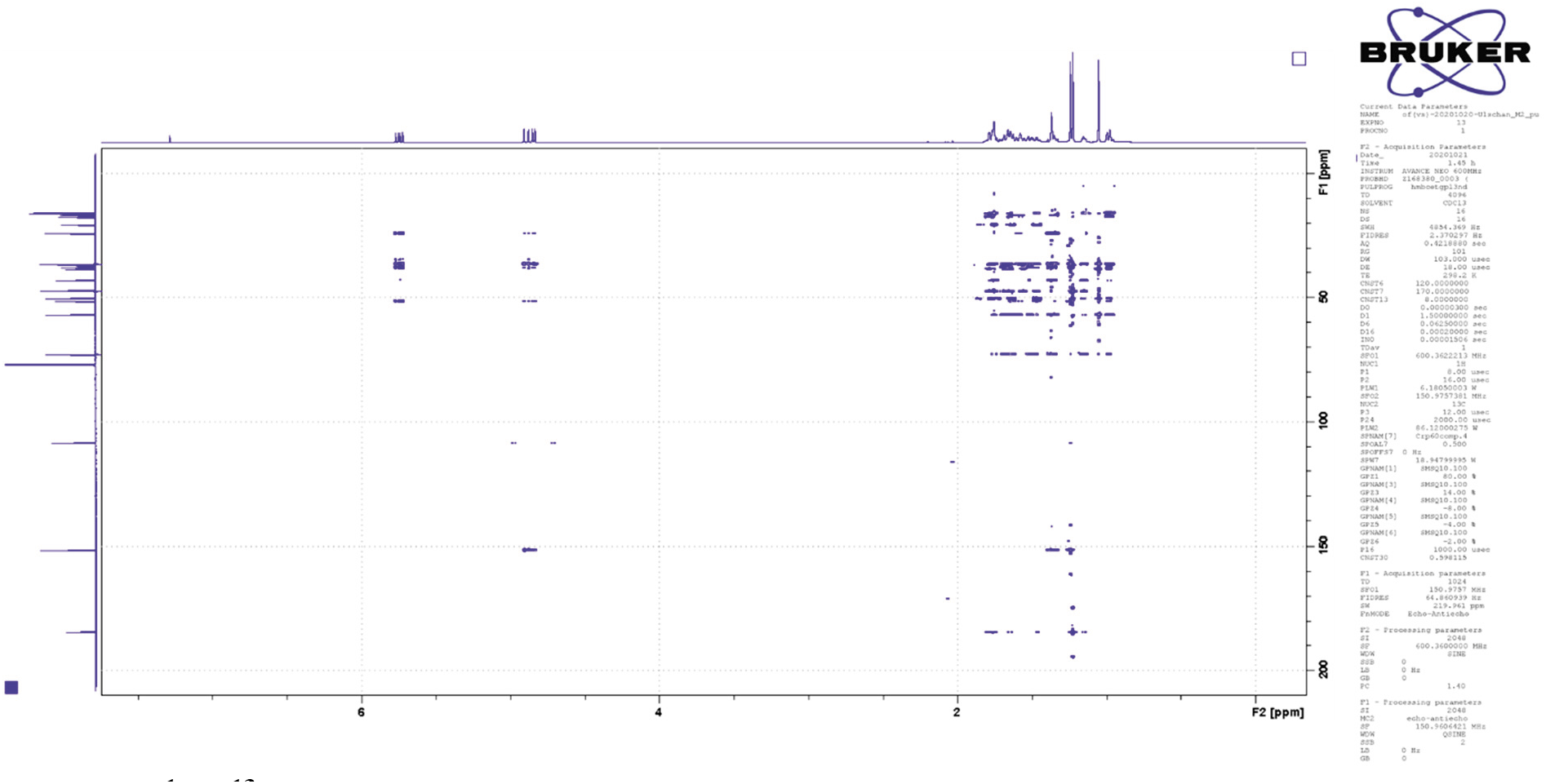
^1^H, ^13^C HMBC NMR spectrum (600/150 MHz, CDCl_3_, 298 K) of 120.

**Fig. S11.**
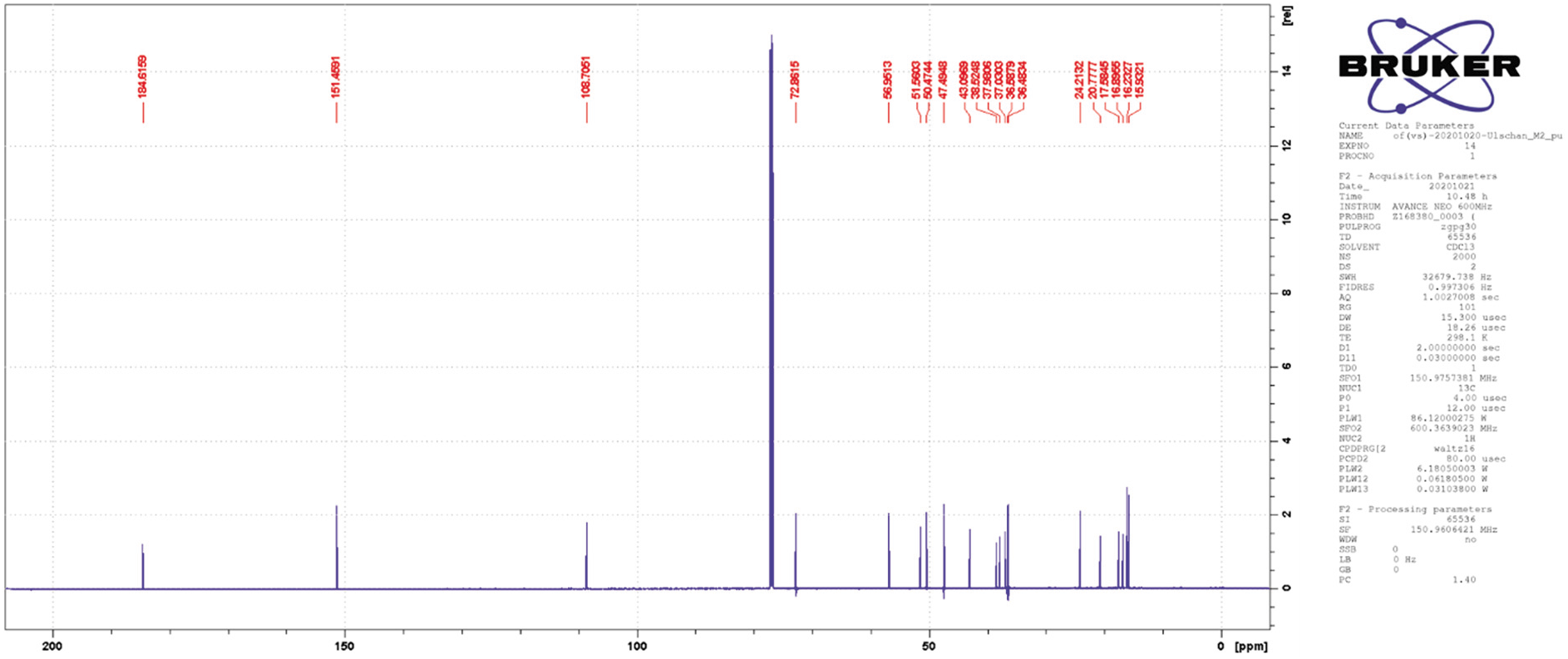
^13^C NMR spectrum (150 MHz, CDCl_3_, 298 K) of 120.

**Fig. S12.**
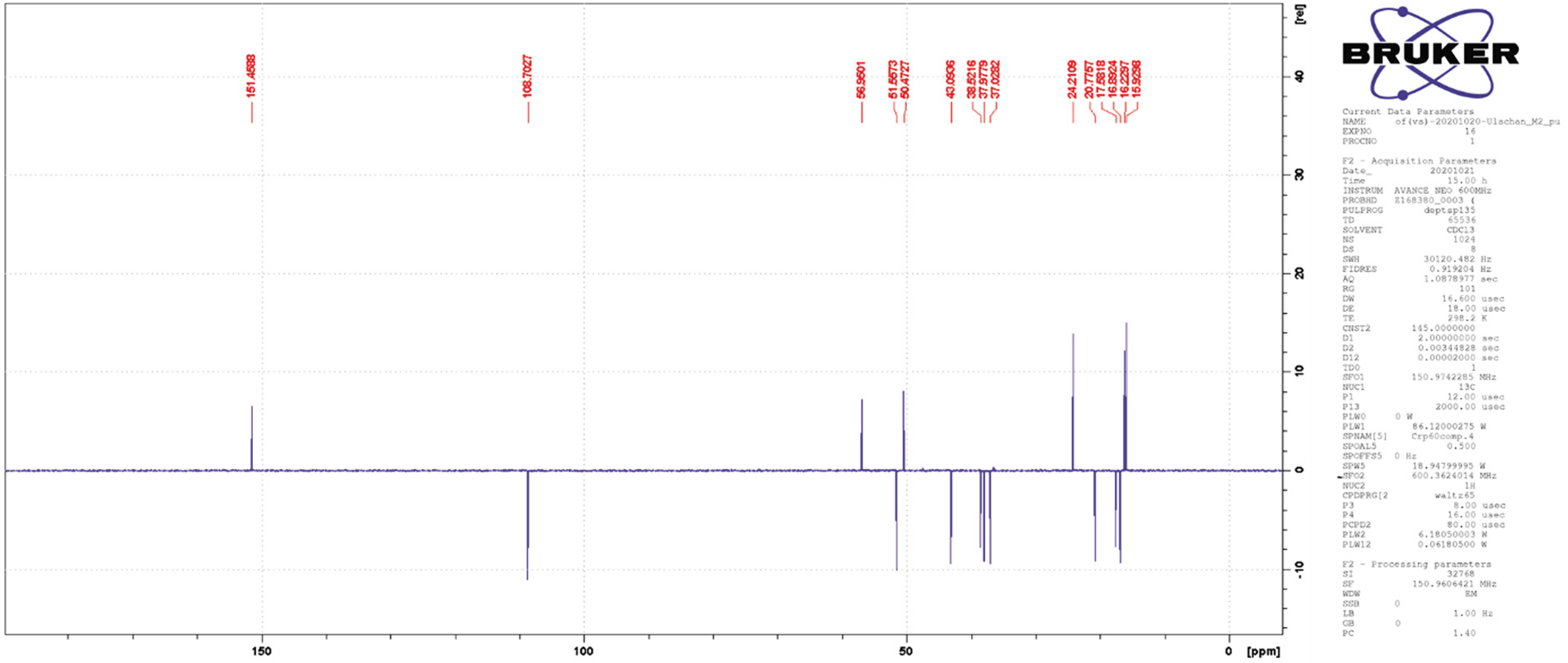
^13^C 135° DEPT NMR spectrum (150 MHz, CDCl_3_, 298 K) of 120.

**Fig. S13.**
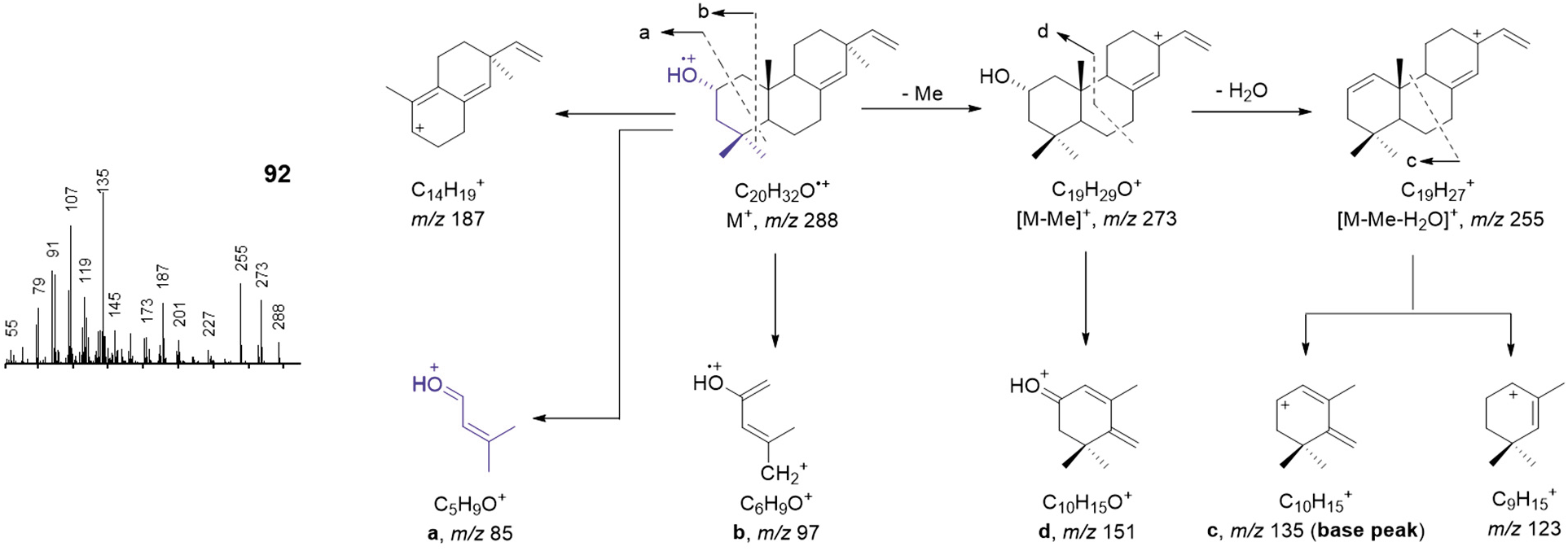
Proposed electron impact ionization (EI) mass spectral fragmentation of 92.

**Fig. S14.**
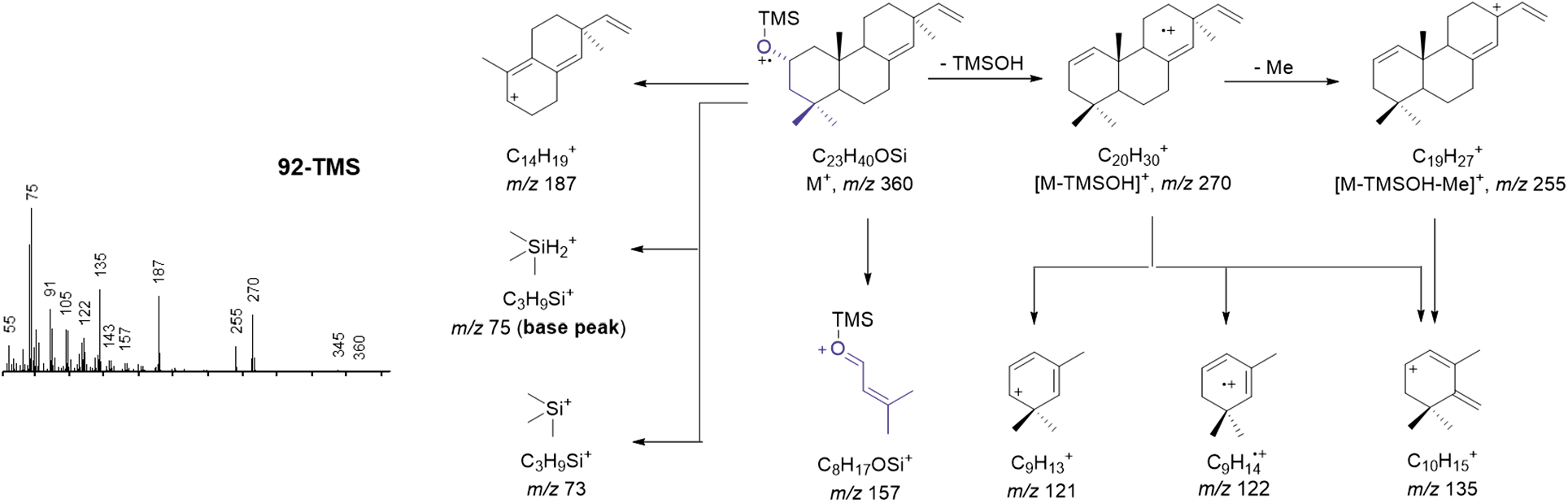
Proposed electron impact ionization (EI) mass spectral fragmentation of silylated 92.

**Fig. S15.**
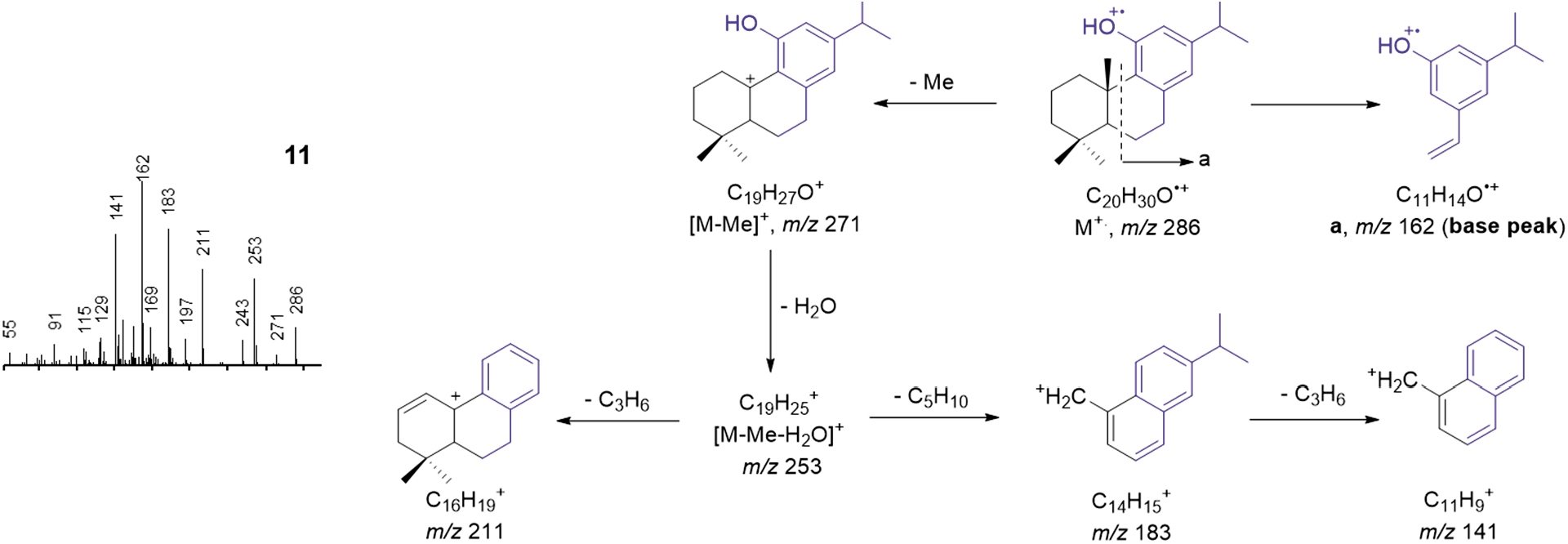
Proposed electron impact ionization (EI) mass spectral fragmentation of 11.

**Fig. S16.**
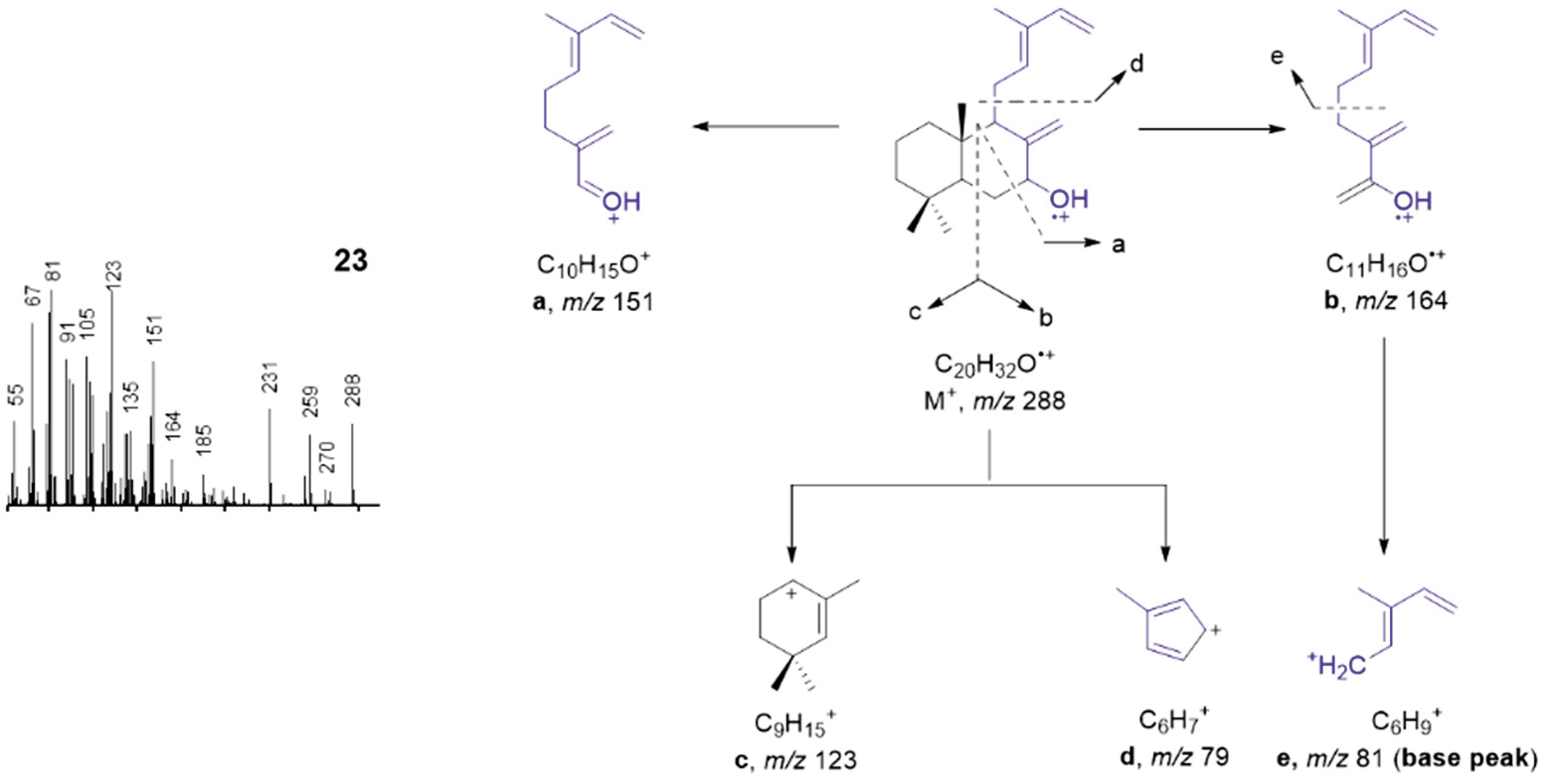
Proposed electron impact ionization (EI) mass spectral fragmentation of 23.

**Fig. S17.**
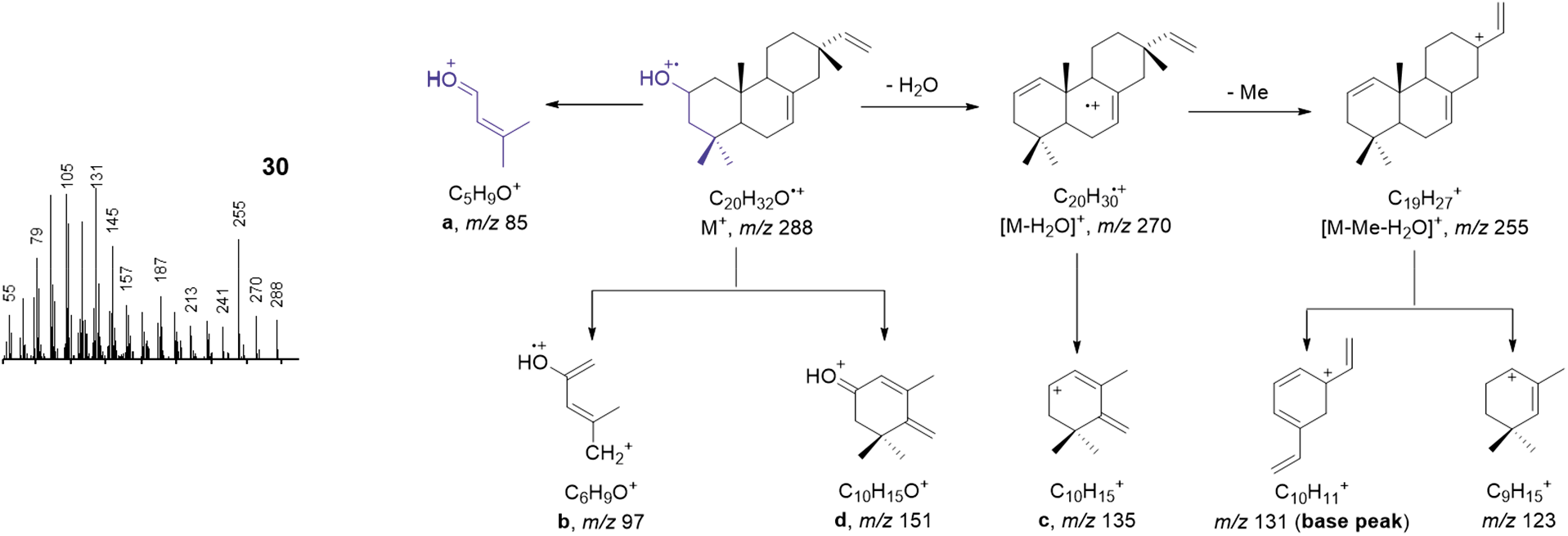
Proposed electron impact ionization (EI) mass spectral fragmentation of 30.

**Fig. S18.**
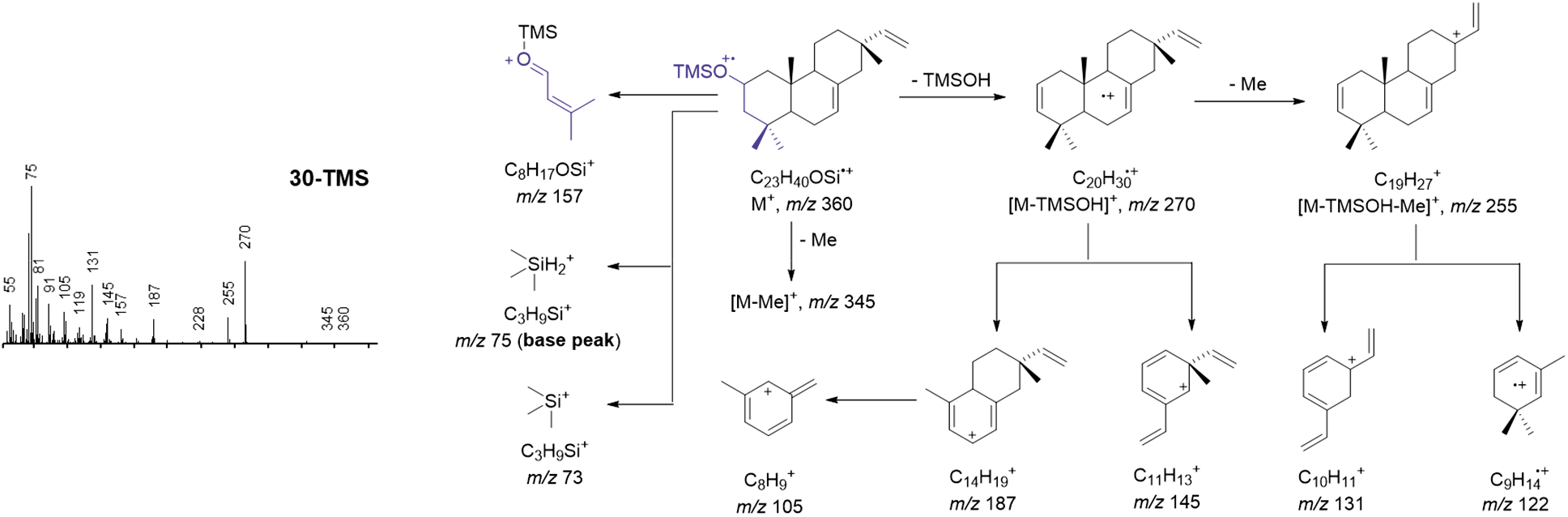
Proposed electron impact ionization (EI) mass spectral fragmentation of silylated 30.

**Fig. S19.**
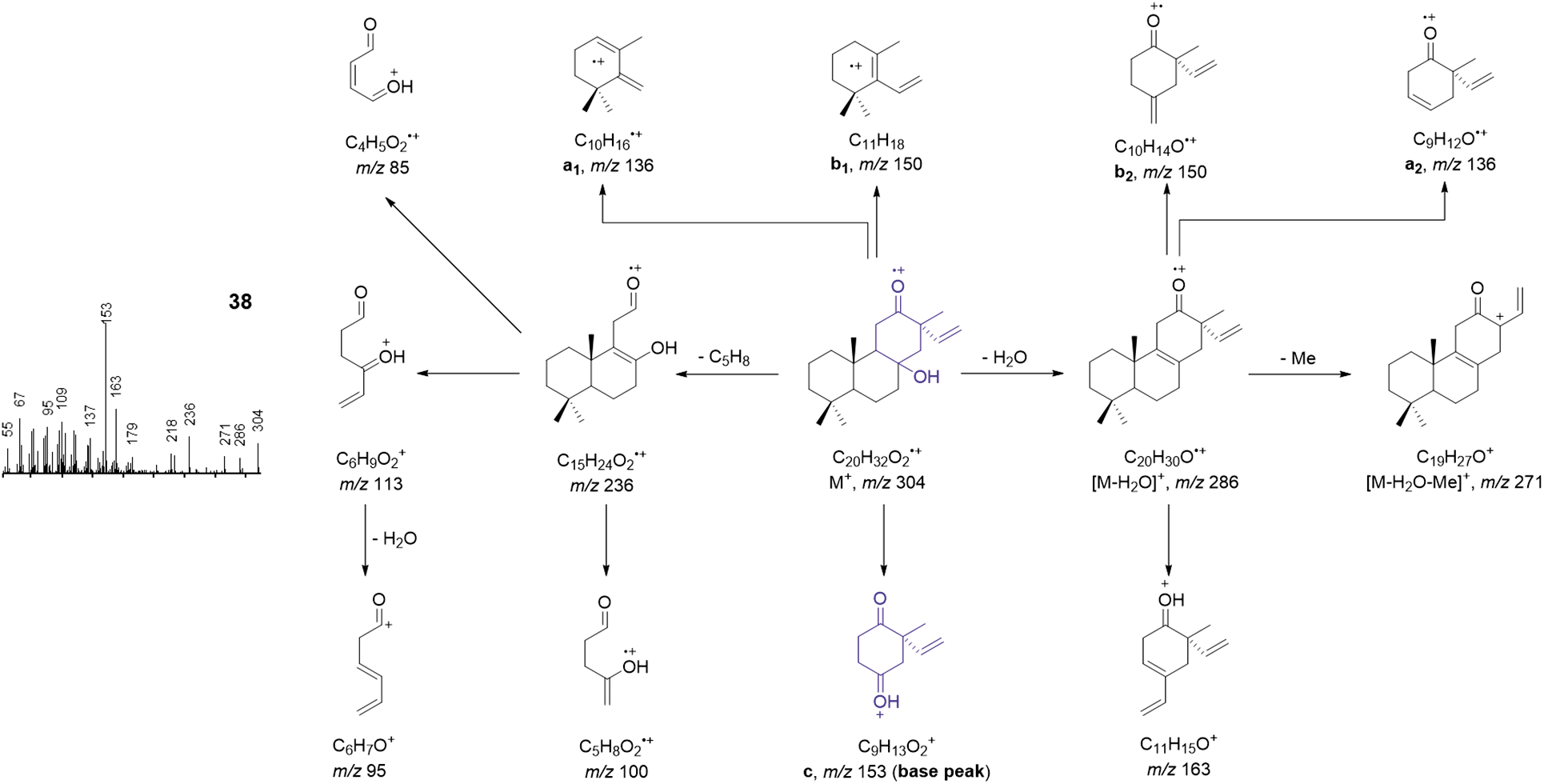
Proposed electron impact ionization (EI) mass spectral fragmentation of 38.

**Fig. S20.**
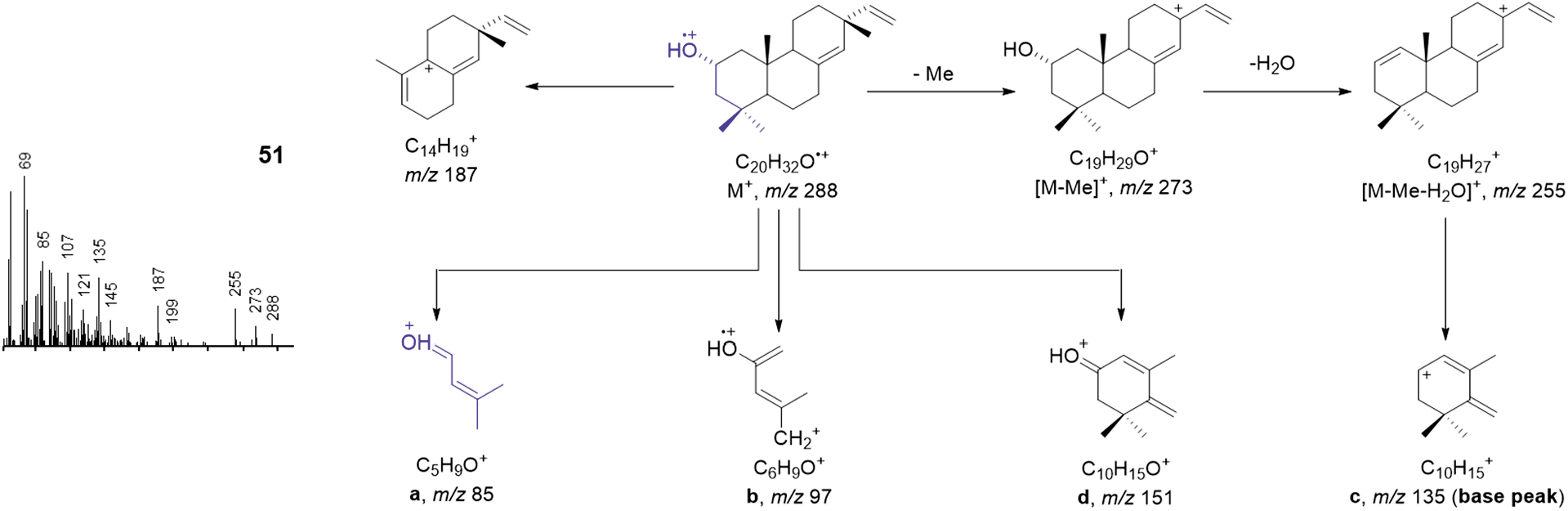
Proposed electron impact ionization (EI) mass spectral fragmentation of 51.

**Fig. S21.**
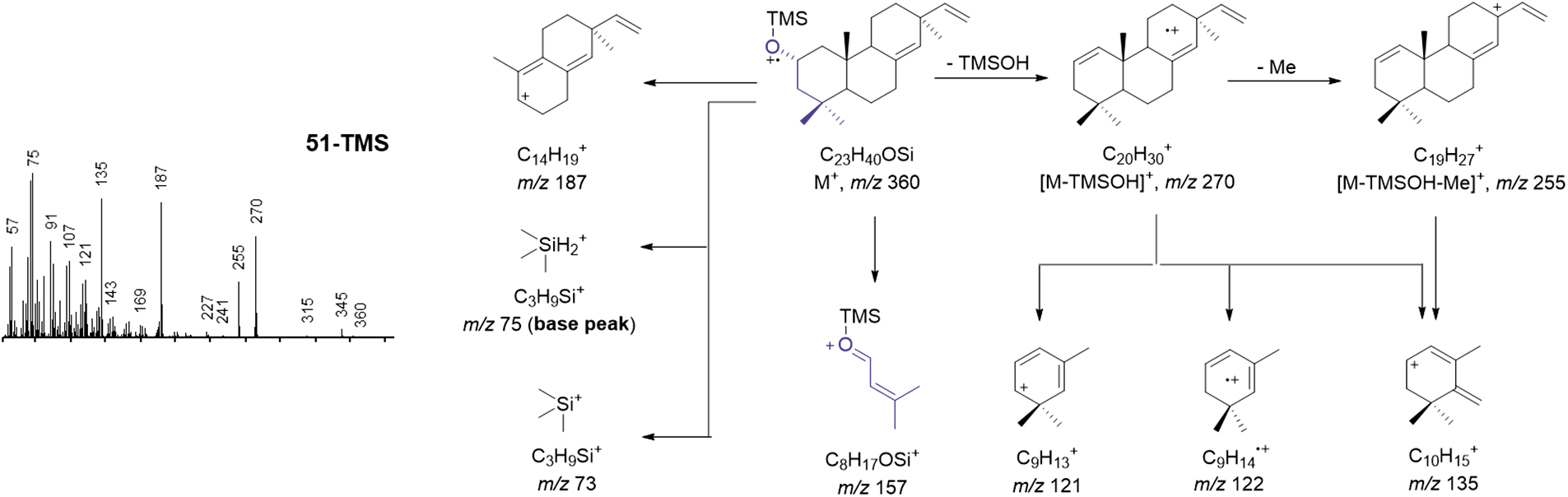
Proposed electron impact ionization (EI) mass spectral fragmentation of silylated 51.

**Fig. S22.**
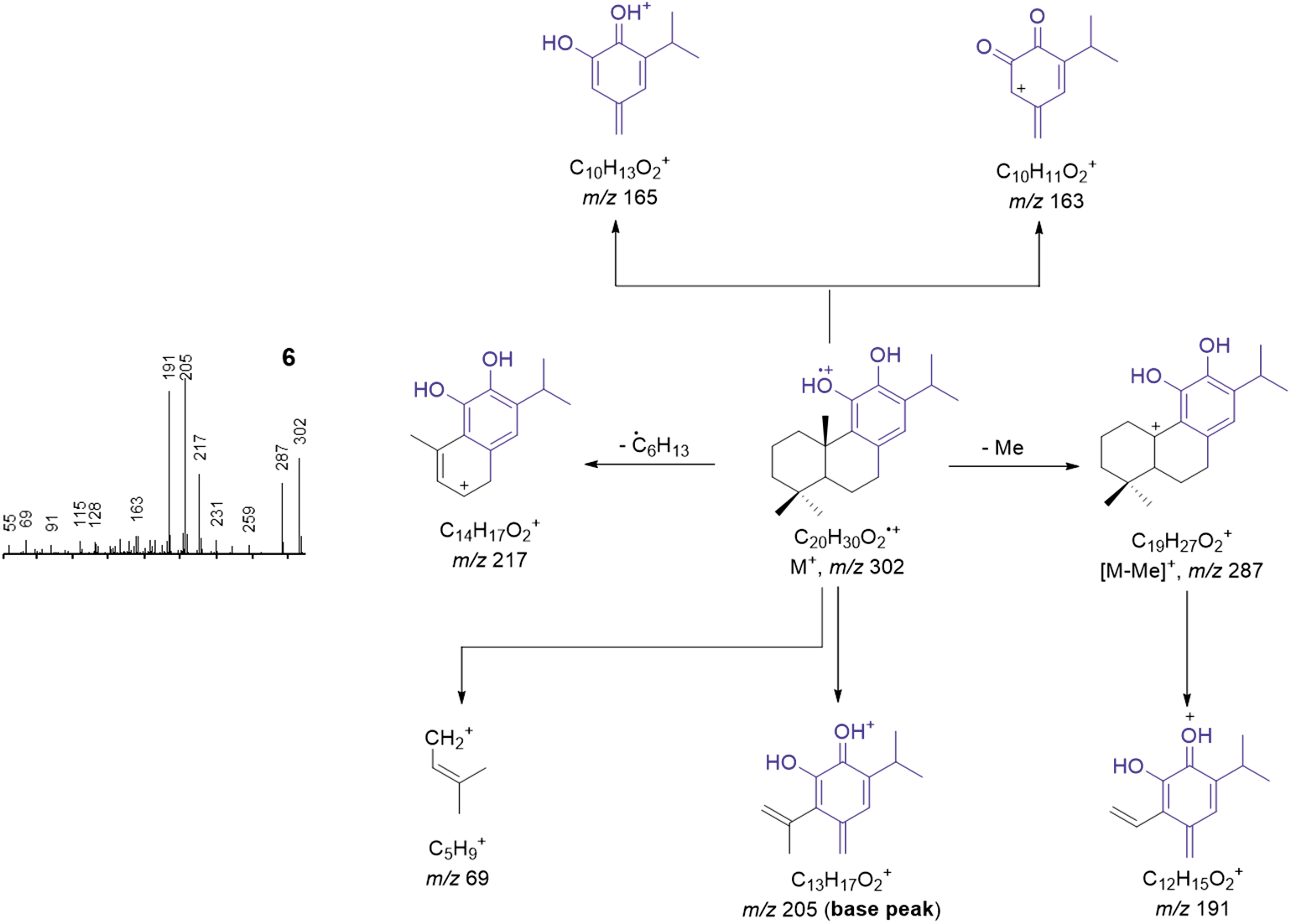
Proposed electron impact ionization (EI) mass spectral fragmentation of 6.

**Fig. S23.**
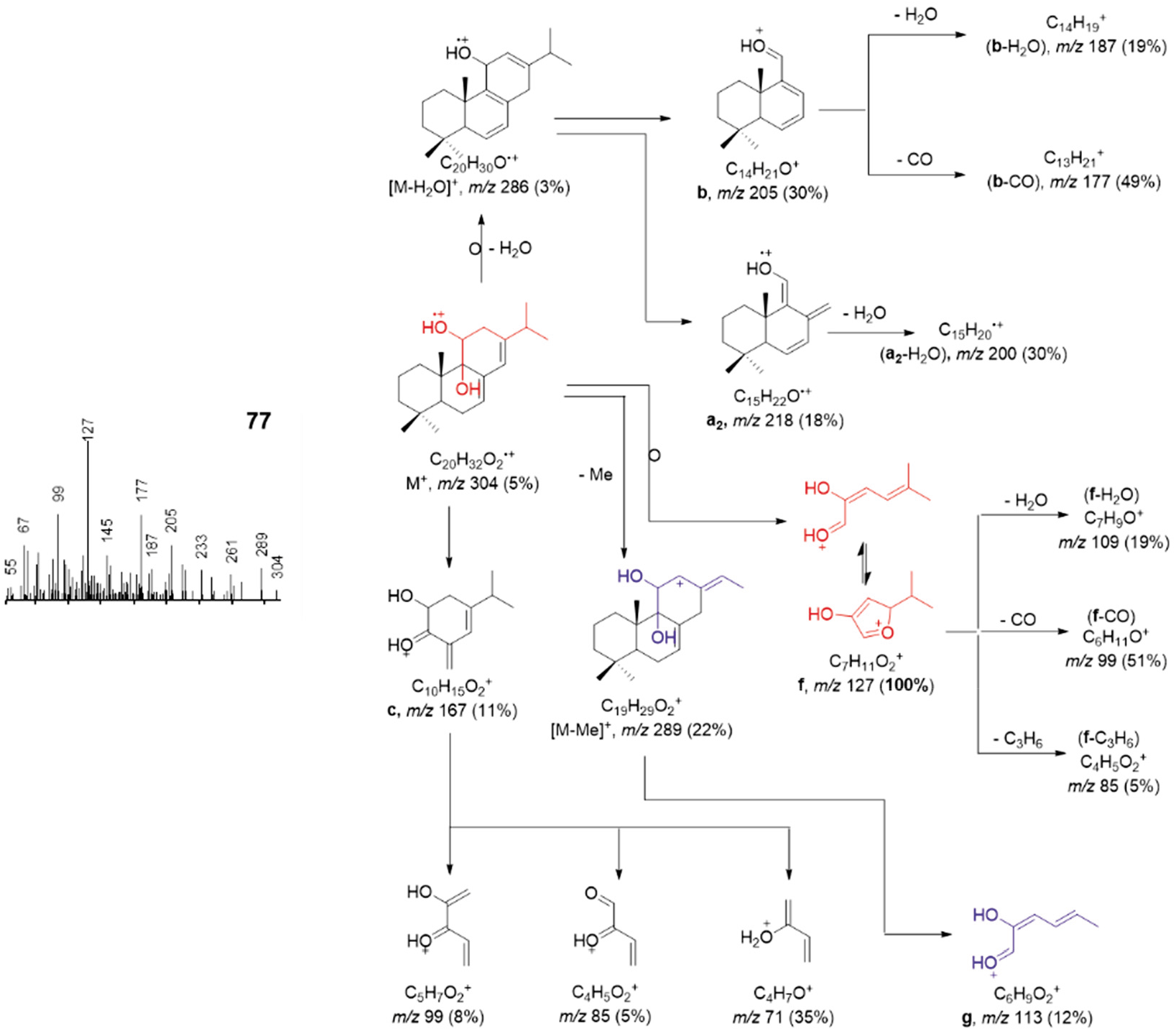
Proposed electron impact ionization (EI) mass spectral fragmentation of 77.

**Fig. S24.**
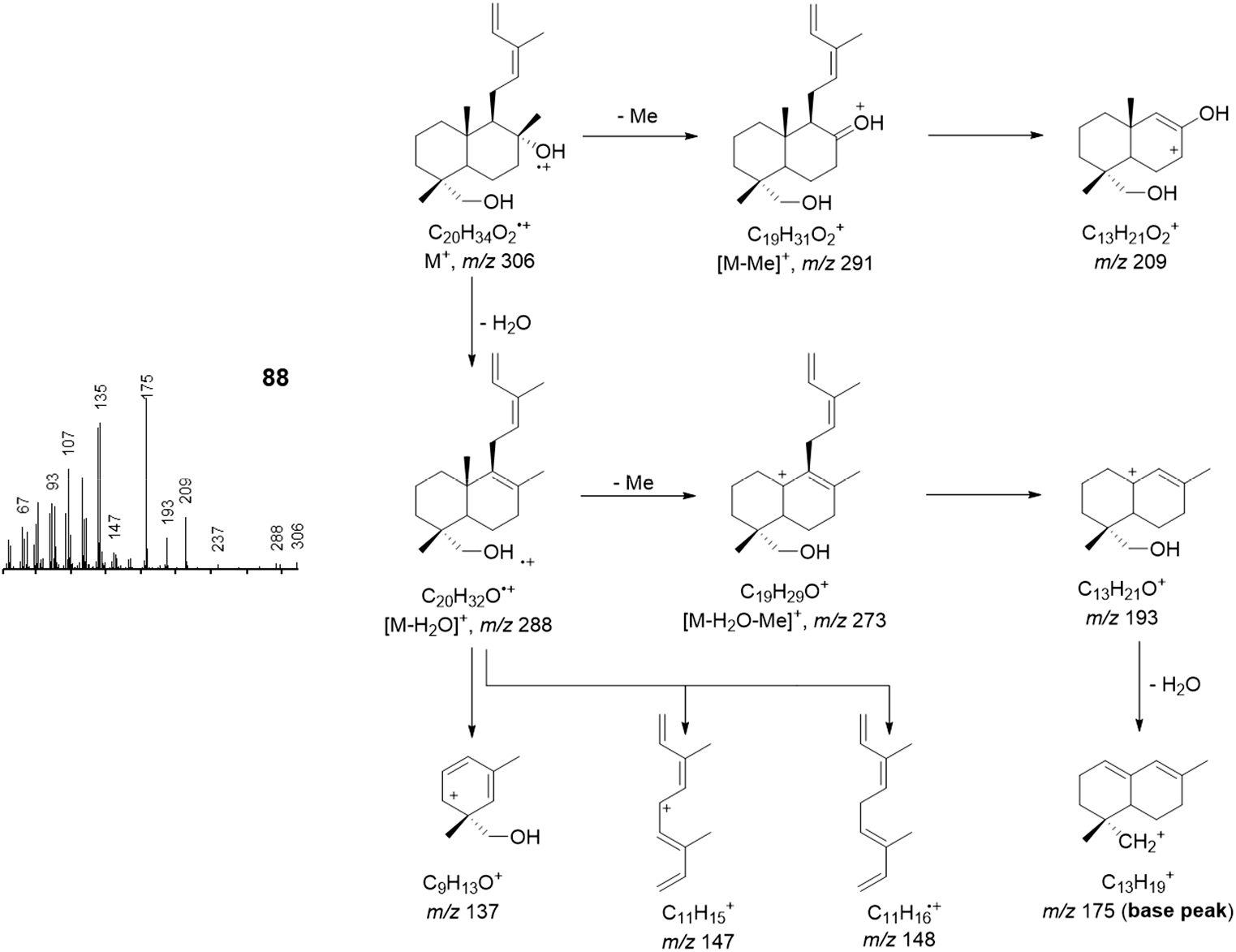
Proposed electron impact ionization (EI) mass spectral fragmentation of 88.

**Fig. S25.**
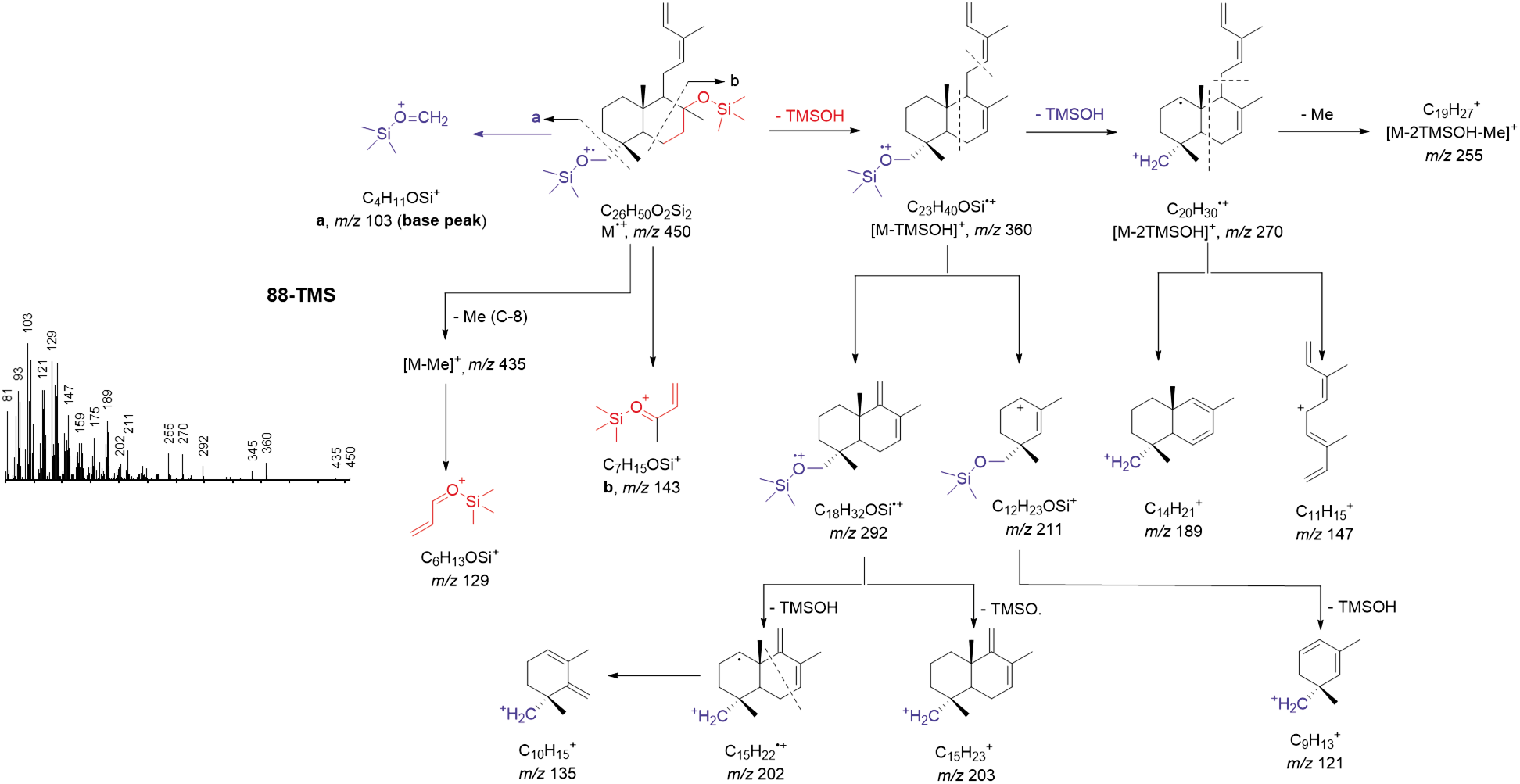
Proposed electron impact ionization (EI) mass spectral fragmentation of silylated 88.

**Fig. S26.**
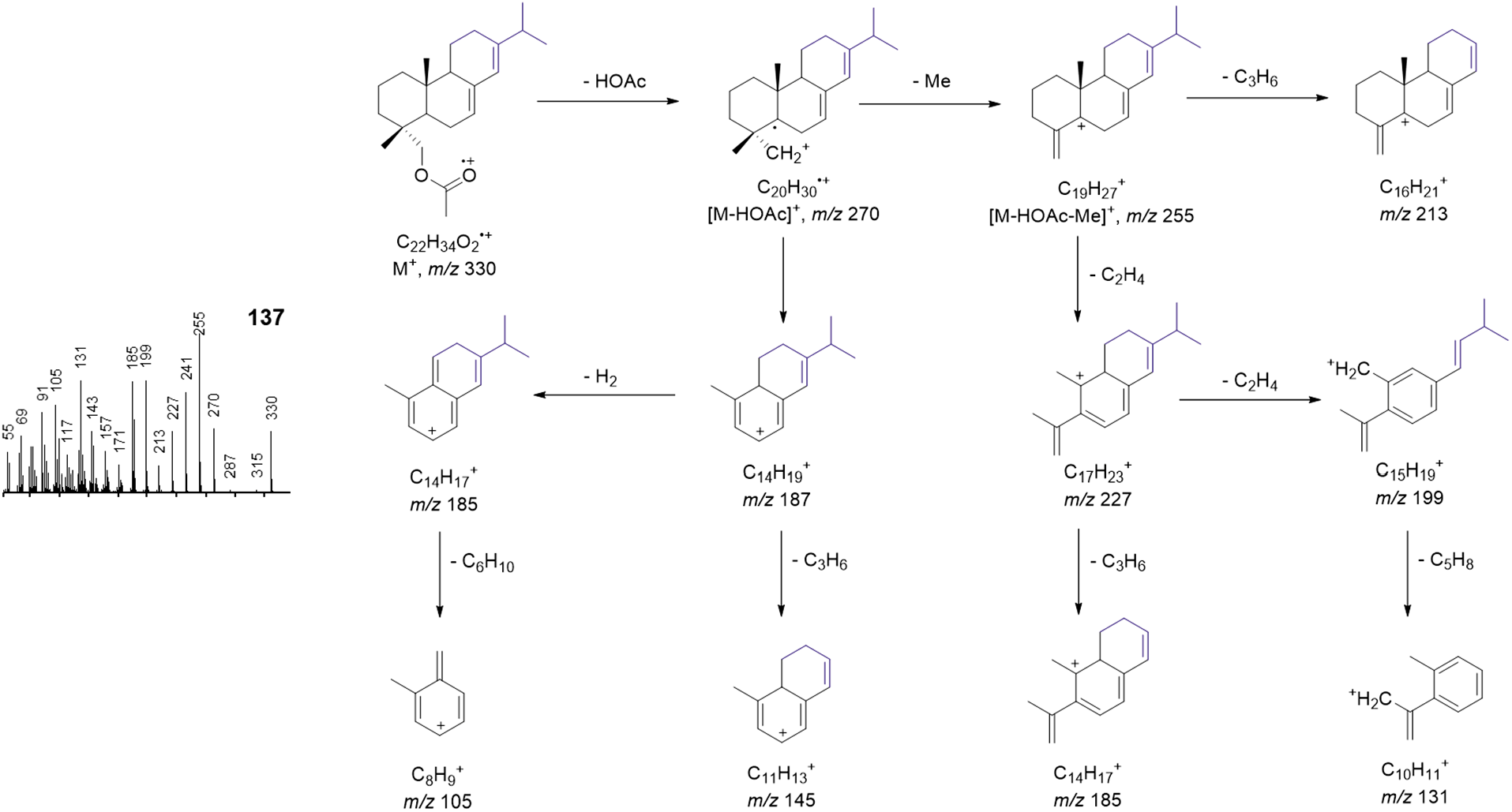
Proposed electron impact ionization (EI) mass spectral fragmentation of 137.

**Fig. S27a.**
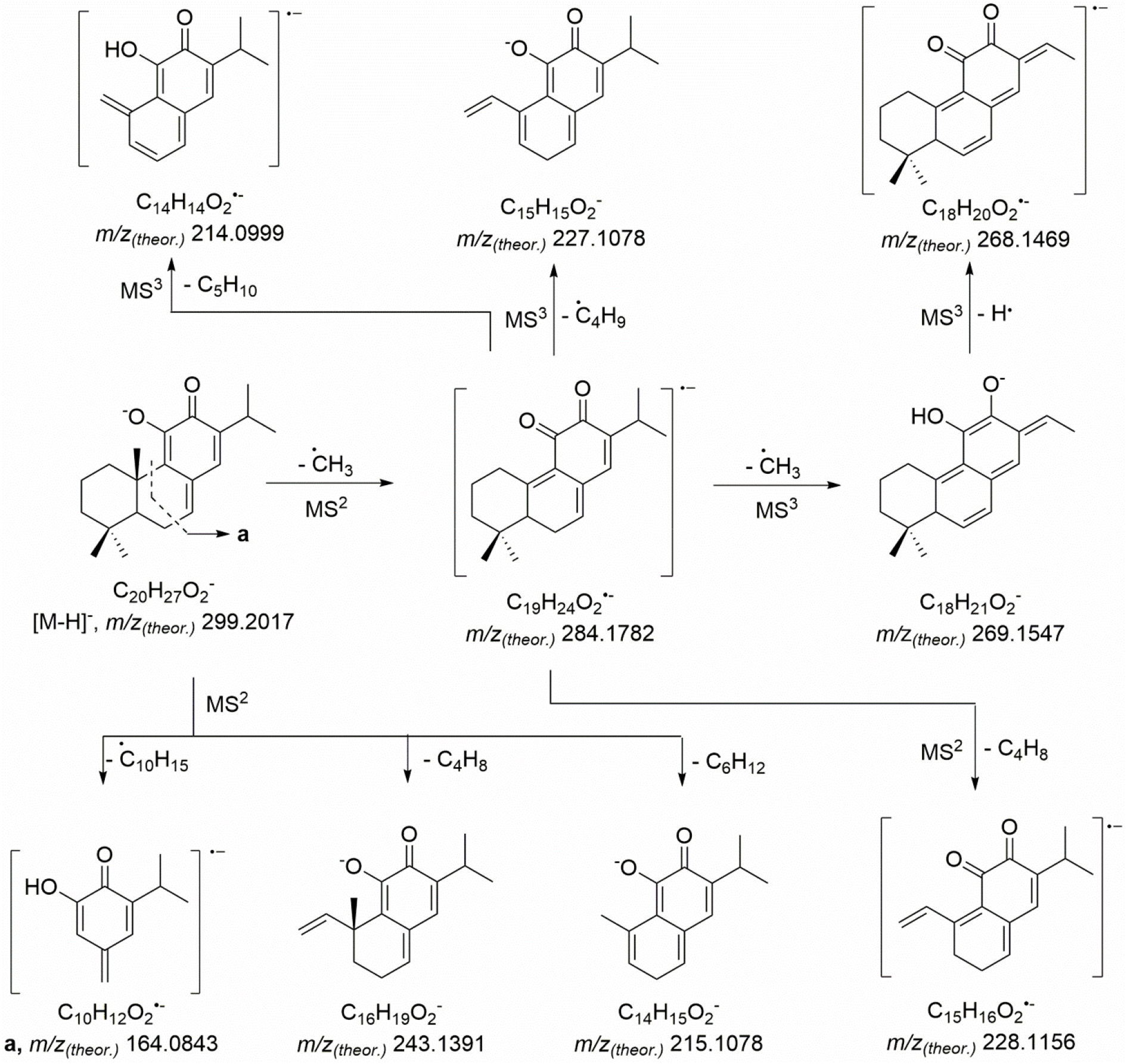
Tandem mass spectrometric fragmentation patterns (MS2, MS3) of 5 (hydroxyferruginol quinone) (peak 7794-11)

**Fig. S27b.**
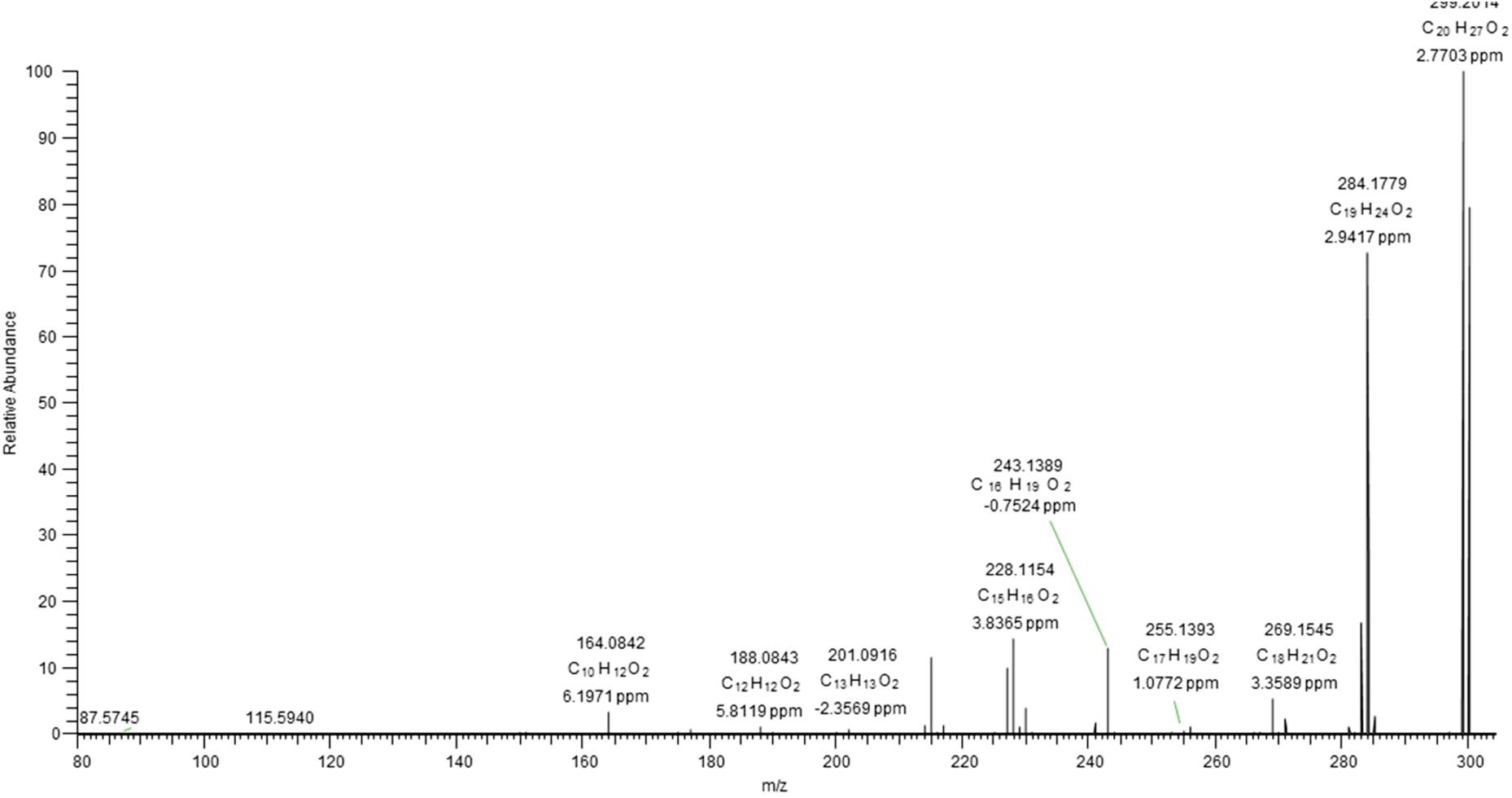
Tandem mass spectrum (MS2) of *m/z* 299.2 ± 1 corresponding to the [M-H]^-^ ion of 5 (hydroxyferruginol quinone) (*m/z* 299.2014, 1.34 ppm). The high resolution (HR) spectra were acquired with CID activation (normalized collision energy 35%) at the resolution of 15000 by a hybrid ESI-LIT-Orbitrap mass spectrometer (Orbitrap Elite, Thermo Fisher Scientific, Bremen, Germany) operated in negative ion mode, after on-line separation of peak 7794-11 by RP-UHPLC as described in the methods section.

**Fig. S27c.**
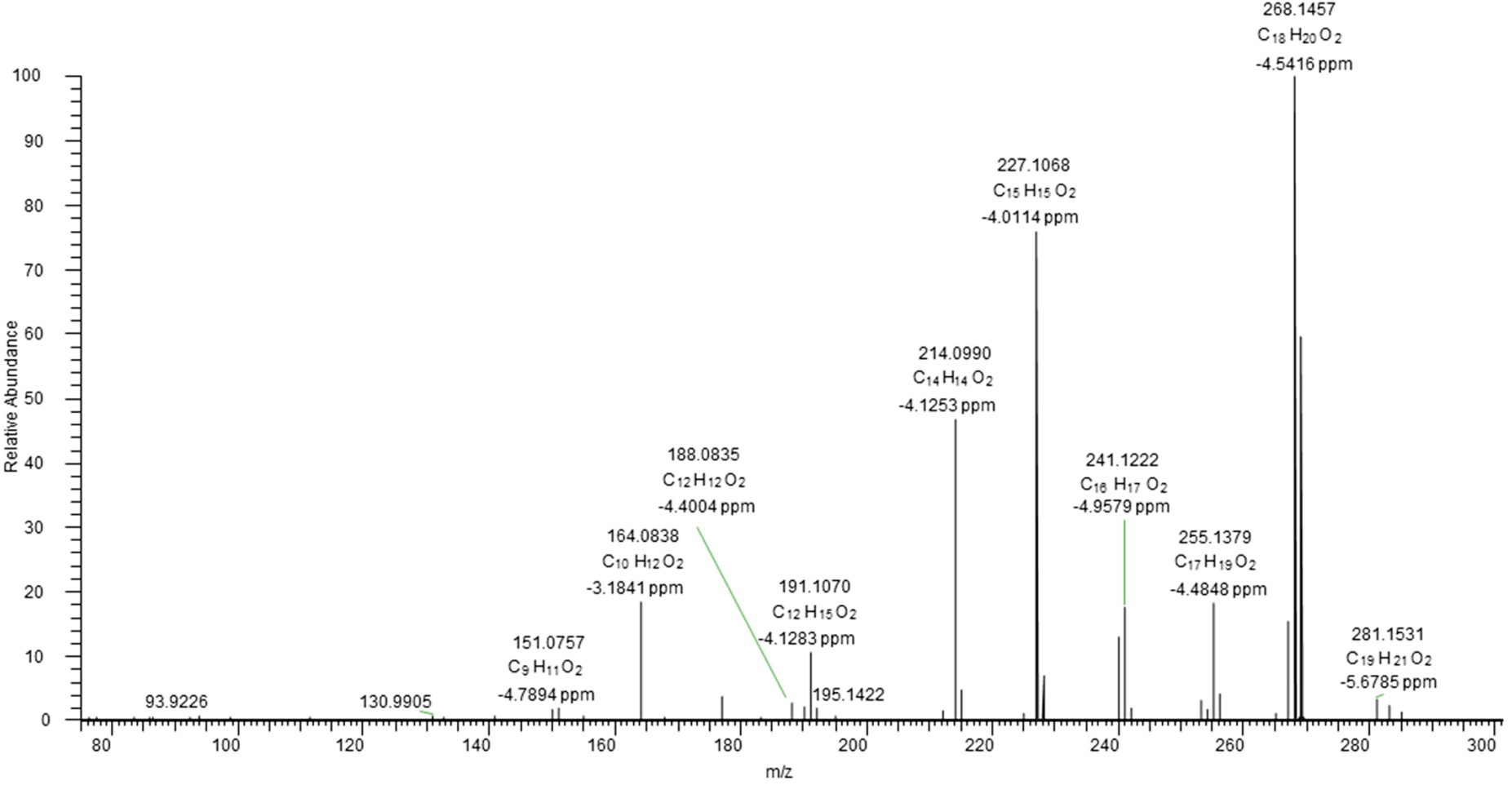
Tandem mass spectrum (MS3) of *m/z* 299.2 ± 1 → *m/z* 284.2 corresponding to the [M-H]^-^ ion of 5 (hydroxyferruginol quinone) (*m/z* 299.2014, 1.34 ppm). The high resolution (HR) spectra were acquired with CID activation (normalized collision energy 45%) at the resolution of 15000 by a hybrid ESI-LIT-Orbitrap mass spectrometer (Orbitrap Elite, Thermo Fisher Scientific, Bremen, Germany) operated in negative ion mode, after on-line separation of paek 7794-11 by RP-UHPLC as described in the Methods section.

**Fig. S27d.**
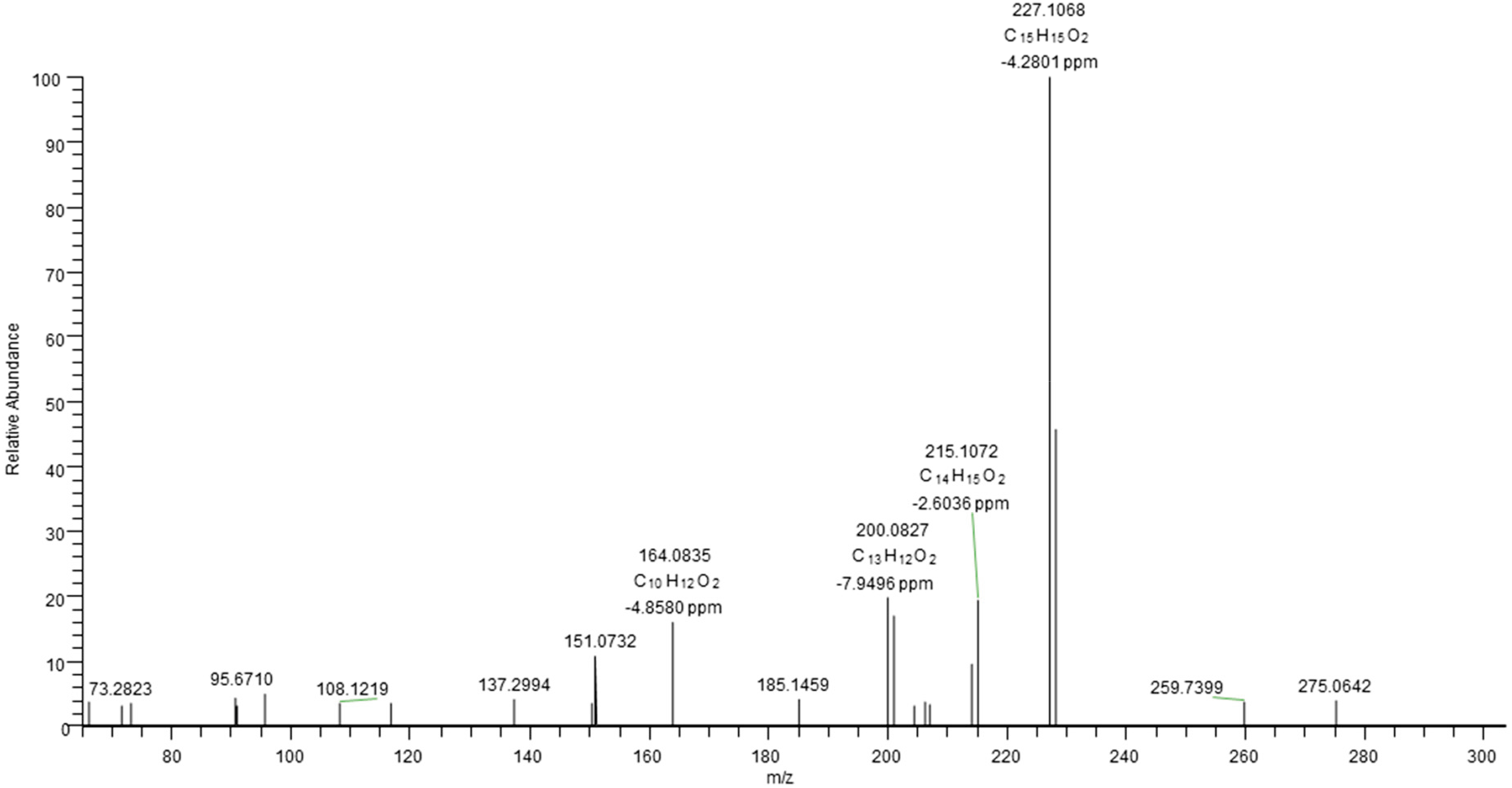
Tandem mass spectrum (MS3) of *m/z* 299.2 ± 1 → *m/z* 243.1 corresponding to the [M-H]^-^ ion of 5 (hydroxyferruginol quinone) (*m/z* 299.2014, 1.34 ppm). The high resolution (HR) spectra were acquired with CID activation (normalized collision energy 45%) at the resolution of 15000 by a hybrid ESI-LIT-Orbitrap mass spectrometer (Orbitrap Elite, Thermo Fisher Scientific, Bremen, Germany) operated in negative ion mode, after on-line separation of peak 7794-11 by RP-UHPLC as described in the Methods section.

**Fig. S27e.**
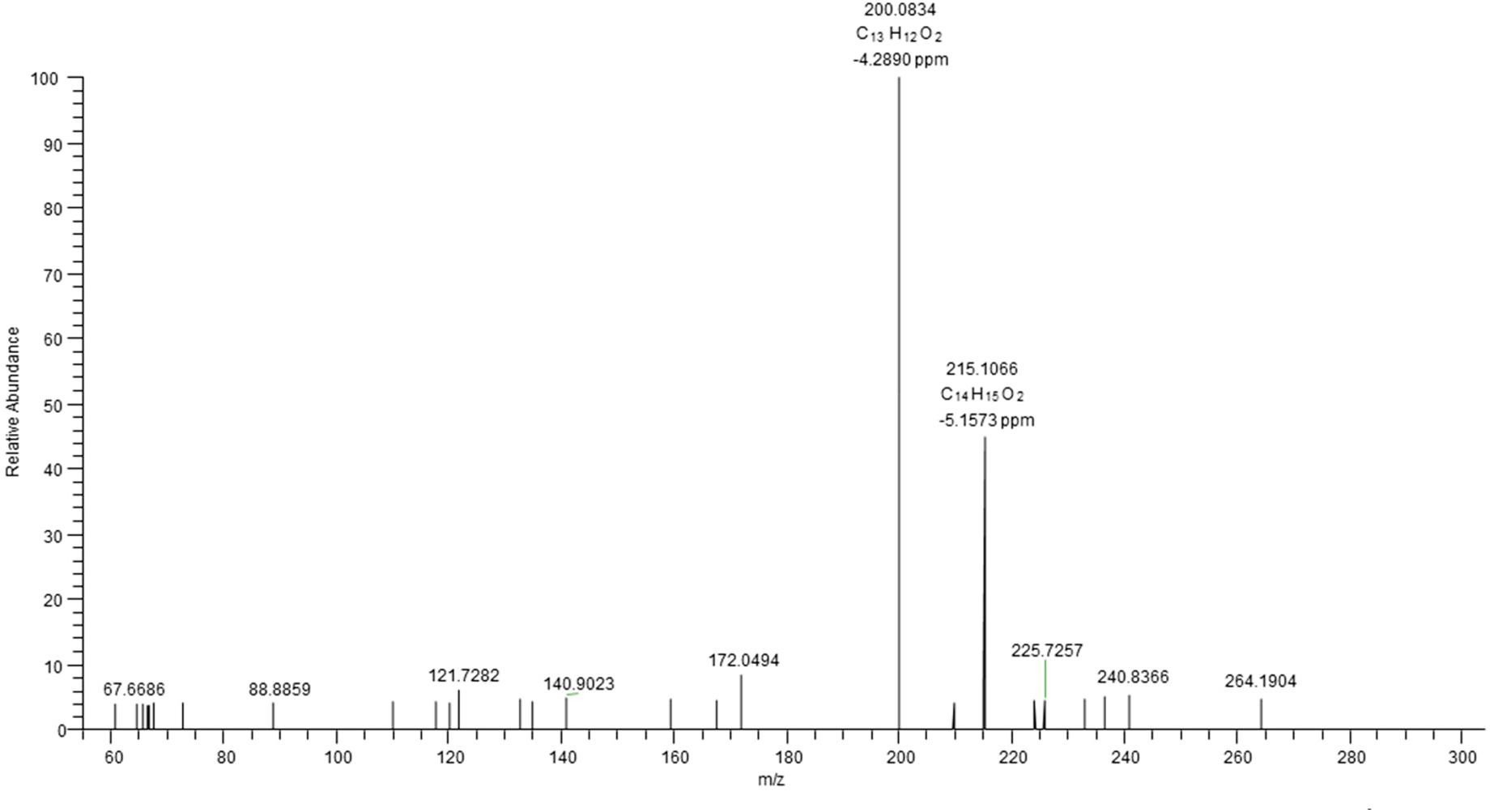
Tandem mass spectrum (MS3) of *m/z* 299.2 ± 1 → *m/z* 215.1 corresponding to the [M-H]^-^ ion of 5 (hydroxyferruginol quinone) (*m/z* 299.2014, 1.34 ppm). The high resolution (HR) spectra were acquired with CID activation (normalized collision energy 35%) at the resolution of 15000 by a hybrid ESI-LIT-Orbitrap mass spectrometer (Orbitrap Elite, Thermo Fisher Scientific, Bremen, Germany) operated in negative ion mode, after on-line separation of the sample 7794_11 by RP- UHPLC as described in the Methods section.

**Fig. S27f.**
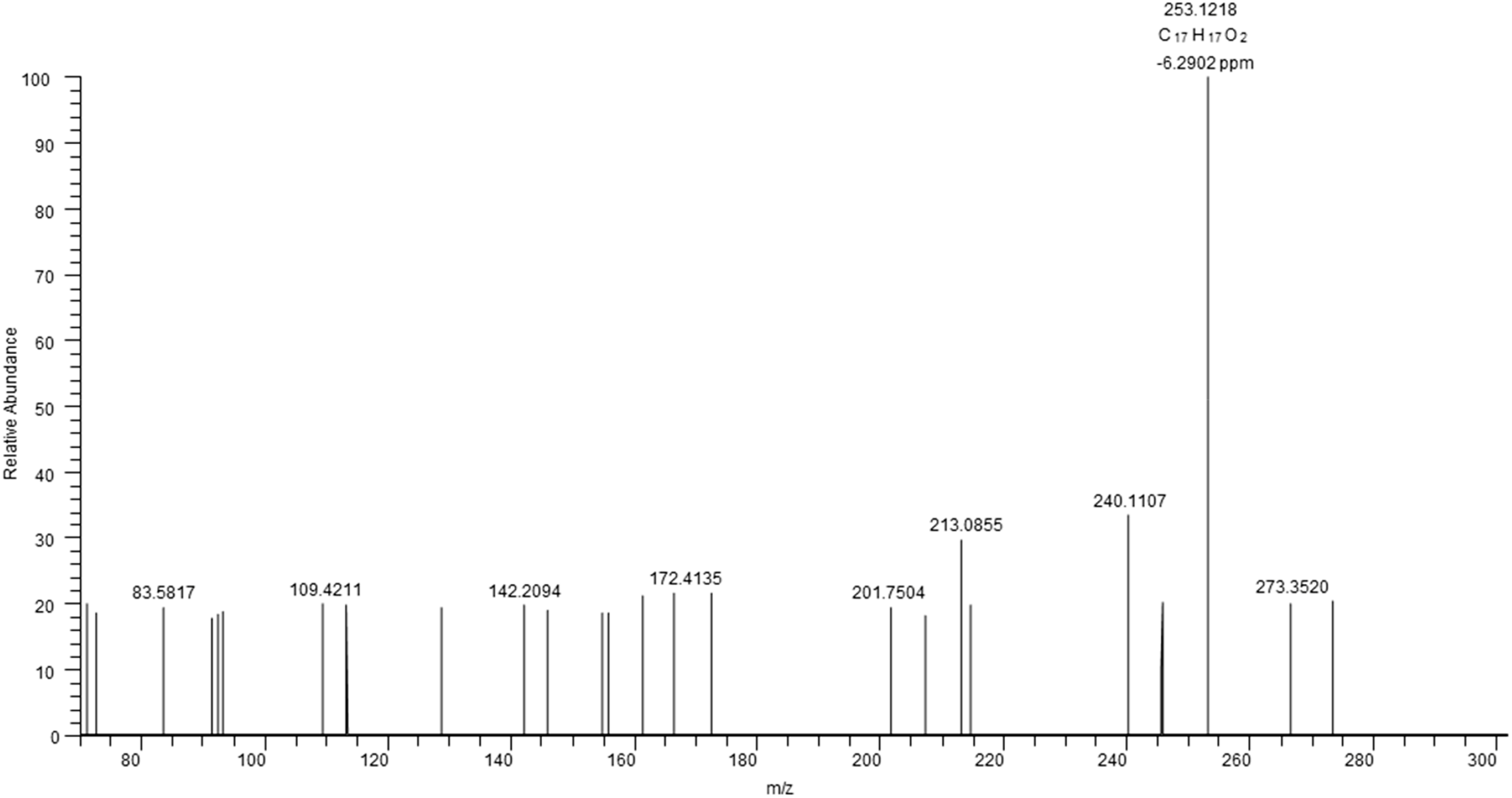
Tandem mass spectrum (MS4) of *m/z* 299.2 ± 1 → *m/z* 283.2 → *m/z* 268.2 corresponding to the [M-H]^-^ ion of 5 (hydroxyferruginol quinone) (*m/z* 299.2014, 1.34 ppm). The high resolution (HR) spectra were acquired with CID activation (normalized collision energy 45%) at the resolution of 15000 by a hybrid ESI-LIT-Orbitrap mass spectrometer (Orbitrap Elite, Thermo Fisher Scientific, Bremen, Germany) operated in negative ion mode, after on-line separation of peak 7794-11 by RP-UHPLC as described in the Methods section.

**Fig. S28a.**
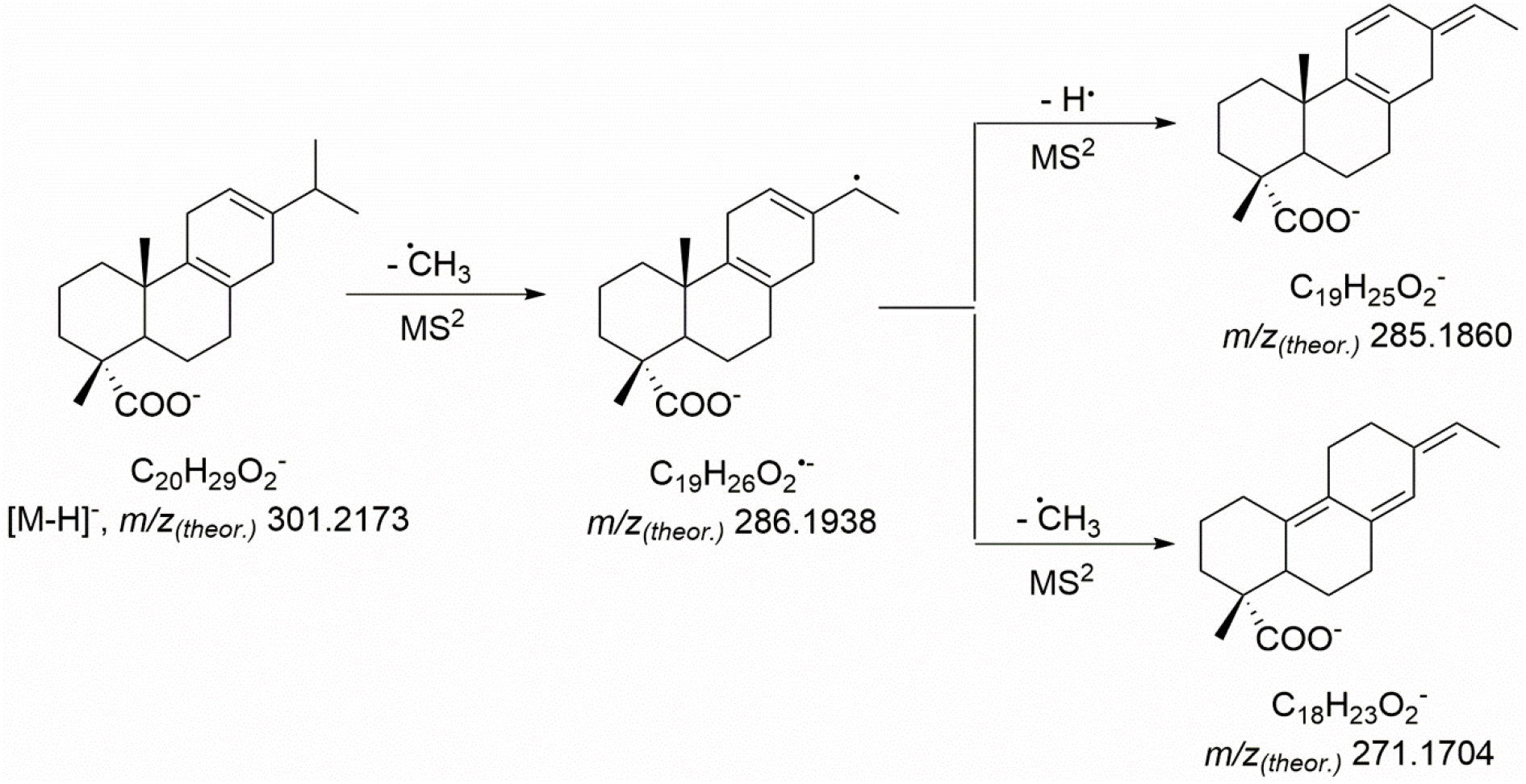
Tandem mass spectrometric fragmentation (MS2) patterns of 19 (miltiradienic-18-acid), peak 7797-7

**Fig. S28b.**
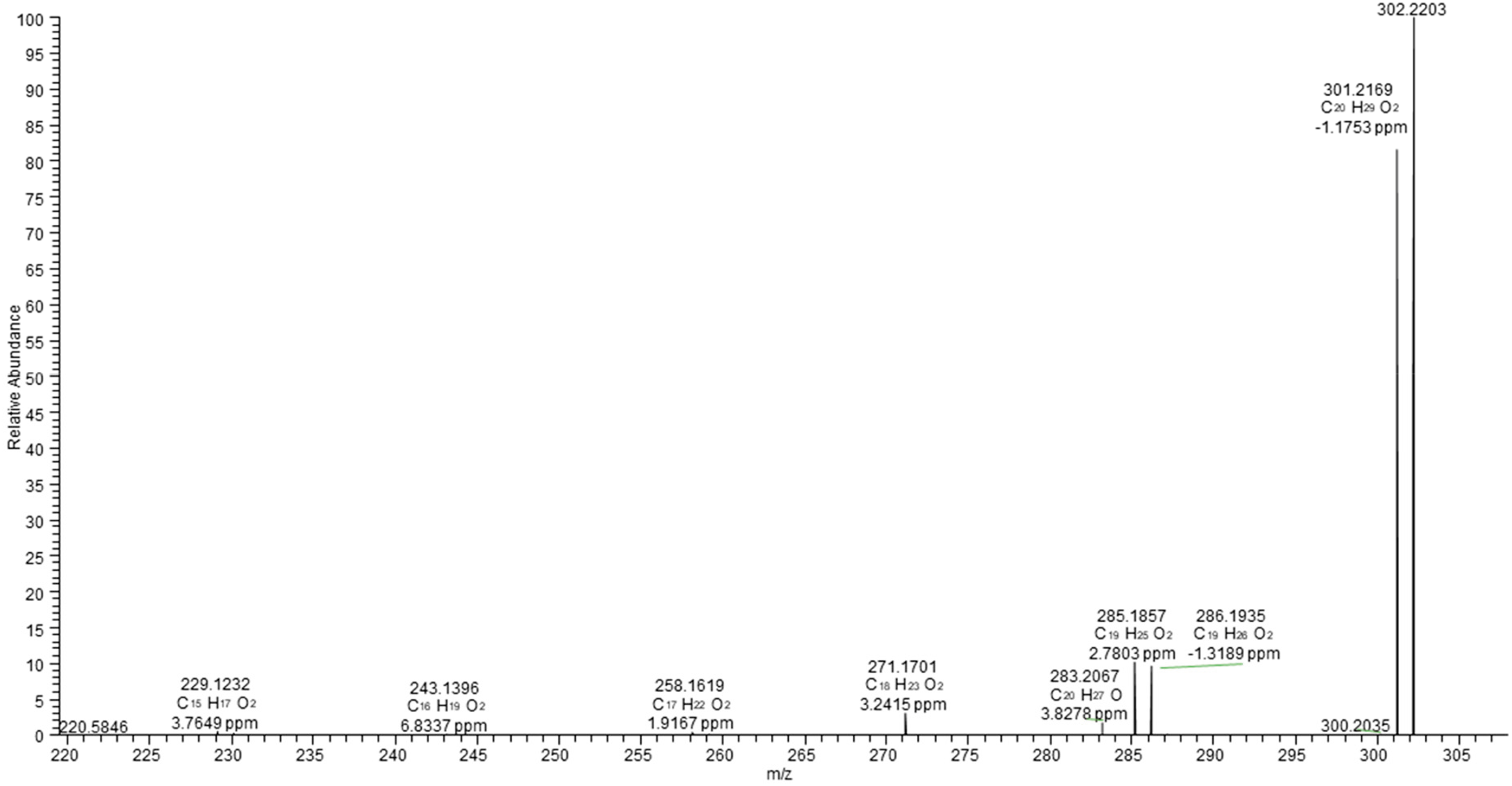
Tandem mass spectrum (MS2) of *m/z* 301.2 ± 1 corresponding to the [M-H]^-^ ion of 19 (miltiradienic-18-acid) (*m/z* 301.2169, 1.33 ppm). The high resolution (HR) spectra were acquired with CID activation (normalized collision energy 35%) at the resolution of 15000 by a hybrid ESI-LIT- Orbitrap mass spectrometer (Orbitrap Elite, Thermo Fisher Scientific, Bremen, Germany) operated in negative ion mode, after on-line separation of peak 7797-7 by RP-UHPLC as described in the Methods section.

**Fig. S29a.**
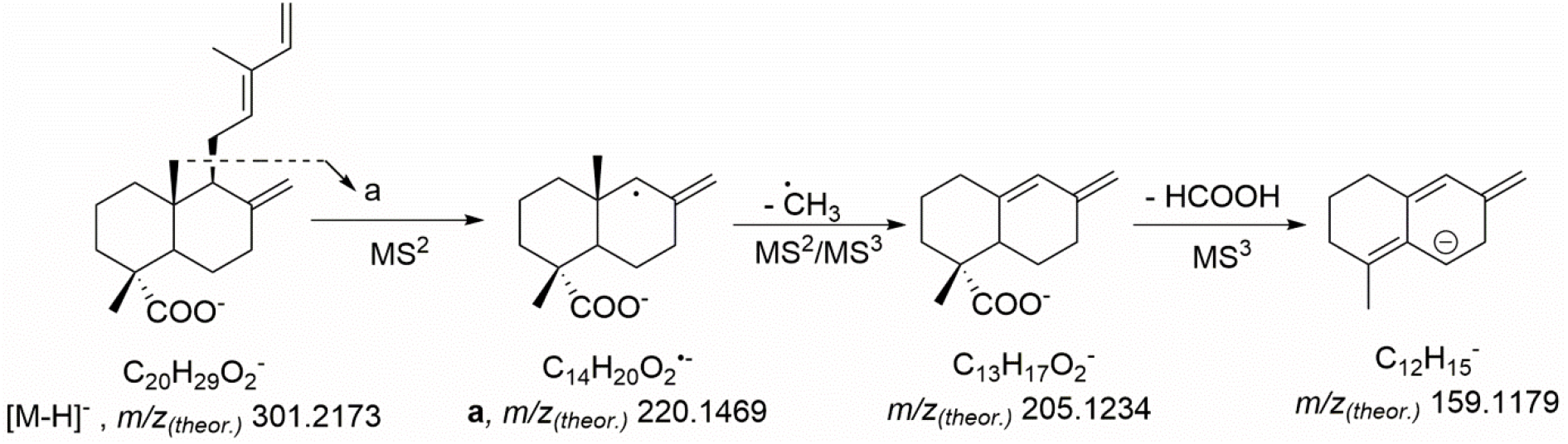
Tandem mass spectrometric fragmentation patterns (MS2 and MS3) of 117 (*trans*- communic-18-acid, peak 7802-4).

**Fig. S29b.**
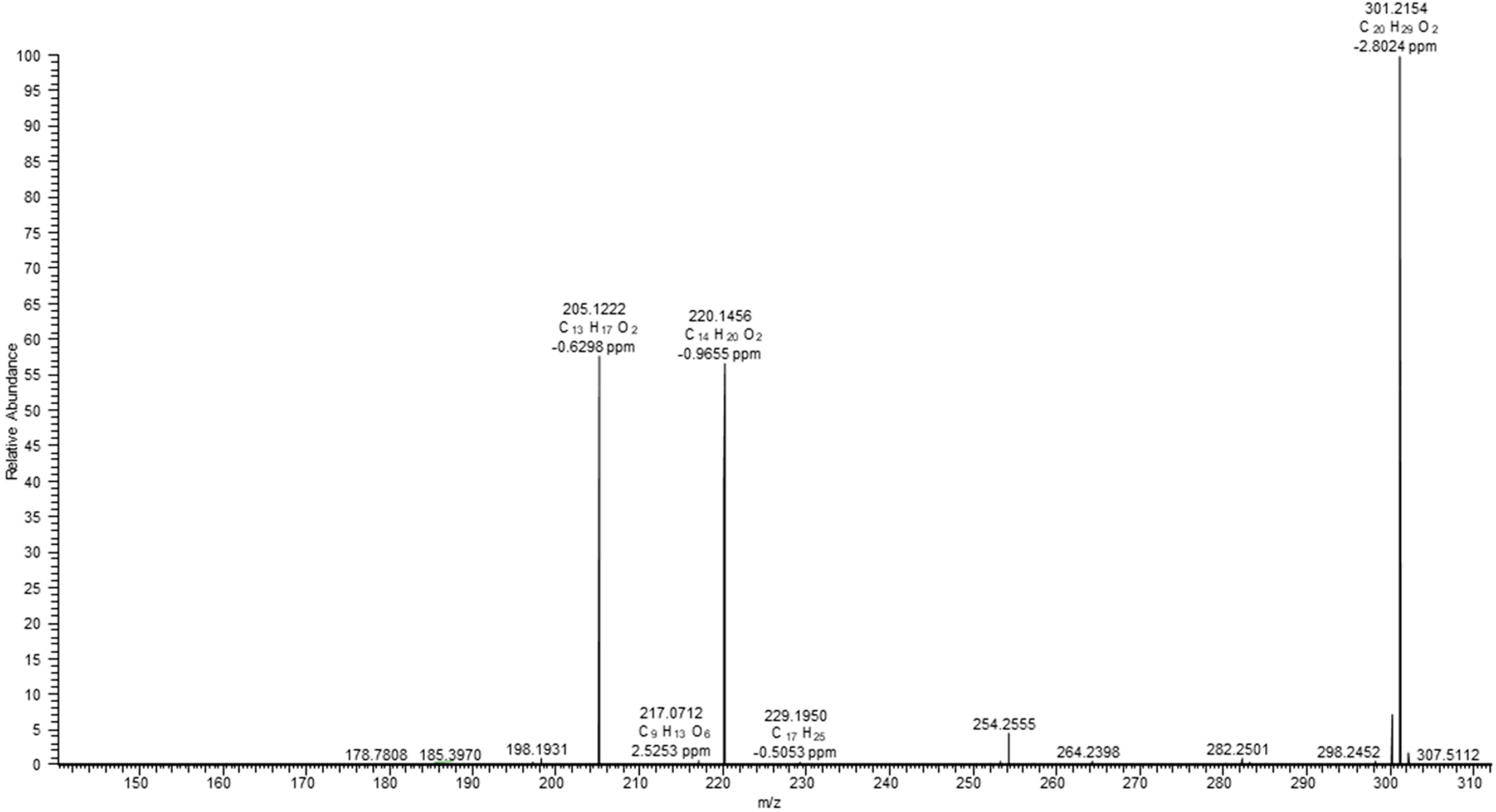
Tandem mass spectrum (MS2) of *m/z* 301.2 ± 1 corresponding to the [M-H]^-^ ion of 117 (*trans*-communic-18-acid) (*m/z* 301.2168, 1.66 ppm). The high resolution (HR) spectra were acquired with CID activation (normalized collision energy 30%) at the resolution of 15000 by a hybrid ESI-LIT- Orbitrap mass spectrometer (Orbitrap Elite, Thermo Fisher Scientific, Bremen, Germany) operated in negative ion mode, after on-line separation of peak 7802-4 by RP-UHPLC as described in the Methods section.

**Fig. S29c.**
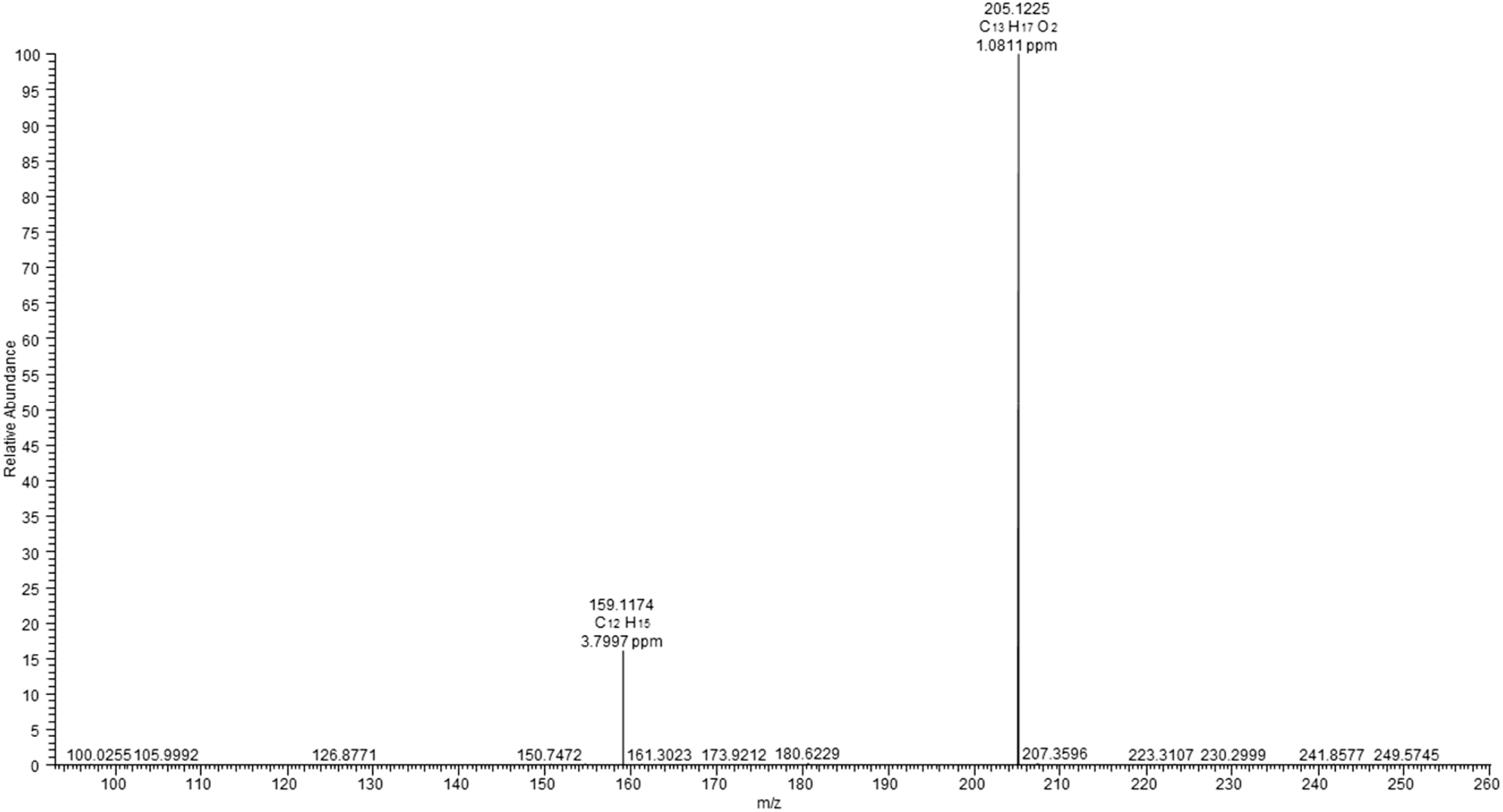
Tandem mass spectrum (MS3) of *m/z* 301.2 ± 1 → *m/z* 205.1 corresponding to the [M-H]^-^ ion of 117 (*trans*-communic-18-acid) (*m/z* 301.2168, 1.66 ppm). The high resolution (HR) spectra were acquired with CID activation (normalized collision energy 25%) at the resolution of 15000 by a hybrid ESI-LIT-Orbitrap mass spectrometer (Orbitrap Elite, Thermo Fisher Scientific, Bremen, Germany) operated in negative ion mode, after on-line separation of peak 7802-4 by RP-UHPLC as described in the Methods section.

**Fig. S29d.**
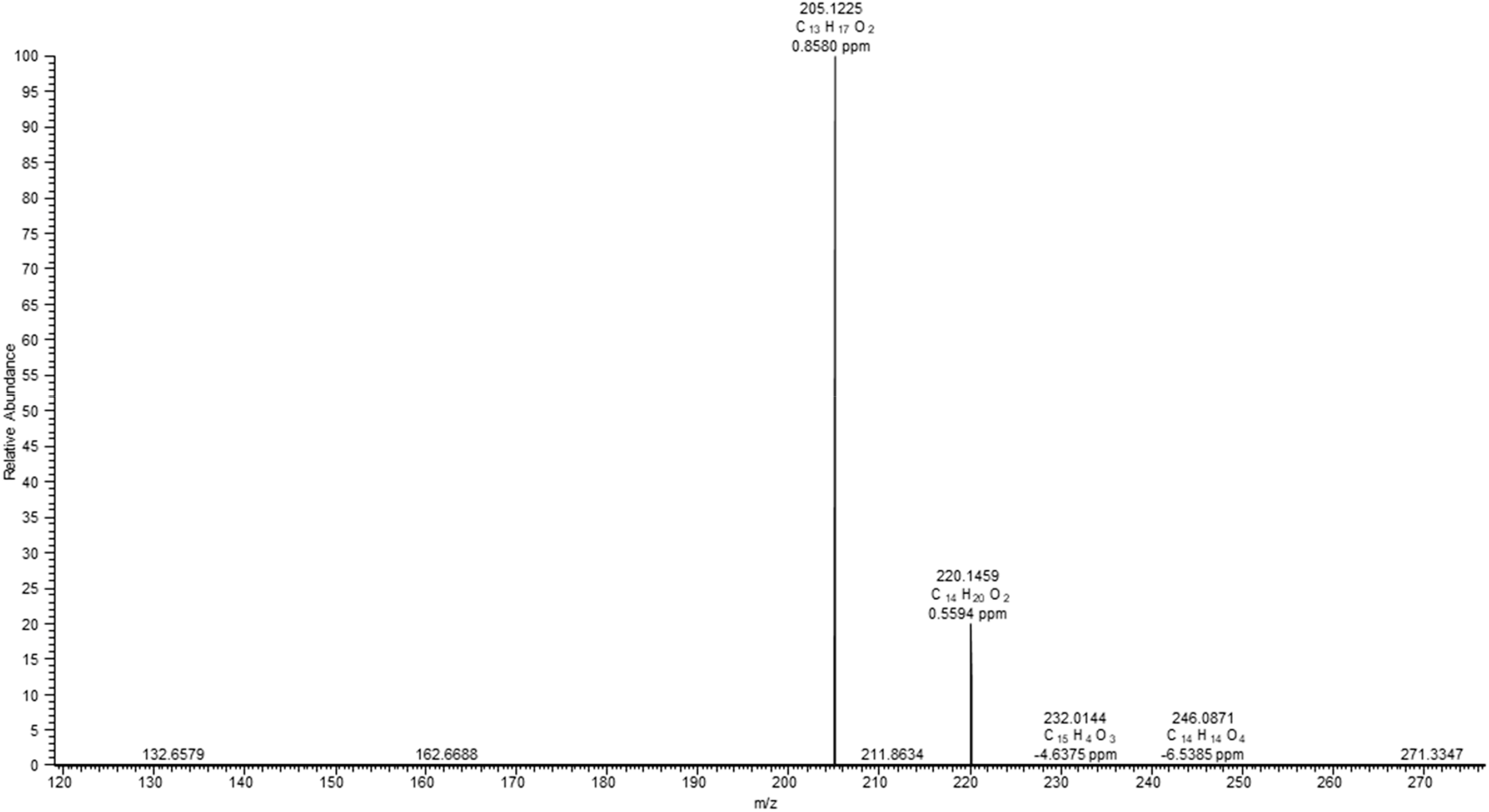
Tandem mass spectrum (MS3) of *m/z* 301.2 ± 1 → *m/z* 220.15 corresponding to the [M-H]^-^ ion of 117 (*trans*-communic-18-acid) (*m/z* 301.2168, 1.66 ppm). The high resolution (HR) spectra were acquired with CID activation (normalized collision energy 25%) at the resolution of 15000 by a hybrid ESI-LIT-Orbitrap mass spectrometer (Orbitrap Elite, Thermo Fisher Scientific, Bremen, Germany) operated in negative ion mode, after on-line separation of peak 7802-4 by RP-UHPLC as described in the methods section.

**Fig. S30a.**
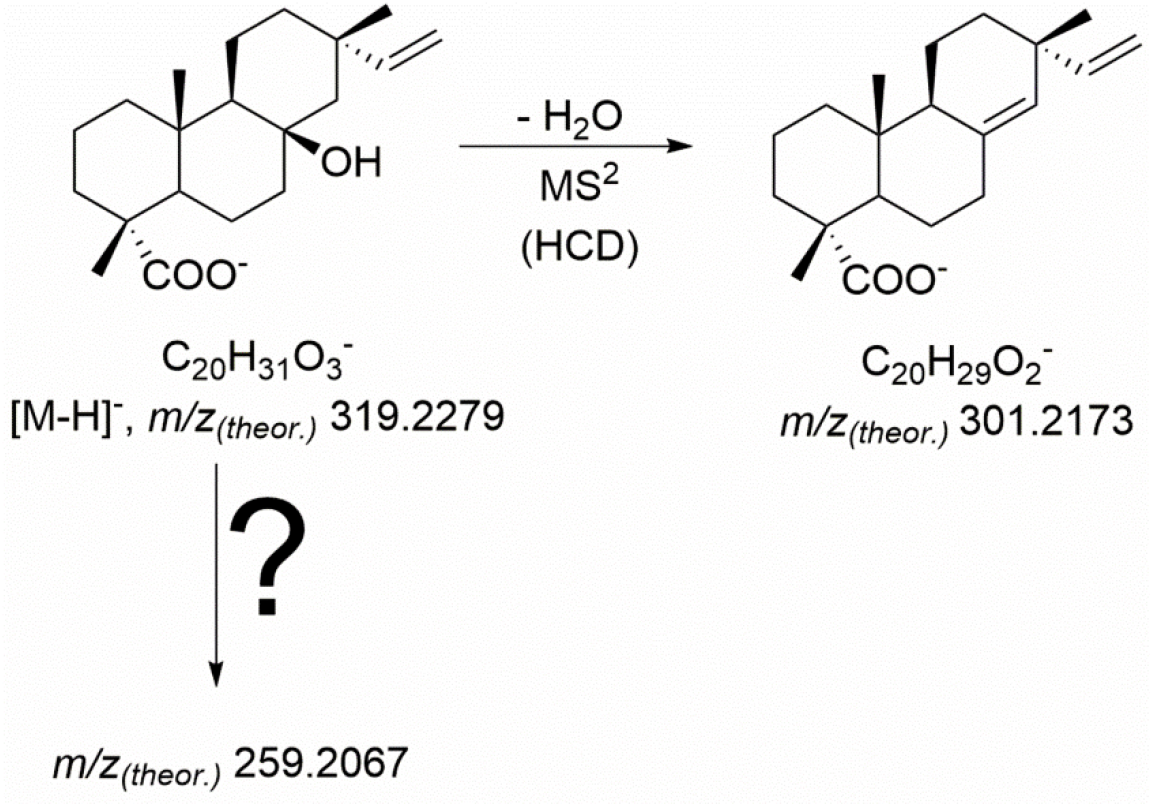
Tandem mass spectrometric fragmentation patterns (MS2) of 120 (nezukolic acid, peak 7808-2).

**Fig. S30b.**
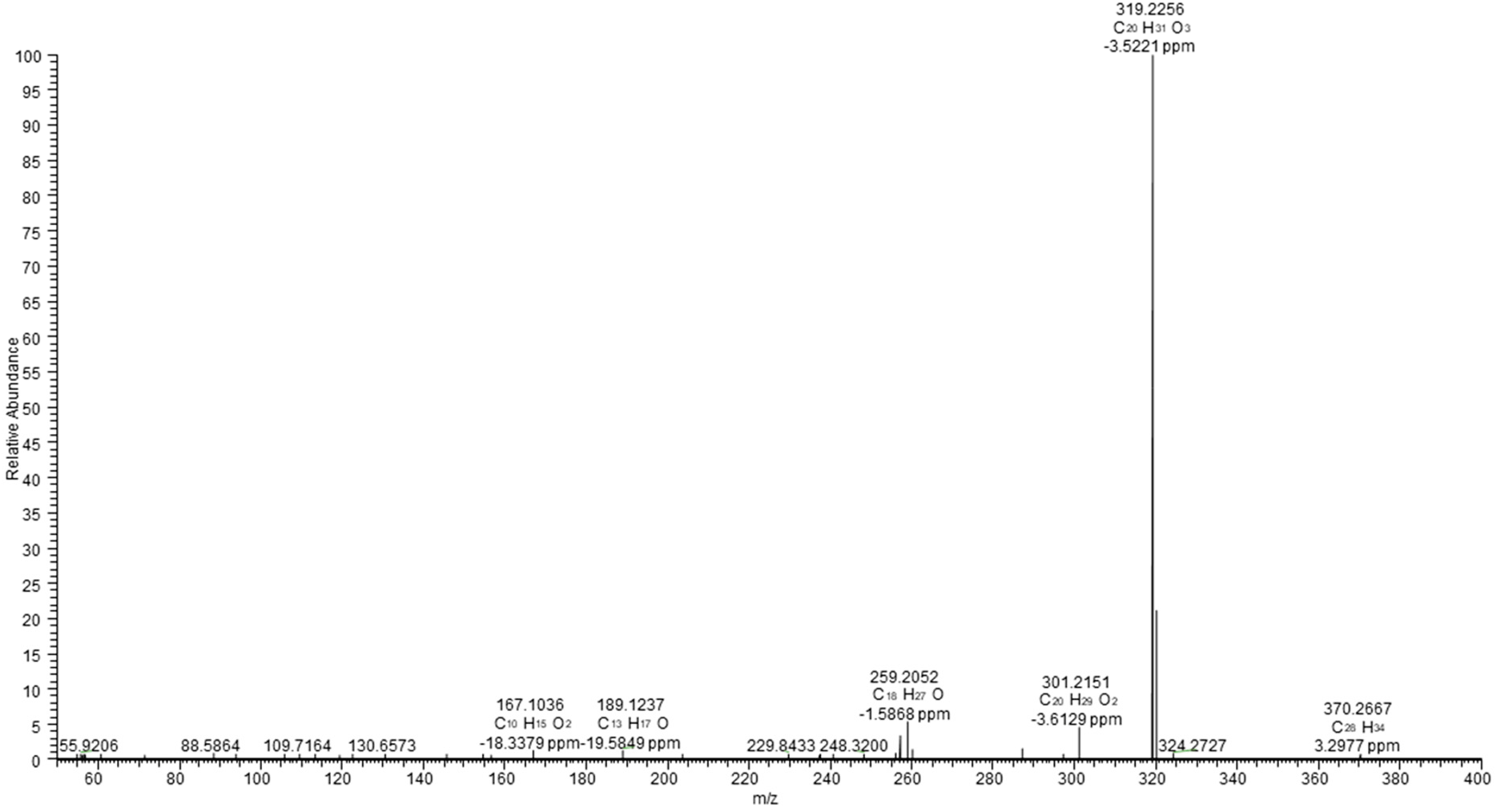
Tandem mass spectrum (MS2) of *m/z* 319.2 ± 1 corresponding to the [M-H]^-^ ion of 120 (nezukolic acid) (*m/z* 319.2274, 1.57 ppm). The high resolution (HR) spectra were acquired with HCD activation (170 eV) at the resolution of 15000 by a hybrid ESI-LIT-Orbitrap mass spectrometer (Orbitrap Elite, Thermo Fisher Scientific, Bremen, Germany) operated in negative ion mode, after on- line separation of peak 7808-2 by RP-UHPLC as described in the Methods section.

**Fig. S31a.**
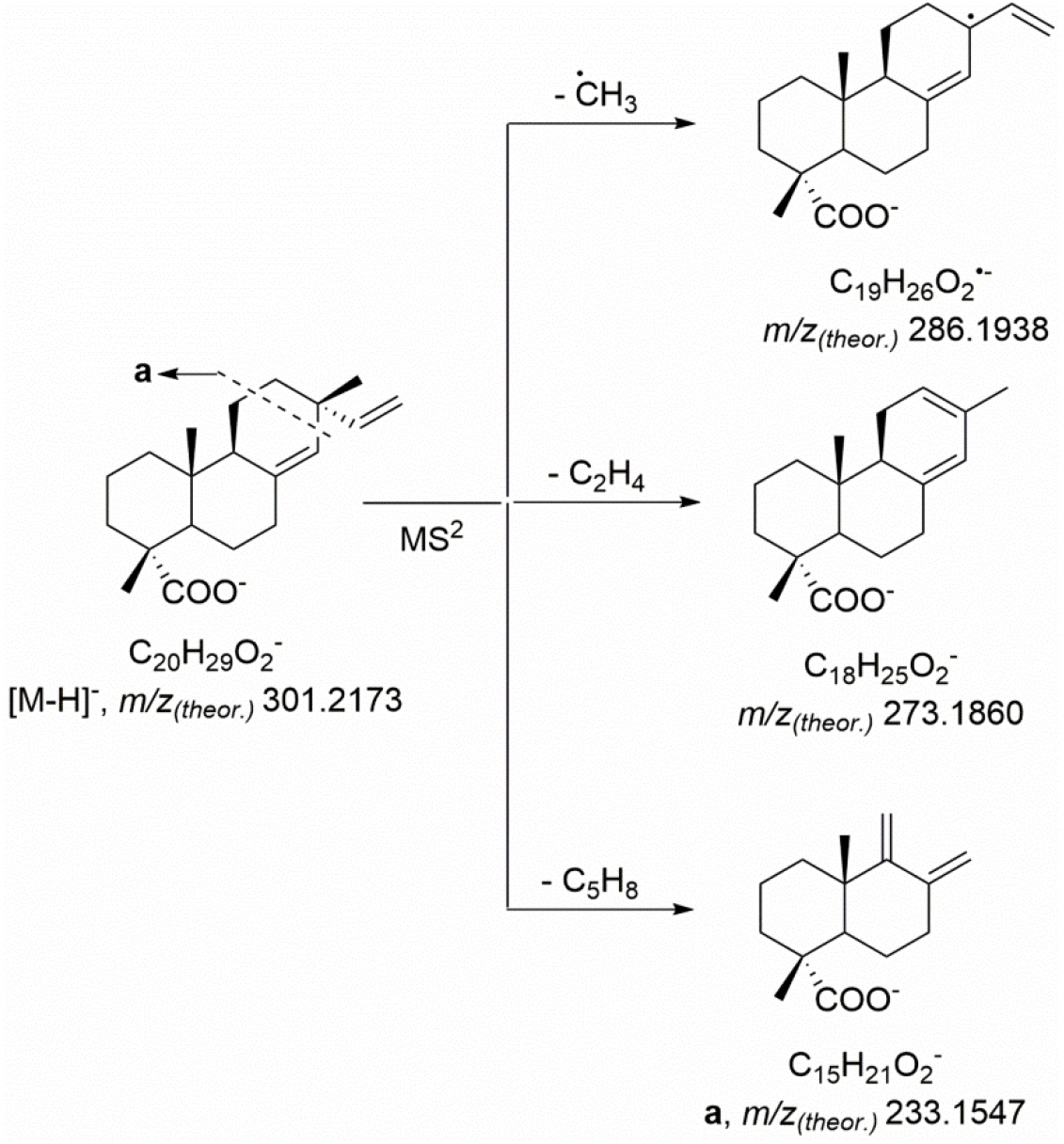
Advanced Tandem mass spectrometric fragmentation patterns (MS2) of 121 (sandaracopimaric-18-acid, peak 7813-3).

**Fig. S31b.**
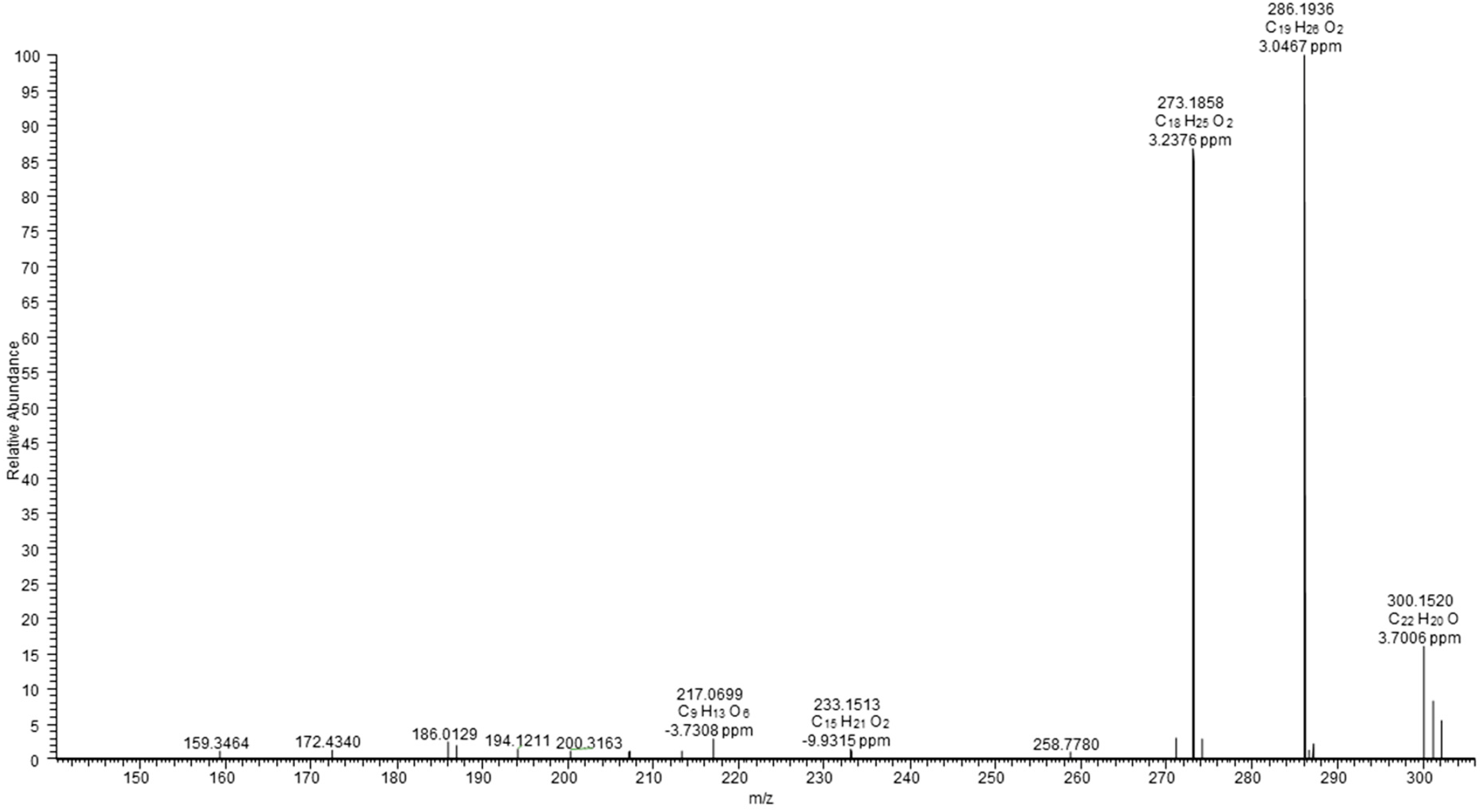
Tandem mass spectrum (MS2) of *m/z* 301.2 ± 1 corresponding to the [M-H]^-^ ion of 121 (sandaracopimaric-18-acid) (*m/z* 301.2168, 1.66 ppm). The high resolution (HR) spectra were acquired with CID activation (normalized collision energy 35%) at the resolution of 15000 by a hybrid ESI-LIT-Orbitrap mass spectrometer (Orbitrap Elite, Thermo Fisher Scientific, Bremen, Germany) operated in negative ion mode, after on-line separation of peak 7813-3 by RP-UHPLC as described in the Methods section.

**Fig. S31c.**
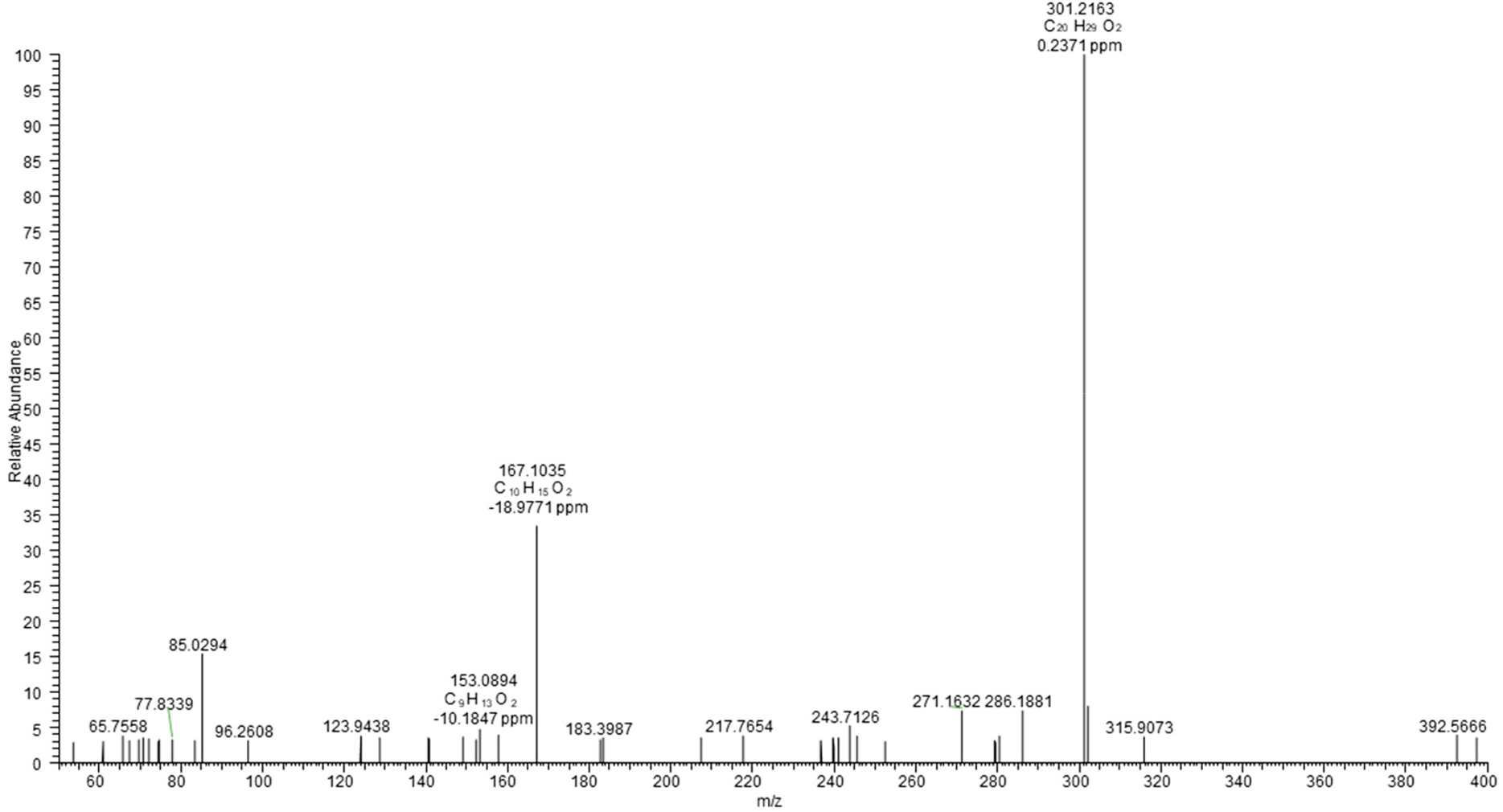
Tandem mass spectrum (MS2) of *m/z* 301.2 ± 1 corresponding to the [M-H]^-^ ion of 121 (sandaracopimaric-18-acid) (*m/z* 301.2168, 1.66 ppm). The high resolution (HR) spectra were acquired with HCD activation (170 eV) at the resolution of 15000 by a hybrid ESI-LIT-Orbitrap mass spectrometer (Orbitrap Elite, Thermo Fisher Scientific, Bremen, Germany) operated in negative ion mode, after on-line separation of peak 7813-3 by RP-UHPLC as described in the methods section.

**Fig. S31d.**
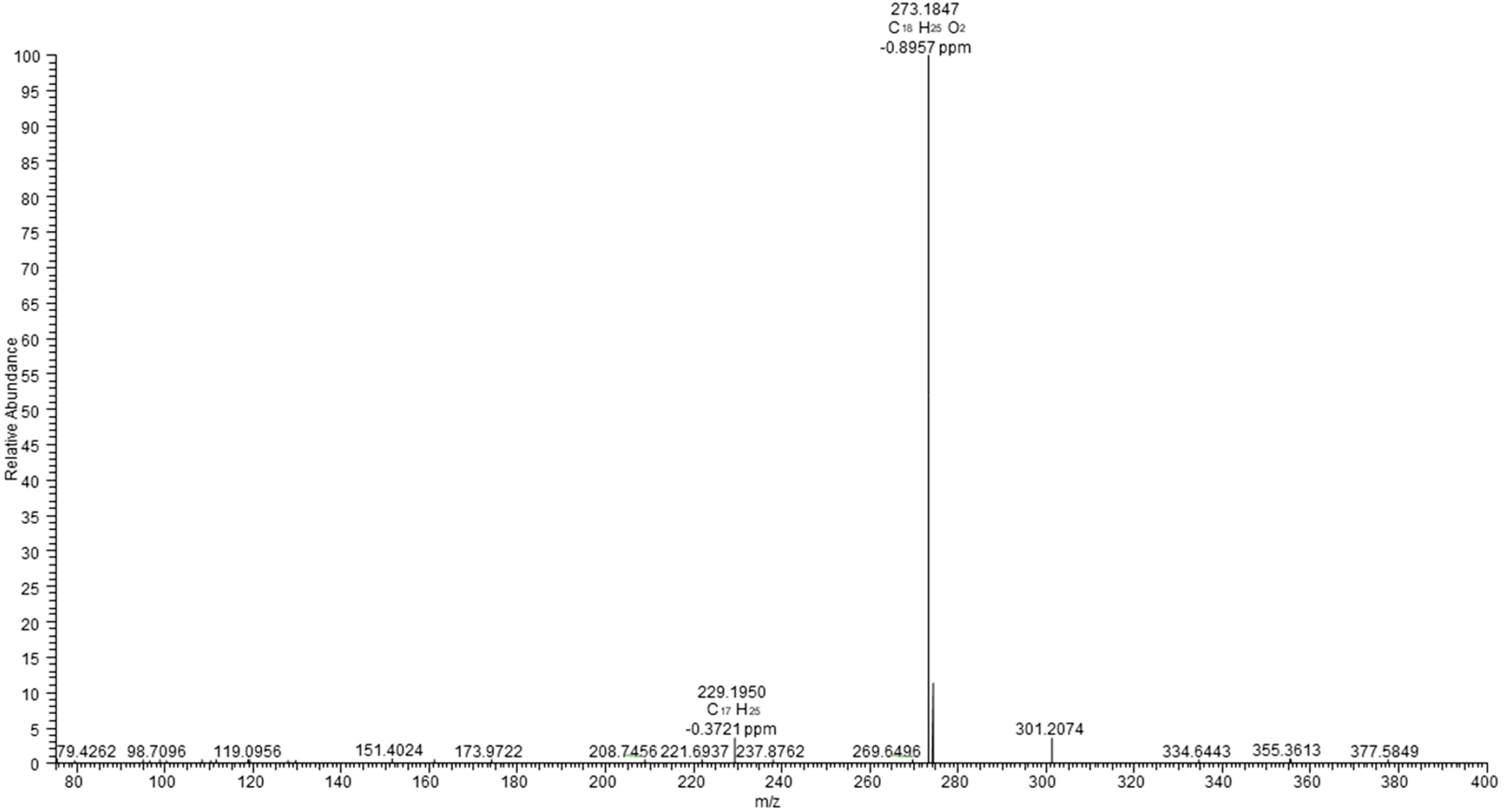
Tandem mass spectrum (MS3) of *m/z* 301.2 ± 1 → *m/z* 273.18 corresponding to the [M-H]^-^ ion of 121 (sandaracopimaric-18-acid) (*m/z* 301.2168, 1.66 ppm). The high resolution (HR) spectra were acquired with CID activation (normalized collision energy 25%) at the resolution of 15000 by a hybrid ESI-LIT-Orbitrap mass spectrometer (Orbitrap Elite, Thermo Fisher Scientific, Bremen, Germany) operated in negative ion mode, after on-line separation of peak 7813-3 by RP-UHPLC as described in the methods section.

**Fig. S32a.**
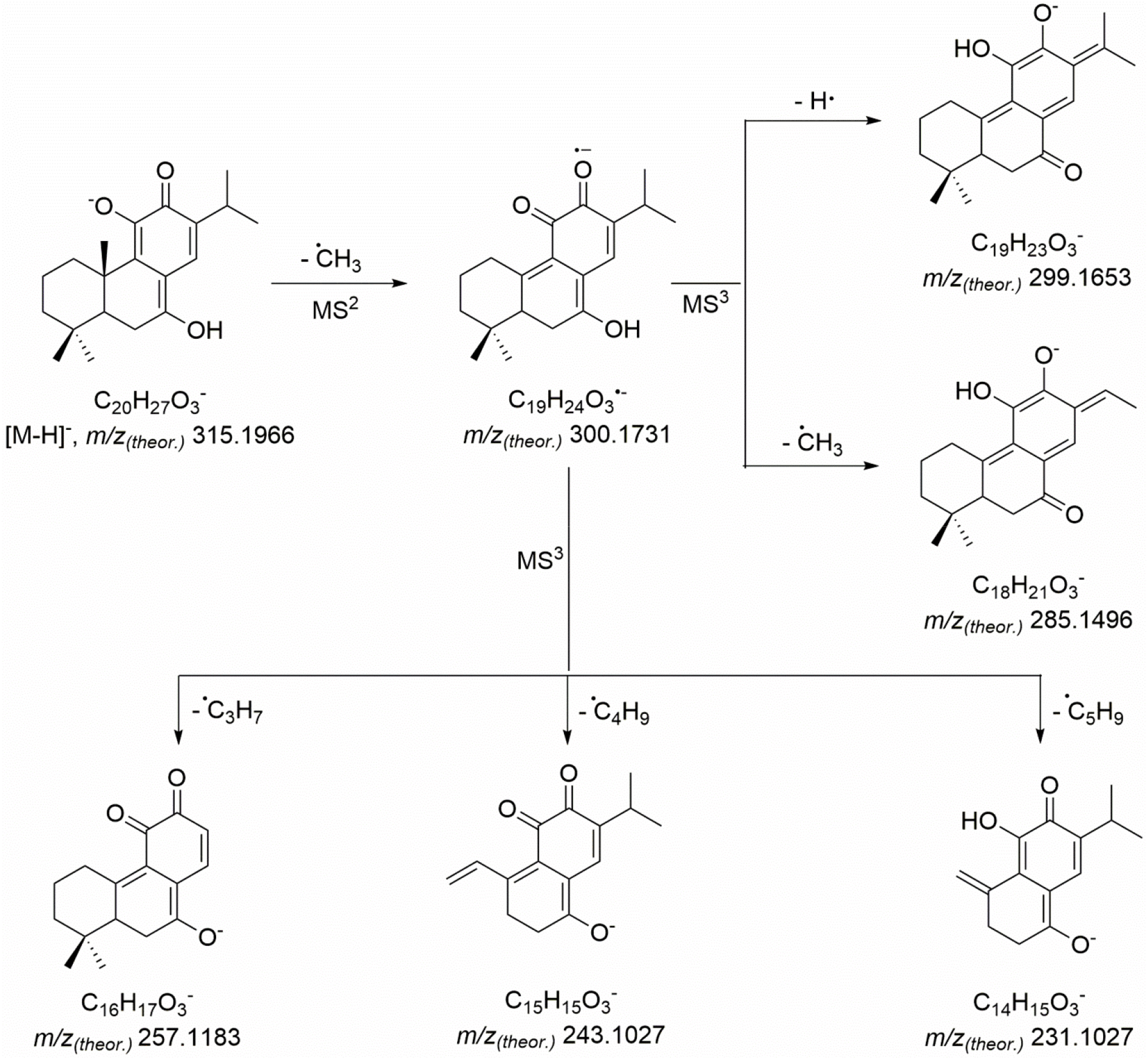
Tandem mass spectrometric fragmentation patterns (MS2, MS3) of 122 (7-hydroxy- 11,12-dioxo-abieta-7,13-diene, peak 7825-20)

**Fig. S32b.**
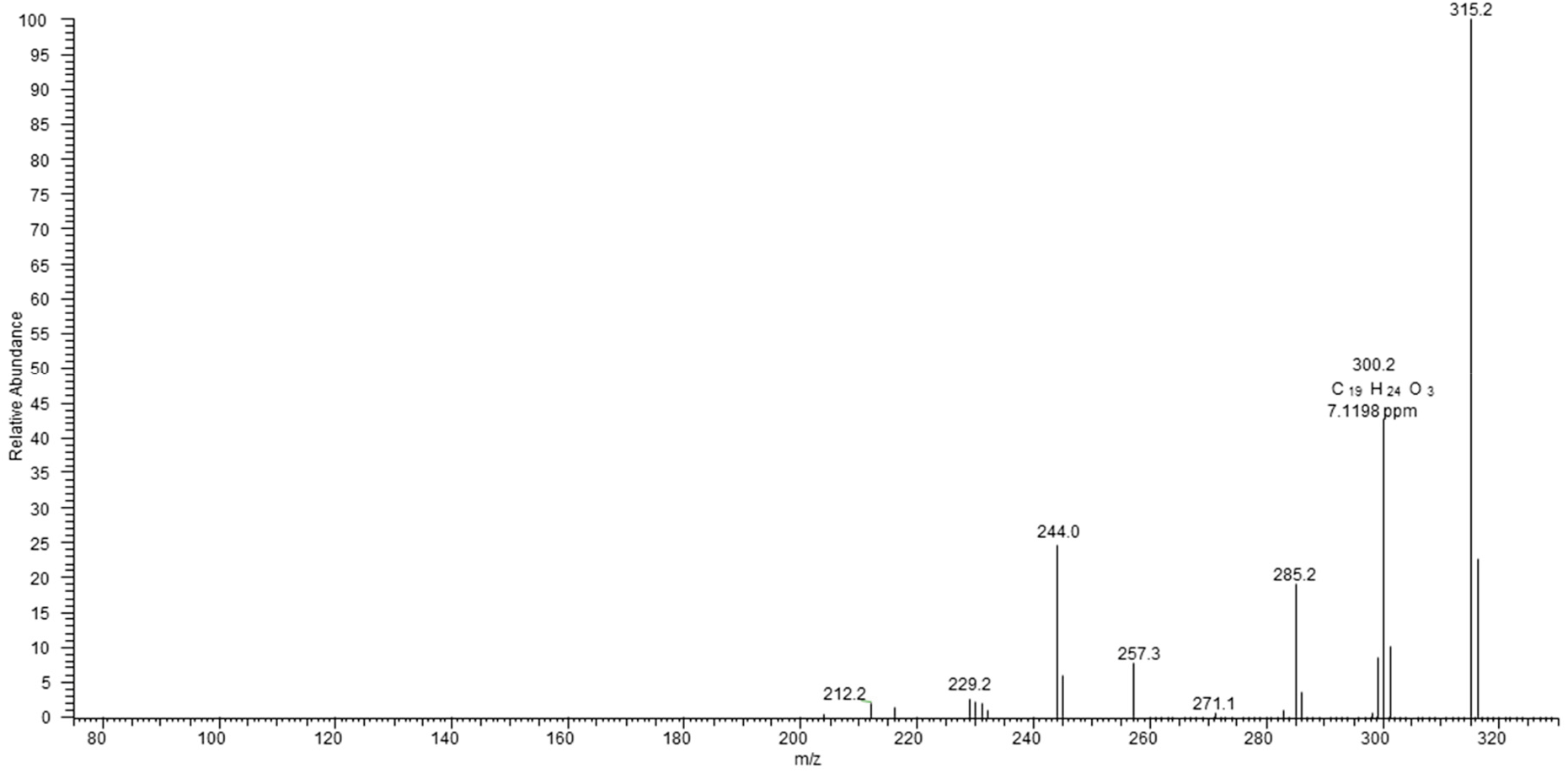
Tandem mass spectrum (MS2) of *m/z* 315.2 ± 1 corresponding to the [M-H]^-^ ion of 122 (7-hydroxy-11,12-dioxo-abieta-7,13-diene) (*m/z* 315.1966, 2.86 ppm) (part 1). The low resolution (LR) spectra were acquired with CID activation (normalized collision energy 35%) by a hybrid ESI- LIT-Orbitrap mass spectrometer (Orbitrap Elite, Thermo Fisher Scientific, Bremen, Germany) operated in negative ion mode, after on-line separation of peak 7825-20 by RP-UHPLC as described in the methods section.

**Fig. S32c.**
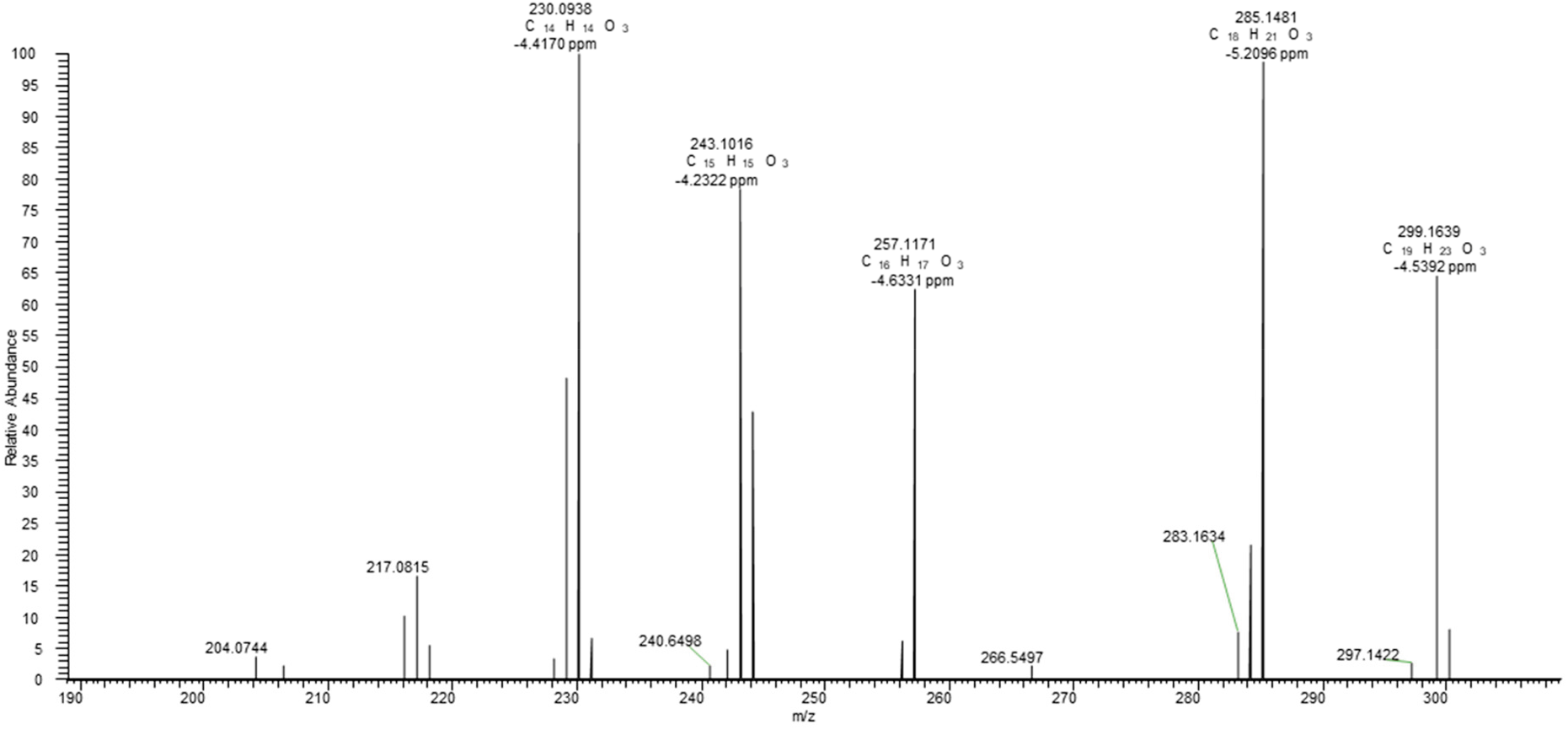
Tandem mass spectrum (MS3) of *m/z* 315.2 ± 1 → *m/z* 300.17 corresponding to the [M-H]^-^ ion of 122 (7-hydroxy-11,12-dioxo-abieta-7,13-diene) (*m/z* 301.2168, 1.66 ppm). The high resolution (HR) spectra were acquired with CID activation (normalized collision energy 35%) at the resolution of 15000 by a hybrid ESI-LIT-Orbitrap mass spectrometer (Orbitrap Elite, Thermo Fisher Scientific, Bremen, Germany) operated in negative ion mode, after on-line separation of peak 7825-20 by RP-UHPLC as described in the methods section.

**Fig. S33a.**
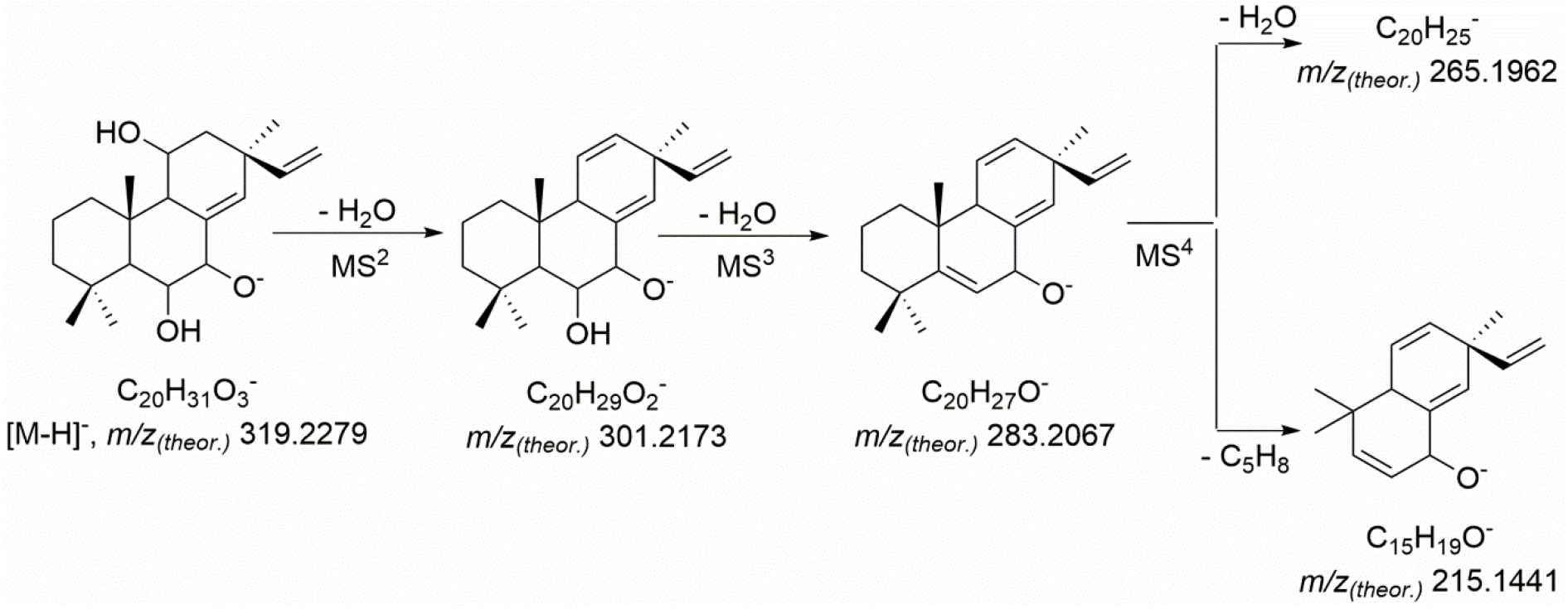
Advanced tandem mass spectrometric fragmentation patterns (MS2, MS3 and MS4) of 126 (6,7,11-trihydroxy pimara-8(14),15-diene, peak 7870-4).

**Fig. S33b.**
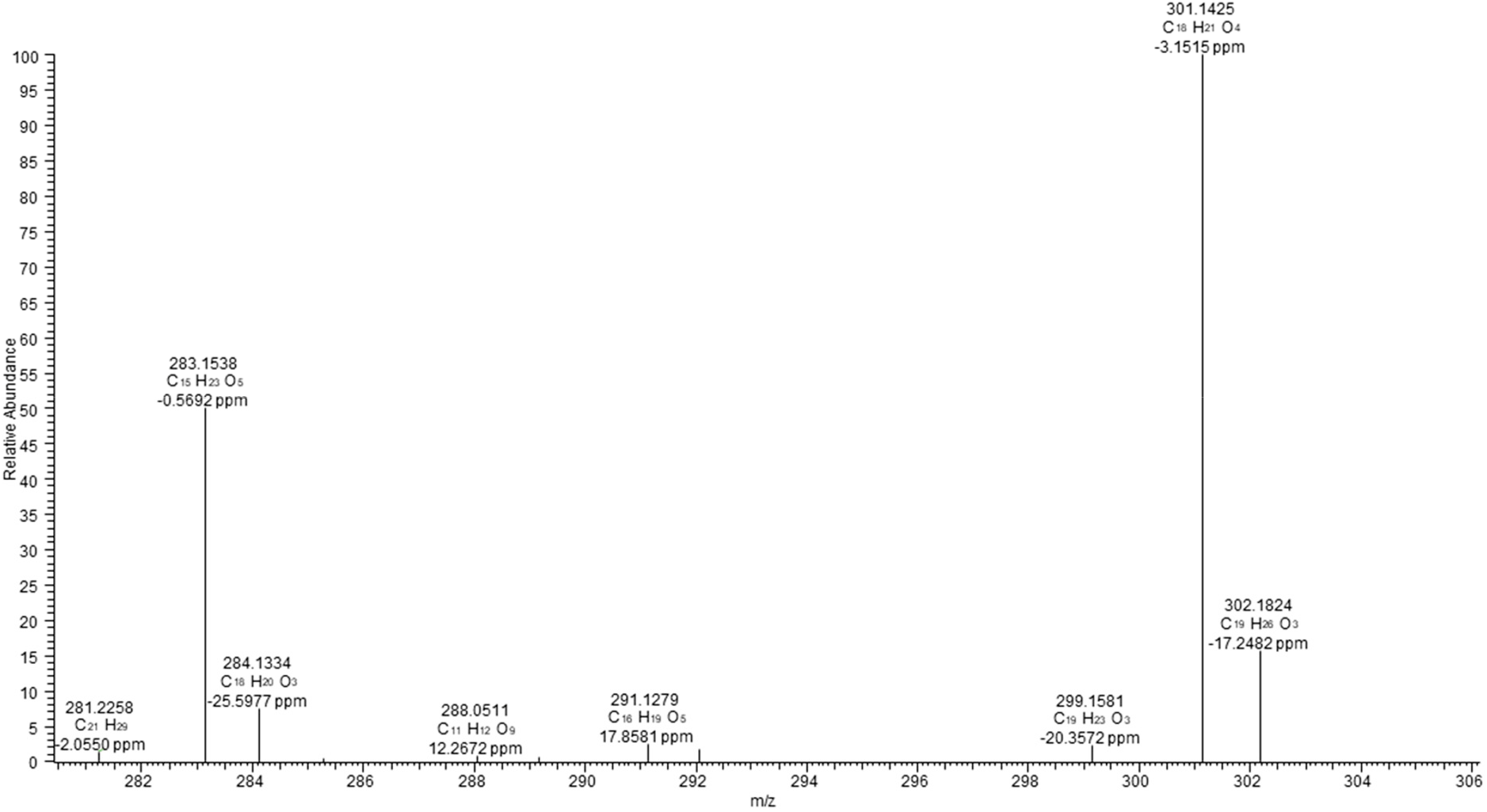
Tandem mass spectrum (MS2) of *m/z* 319.2 ± 1 corresponding to the [M-H]^-^ ion of 126 (6,7,11-trihydroxy pimara-8(14),15-diene) (*m/z* 319.2273, 1.88 ppm). The high resolution (HR) spectra were acquired with CID activation (normalized collision energy 35%) at the resolution of 15000 by a hybrid ESI-LIT-Orbitrap mass spectrometer (Orbitrap Elite, Thermo Fisher Scientific, Bremen, Germany) operated in negative ion mode, after on-line separation of the peak 7970-4 by RP-UHPLC as described in the Methods section.

**Fig. S33c.**
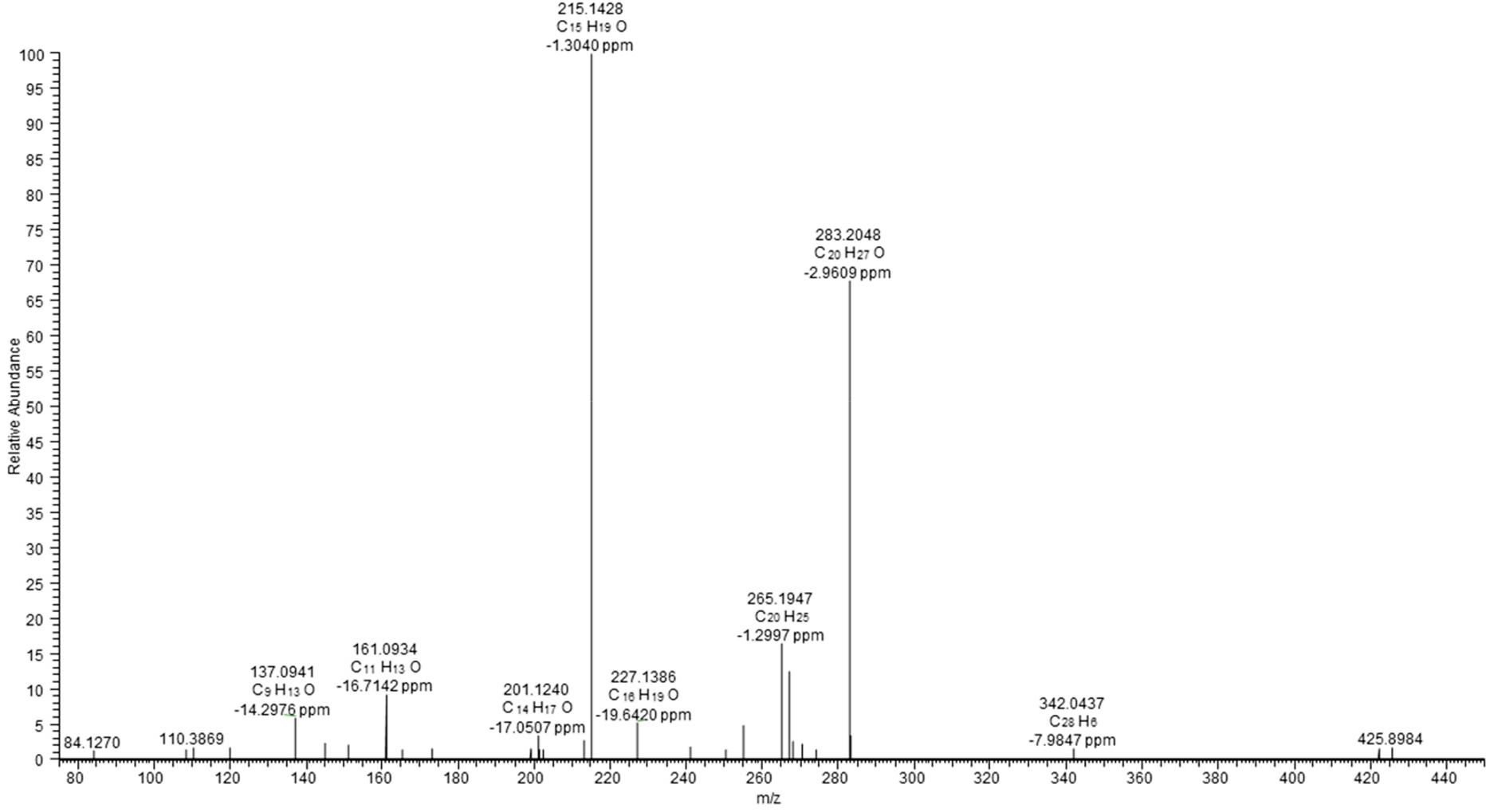
Tandem mass spectrum (MS3) of *m/z* 319.2 ± 1 → *m/z* 283.15 corresponding to the [M-H]^-^ ion of 126 (6,7,11-trihydroxy pimara-8(14),15-diene) (*m/z* 319.2273, 1.88 ppm). The high resolution (HR) spectra were acquired with CID activation (normalized collision energy 30%) at the resolution of 15000 by a hybrid ESI-LIT-Orbitrap mass spectrometer (Orbitrap Elite, Thermo Fisher Scientific, Bremen, Germany) operated in negative ion mode, after on-line separation of peak 7970-4 by RP-UHPLC as described in the Methods section.

**Fig. S33d.**
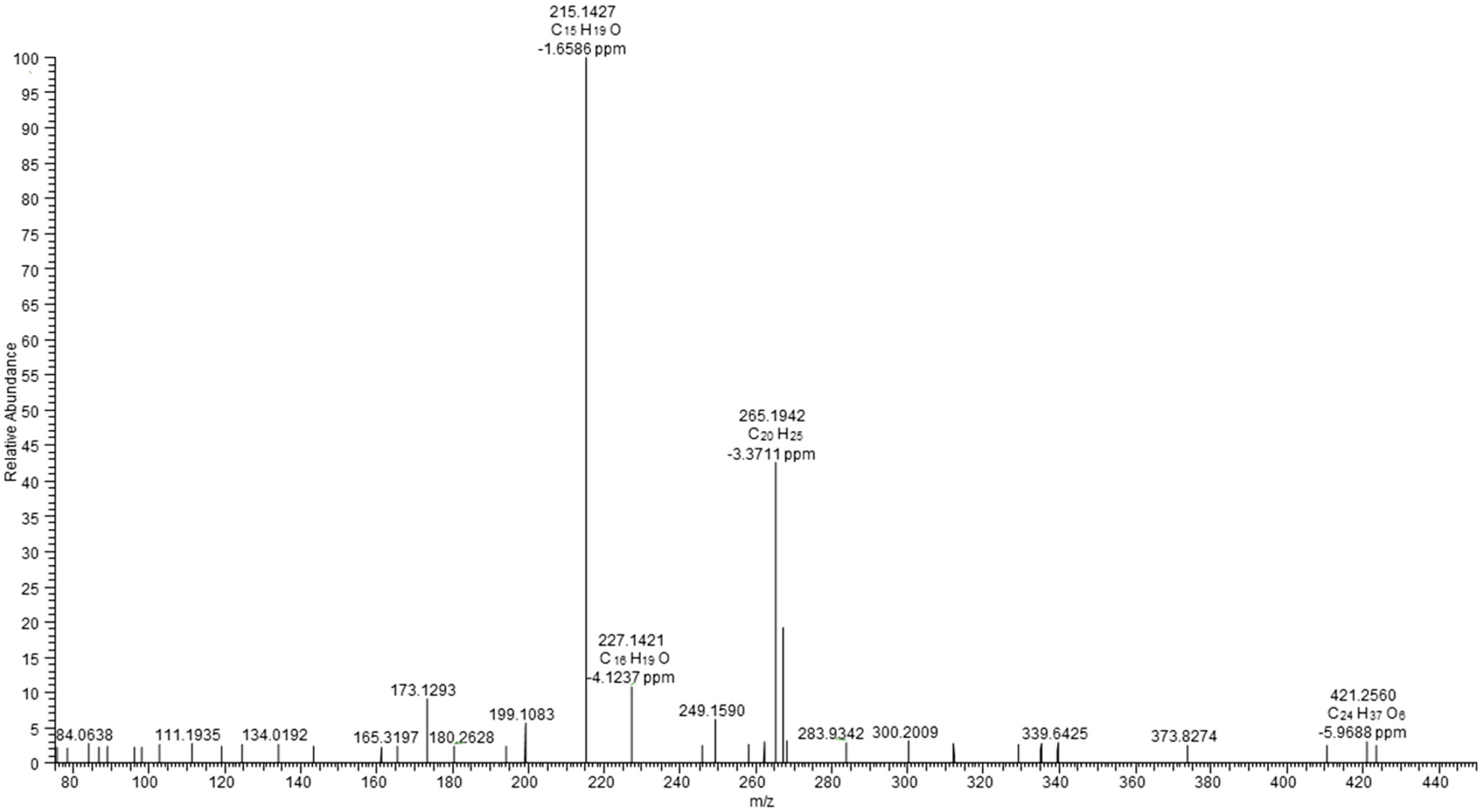
Tandem mass spectrum (MS4) of *m/z* 319.2 ± 1 → *m/z* 301.16 → *m/z* 283.20 corresponding to the [M-H]^-^ ion of 126 (6,7,11-trihydroxy pimara-8(14),15-diene) (*m/z* 319.2273, 1.88 ppm). The high resolution (HR) spectra were acquired with CID activation (normalized collision energy 40%) at the resolution of 15000 by a hybrid ESI-LIT-Orbitrap mass spectrometer (Orbitrap Elite, Thermo Fisher Scientific, Bremen, Germany) operated in negative ion mode, after on-line separation of peak 7970-4 by RP-UHPLC as described in the Methods section.

**Fig. S34a.**
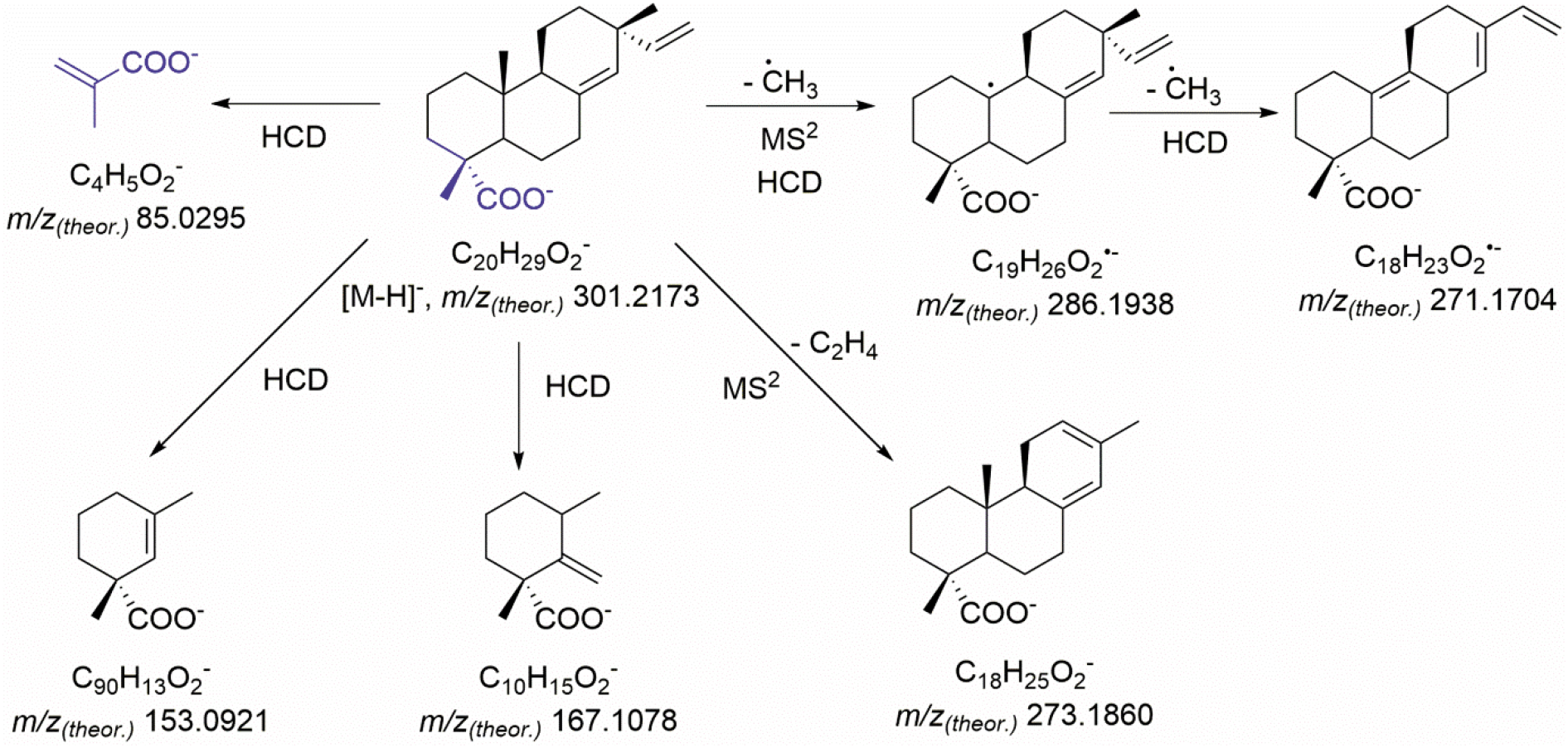
Tandem mass spectrometric fragmentation patterns (MS2) of 127 (pimaric-18-acid, peak 7971-4).

**Figure S34b.**
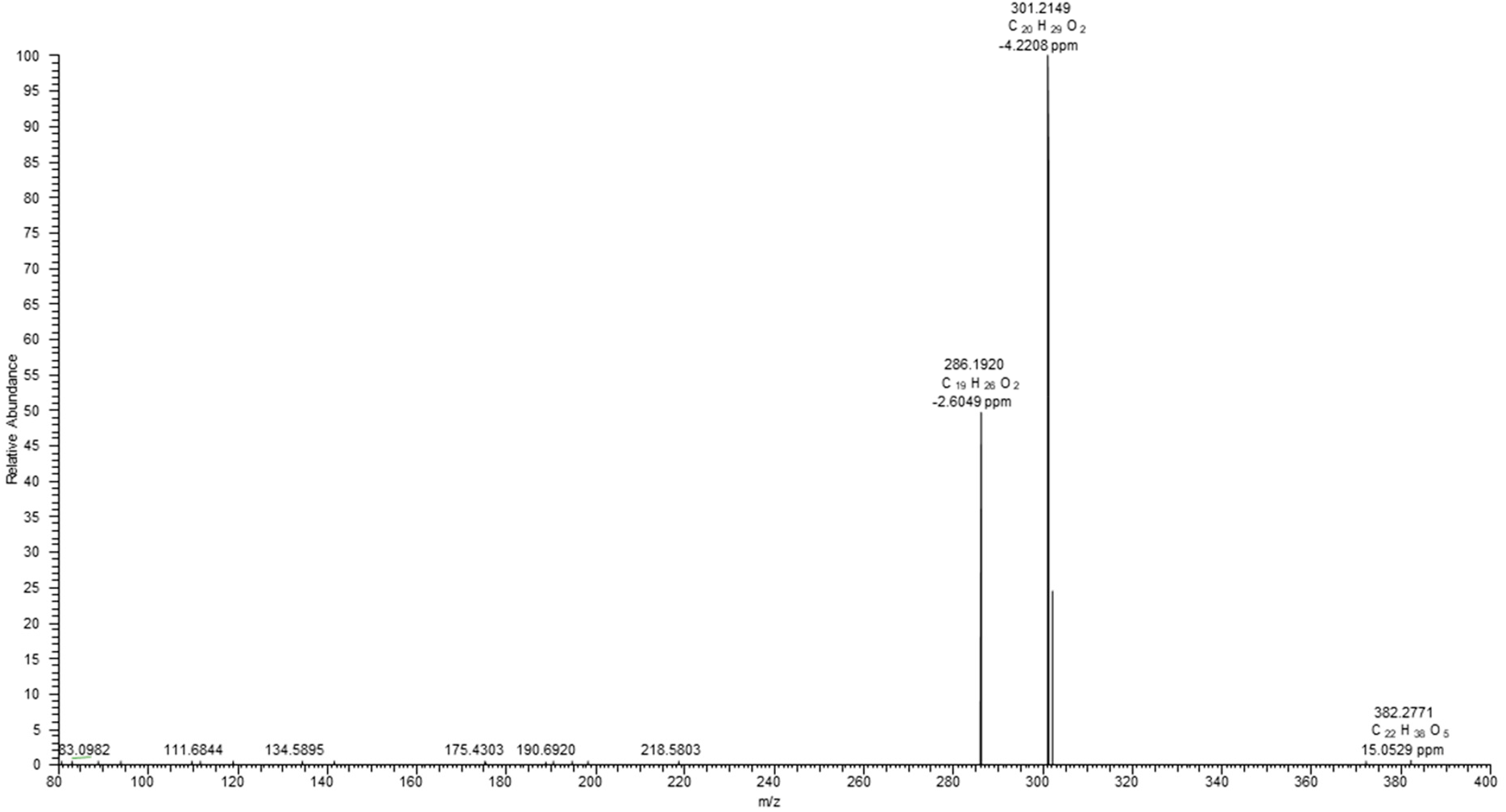
Tandem mass spectrum (MS2) of *m/z* 301.2 ± 1 corresponding to the [M-H]^-^ ion of 127 (pimaric-18-acid) (*m/z* 301.2167, 1.99 ppm). The high resolution (HR) spectra were acquired with CID activation (normalized collision energy 30%, activation time 20 ms) at the resolution of 15000 by a hybrid ESI-LIT-Orbitrap mass spectrometer (Orbitrap Elite, Thermo Fisher Scientific, Bremen, Germany) operated in negative ion mode, after on-line separation of the sample 7971-4 by RP-UHPLC as described in Methods section.

**Fig. S34c.**
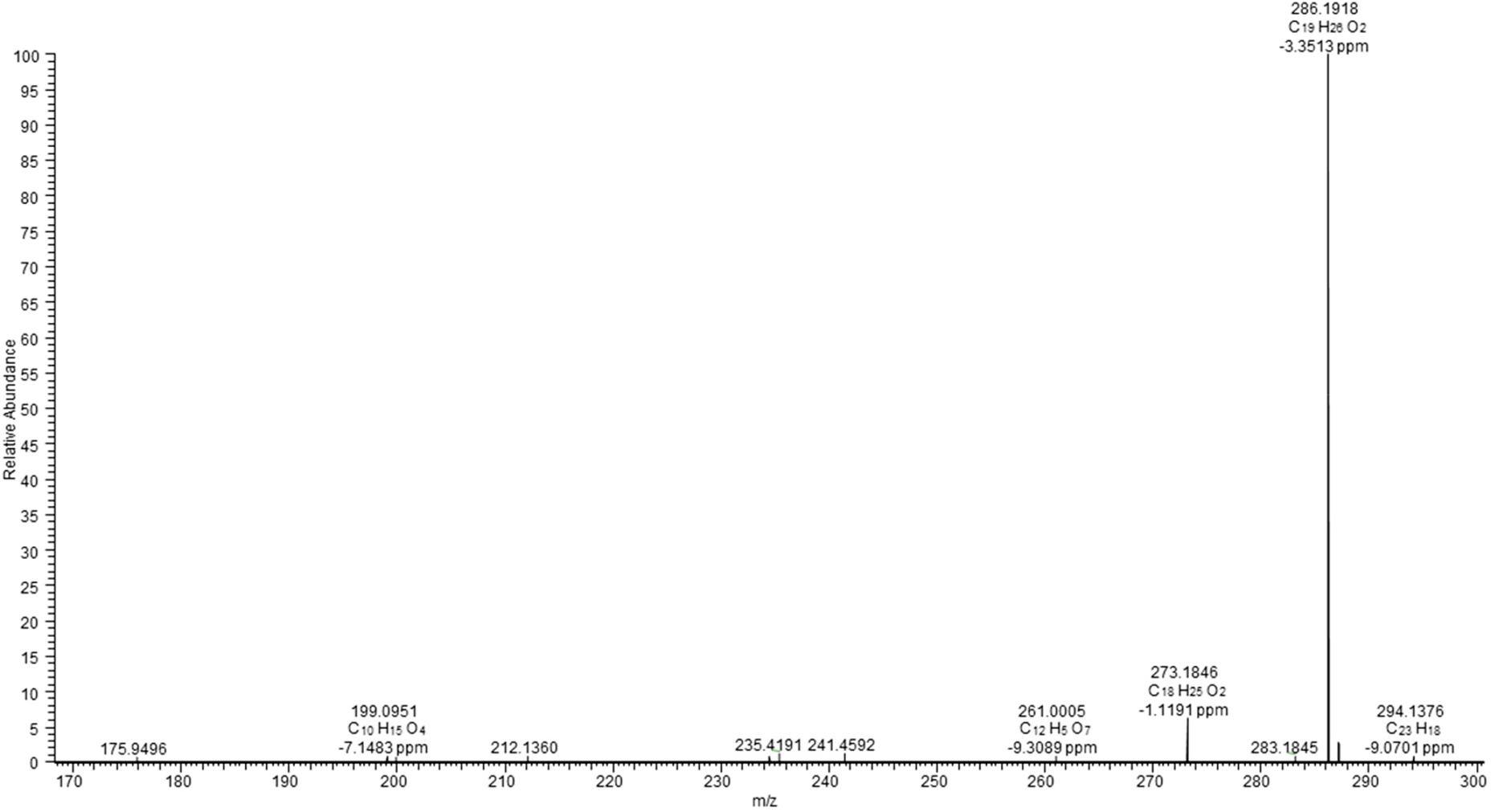
Tandem mass spectrum (MS2) of *m/z* 301.2 ± 1 corresponding to the [M-H]^-^ ion of 127 (pimaric-18-acid) (*m/z* 301.2167, 1.99 ppm). The high resolution (HR) spectra were acquired with CID activation (normalized collision energy 35%, activation time 200 ms) at the resolution of 15000 by a hybrid ESI-LIT-Orbitrap mass spectrometer (Orbitrap Elite, Thermo Fisher Scientific, Bremen, Germany) operated in negative ion mode, after on-line separation of peak 7971-4 by RP-UHPLC as described in the Methods section.

**Fig. S34d.**
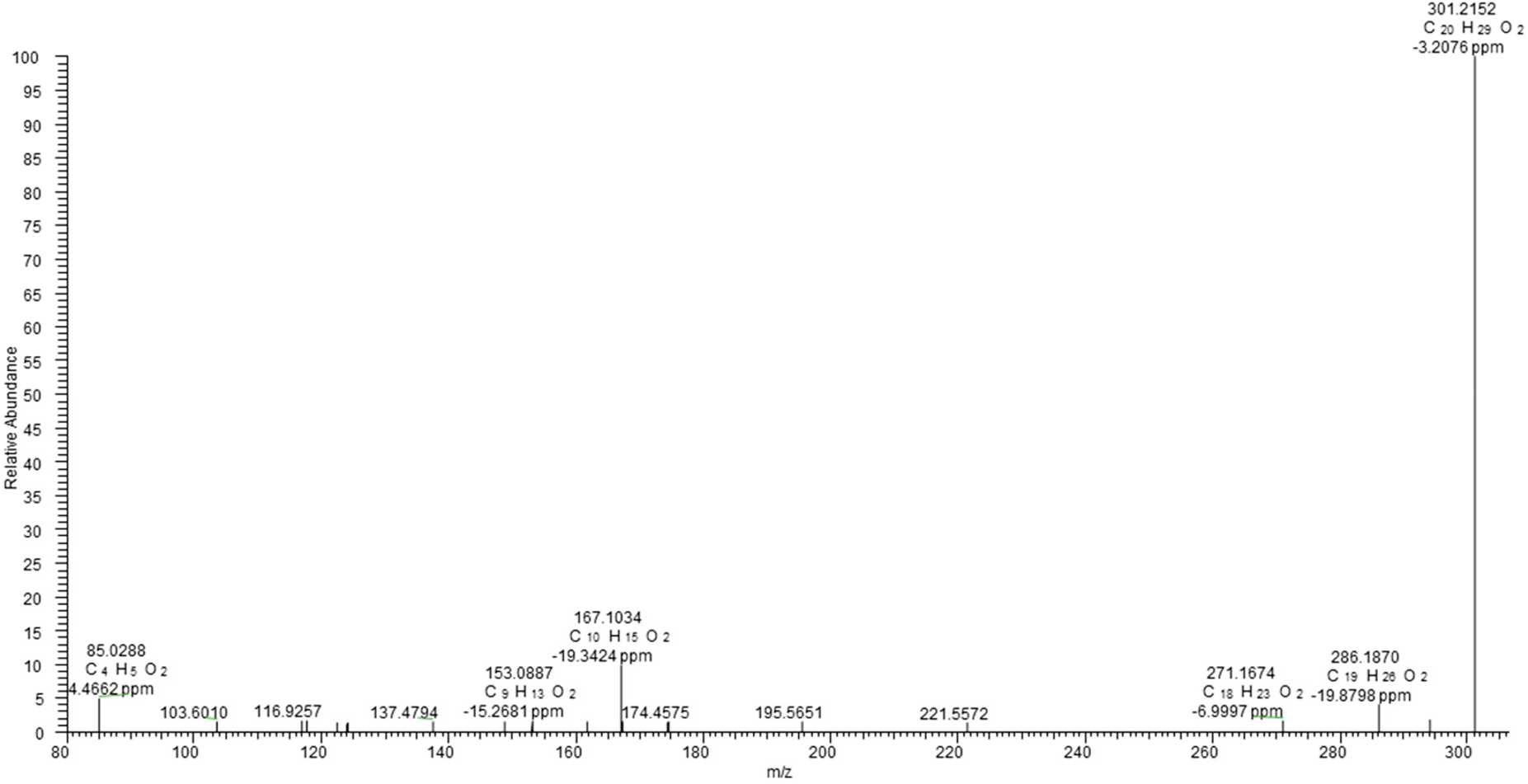
Tandem mass spectrum (MS2) of *m/z* 301.2 ± 1 corresponding to the [M-H]^-^ ion of 127 (pimaric-18-acid) (*m/z* 301.2167, 1.99 ppm). The high resolution (HR) spectra were acquired with HCD activation (160 eV) at the resolution of 15000 by a hybrid ESI-LIT-Orbitrap mass spectrometer (Orbitrap Elite, Thermo Fisher Scientific, Bremen, Germany) operated in negative ion mode, after on- line separation of peak 7971-4 by RP-UHPLC as described in the Methods section.

**Fig. S35a.**
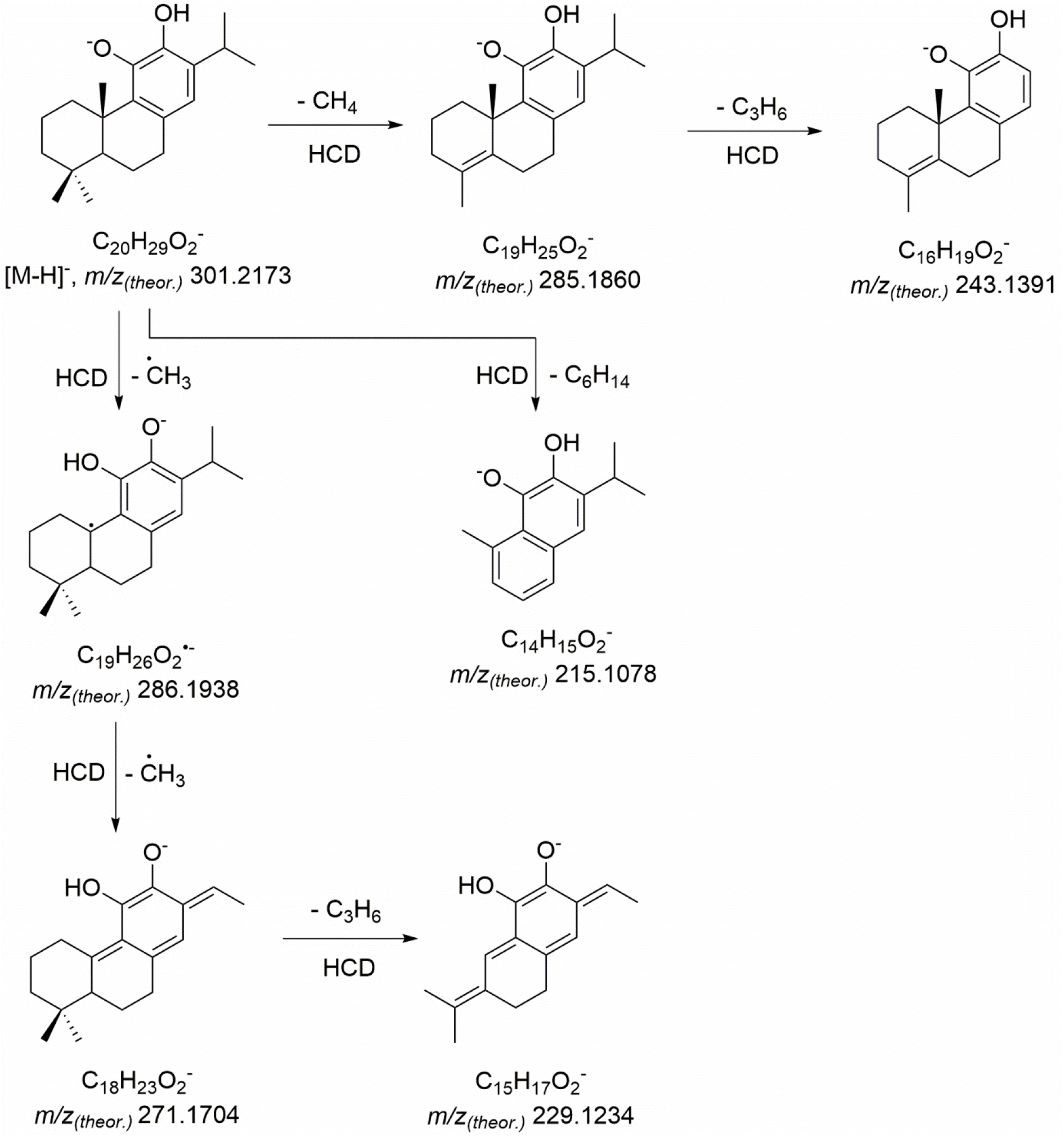
Advanced tandem mass spectrometric fragmentation patterns (MS2) of 6 (11- hydroxyferruginol, peak 7973-13).

**Fig. S35b.**
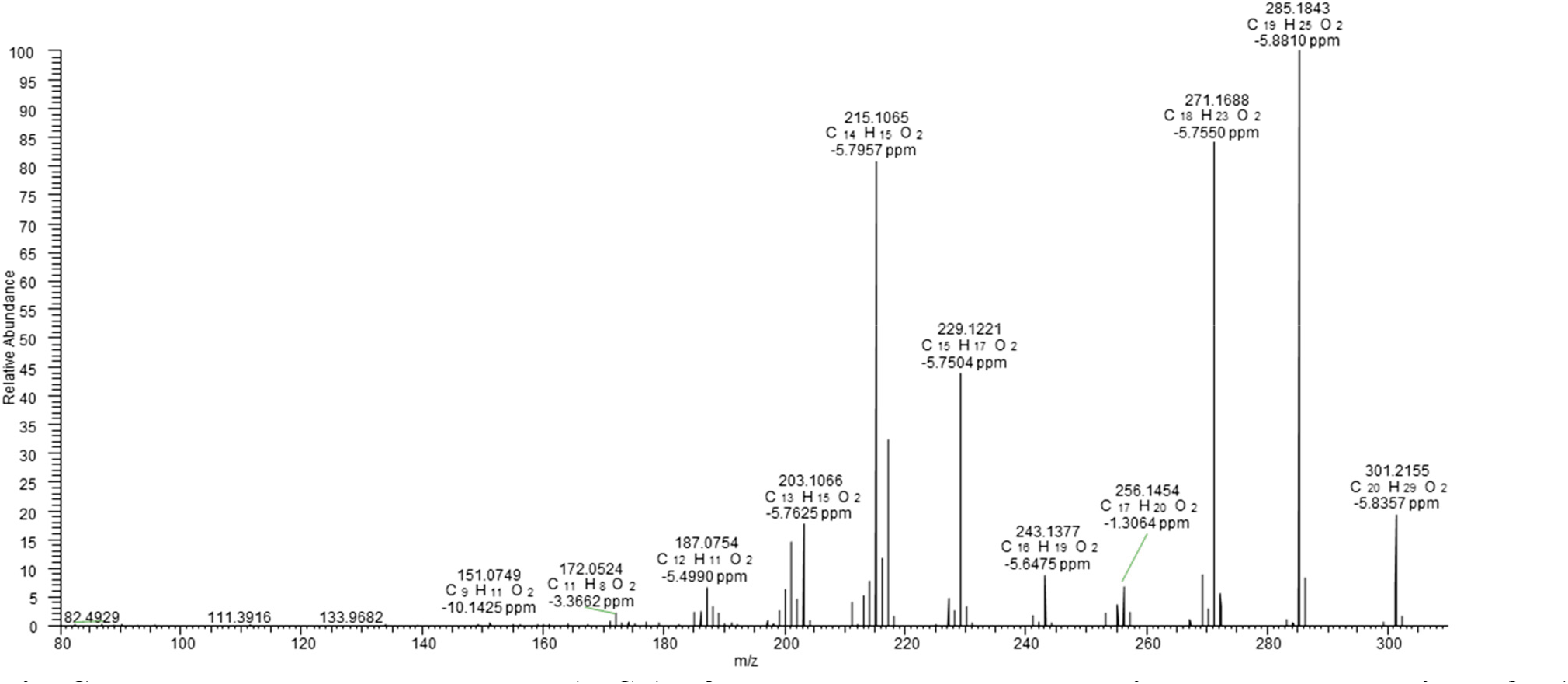
Tandem mass spectrum (MS2) of *m/z* 301.2 ± 1 corresponding to the [M-H]^-^ ion of 6 (11- hydroxyferruginol) (*m/z* 301.2165, 2.66 ppm). The high resolution (HR) spectra were acquired with HCD activation (200 eV) at the resolution of 15000 by a hybrid ESI-LIT-Orbitrap mass spectrometer (Orbitrap Elite, Thermo Fisher Scientific, Bremen, Germany) operated in negative ion mode, after on- line separation of peak 7973-13 by RP-UHPLC as described in the Methods section.

**Fig. S35c.**
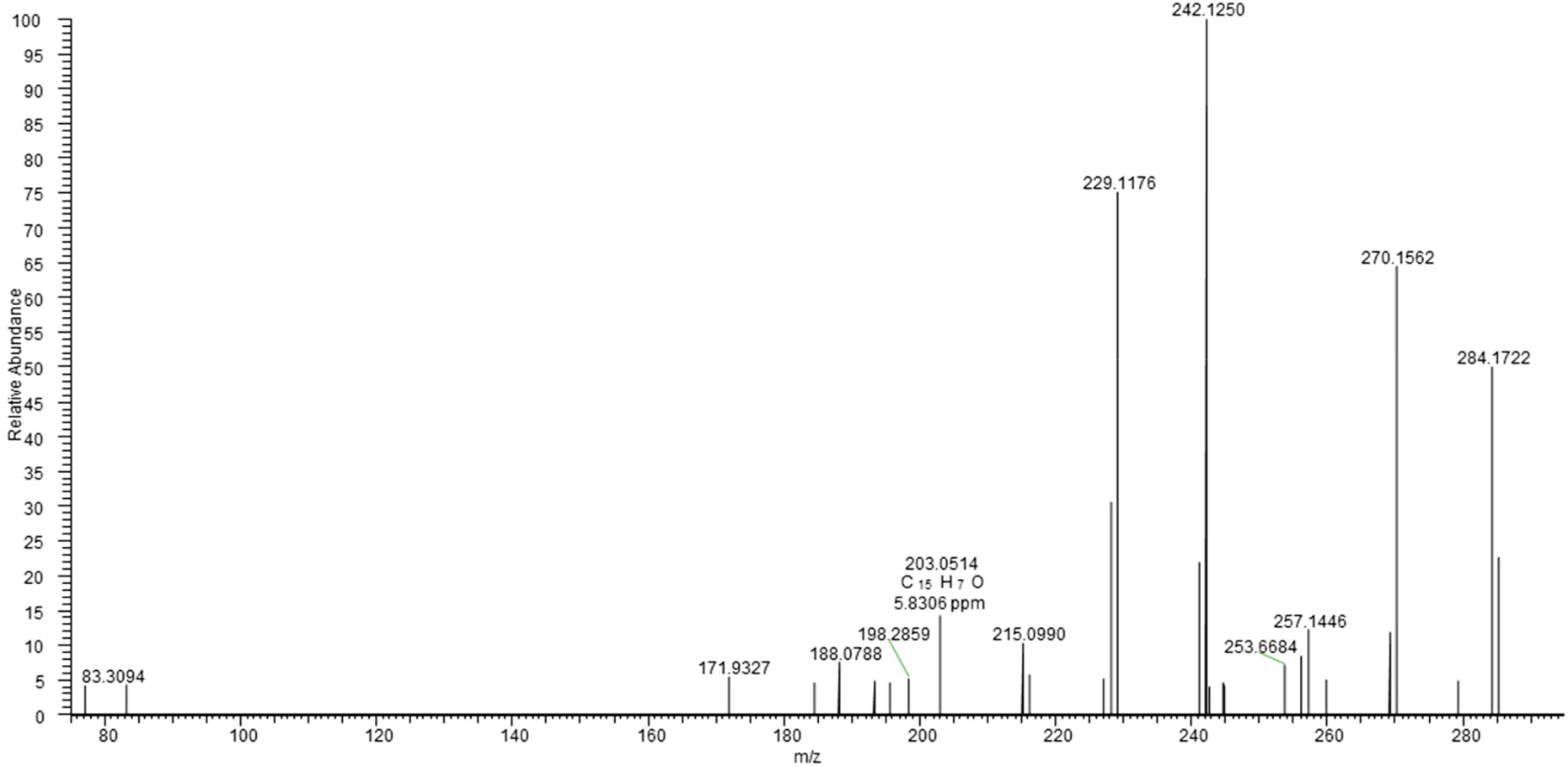
Tandem mass spectrum (MS3) of *m/z* 301.2 ± 1 → *m/z* 285.2 corresponding to the [M-H]^-^ ion of 6 (11-hydroxyferruginol) (*m/z* 301.2165, 2.66 ppm). The high resolution (HR) spectra were acquired with CID activation (normalized collision energy 35%) at the resolution of 15000 by a hybrid ESI-LIT-Orbitrap mass spectrometer (Orbitrap Elite, Thermo Fisher Scientific, Bremen, Germany) operated in negative ion mode, after on-line separation of peak 7973-13 by RP-UHPLC as described in Methods section.

**Fig. S35d.**
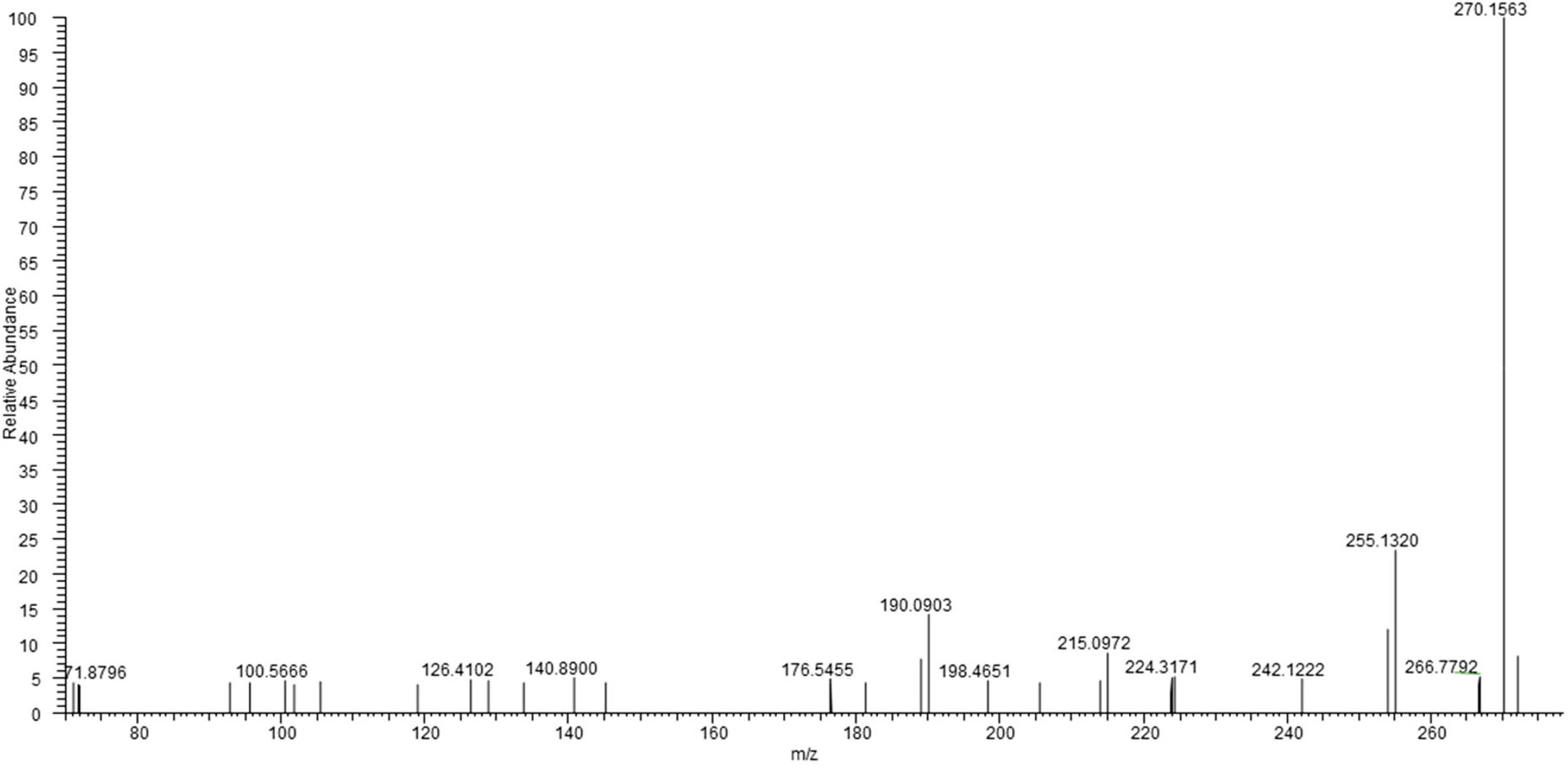
Tandem mass spectrum (MS3) of *m/z* 301.2 ± 1 → *m/z* 271.2 corresponding to the [M-H]^-^ ion of 6 (11-hydroxyferruginol) (*m/z* 301.2165, 2.66 ppm). The high resolution (HR) spectra were acquired with CID activation (normalized collision energy 35%) at the resolution of 15000 by a hybrid ESI-LIT-Orbitrap mass spectrometer (Orbitrap Elite, Thermo Fisher Scientific, Bremen, Germany) operated in negative ion mode, after on-line separation of peak 7973-13 by RP-UHPLC as described in the Methods section.

**Fig. S36a.**
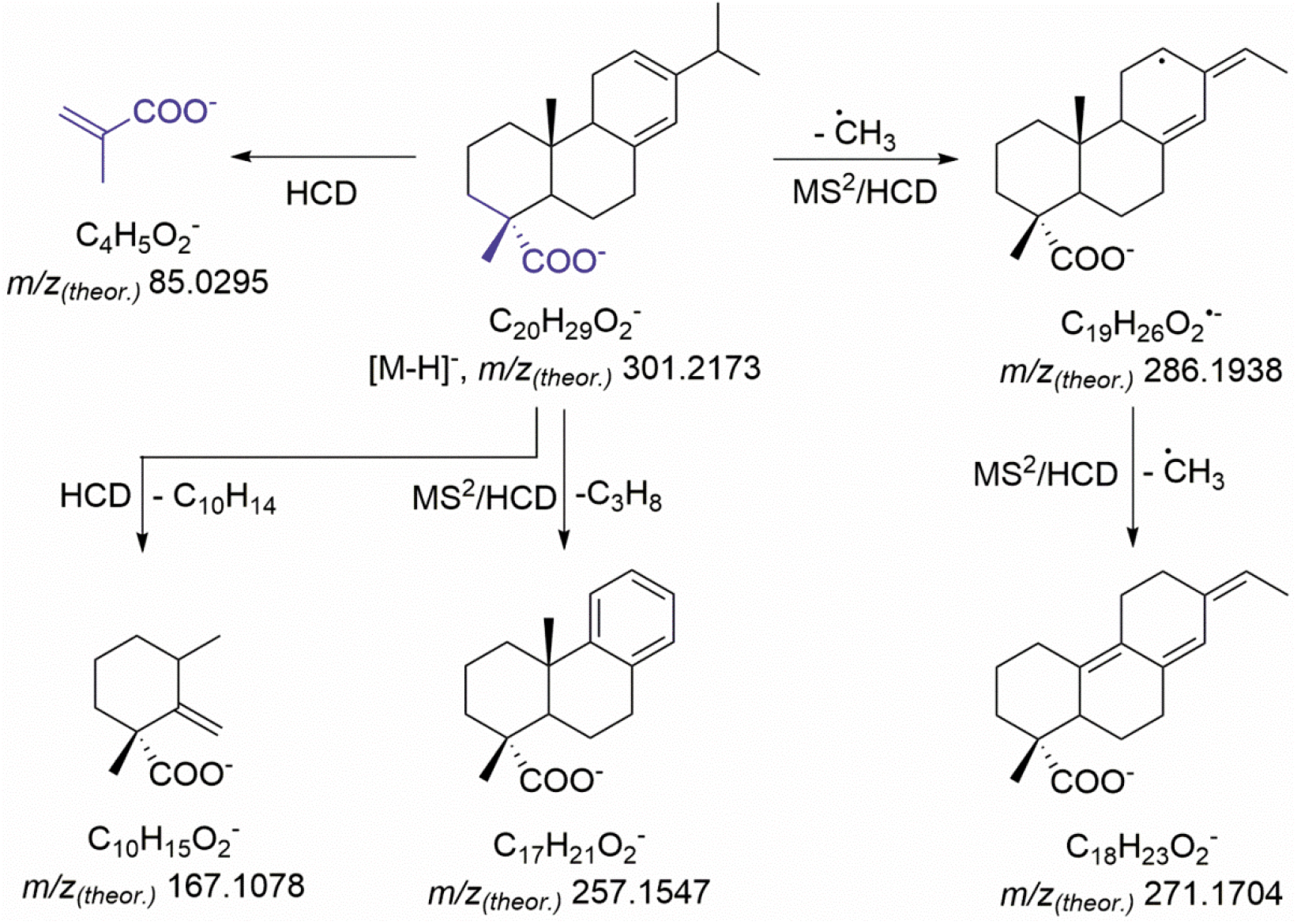
Tandem mass spectrometric fragmentation patterns (MS2) of 128 (levopimaric-18-acid, peak 7976-9).

**Fig. S36b.**
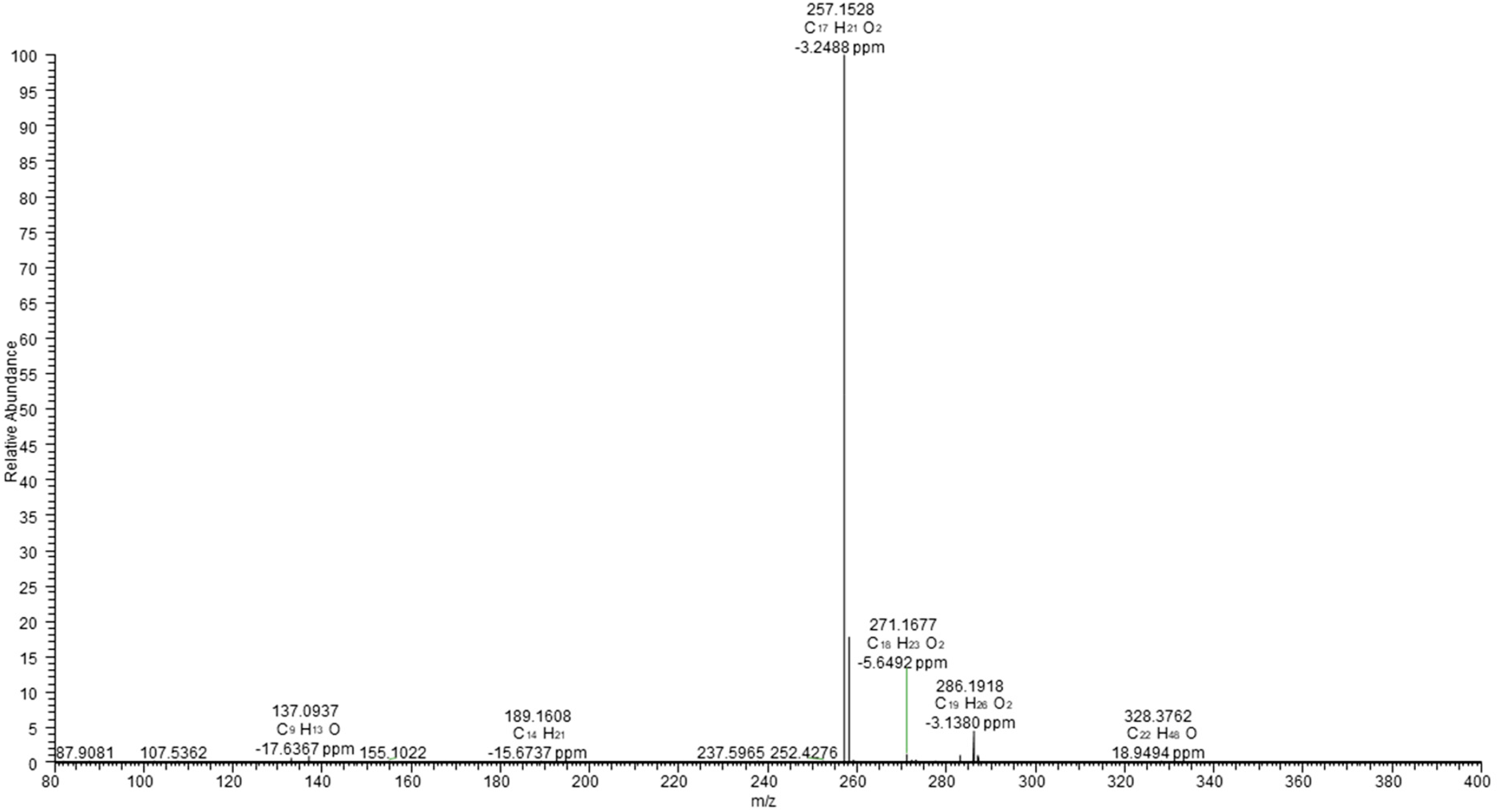
Tandem mass spectrum (MS2) of *m/z* 301.2 ± 1 corresponding to the [M-H]^-^ ion of 128 (levopimaric-18-acid) (*m/z* 301.2165, 2.66 ppm). The high resolution (HR) spectra were acquired with CID activation (normalized collision energy 45%) at the resolution of 15000 by a hybrid ESI-LIT- Orbitrap mass spectrometer (Orbitrap Elite, Thermo Fisher Scientific, Bremen, Germany) operated in negative ion mode, after on-line separation of peak 7976-9 by RP-UHPLC as described in the Methods section.

**Fig. S36c.**
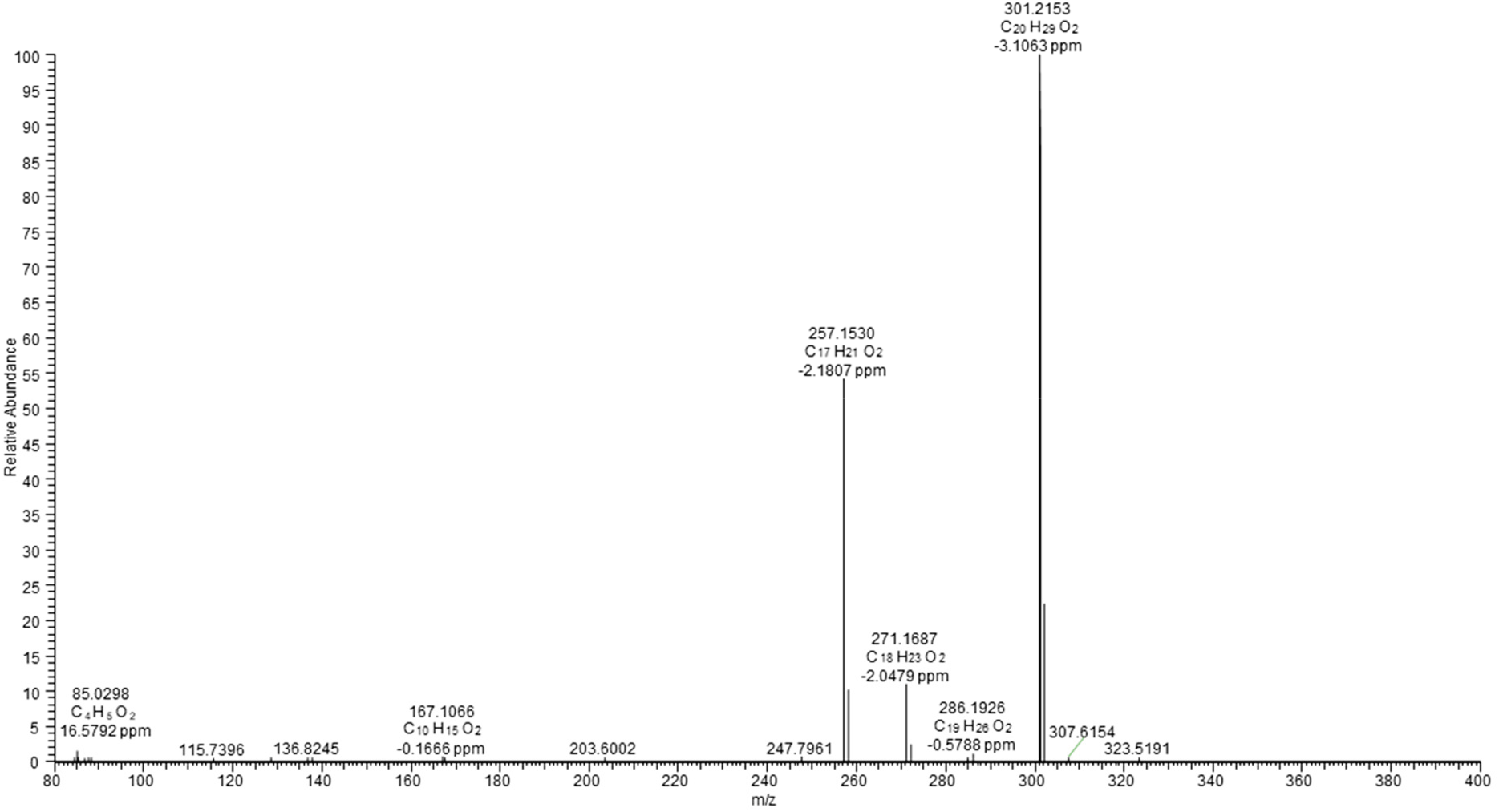
Tandem mass spectrum (MS2) of *m/z* 301.2 ± 1 corresponding to the [M-H]- ion of 128 (levopimaric-18-acid) (*m/z* 301.2165, 2.66 ppm). The high resolution (HR) spectra were acquired with HCD activation (160 eV) at the resolution of 15000 by a hybrid ESI-LIT-Orbitrap mass spectrometer (Orbitrap Elite, Thermo Fisher Scientific, Bremen, Germany) operated in negative ion mode, after on- line separation of peak 7976-9 by RP-UHPLC as described in the methods section.

**Fig. S37a.**
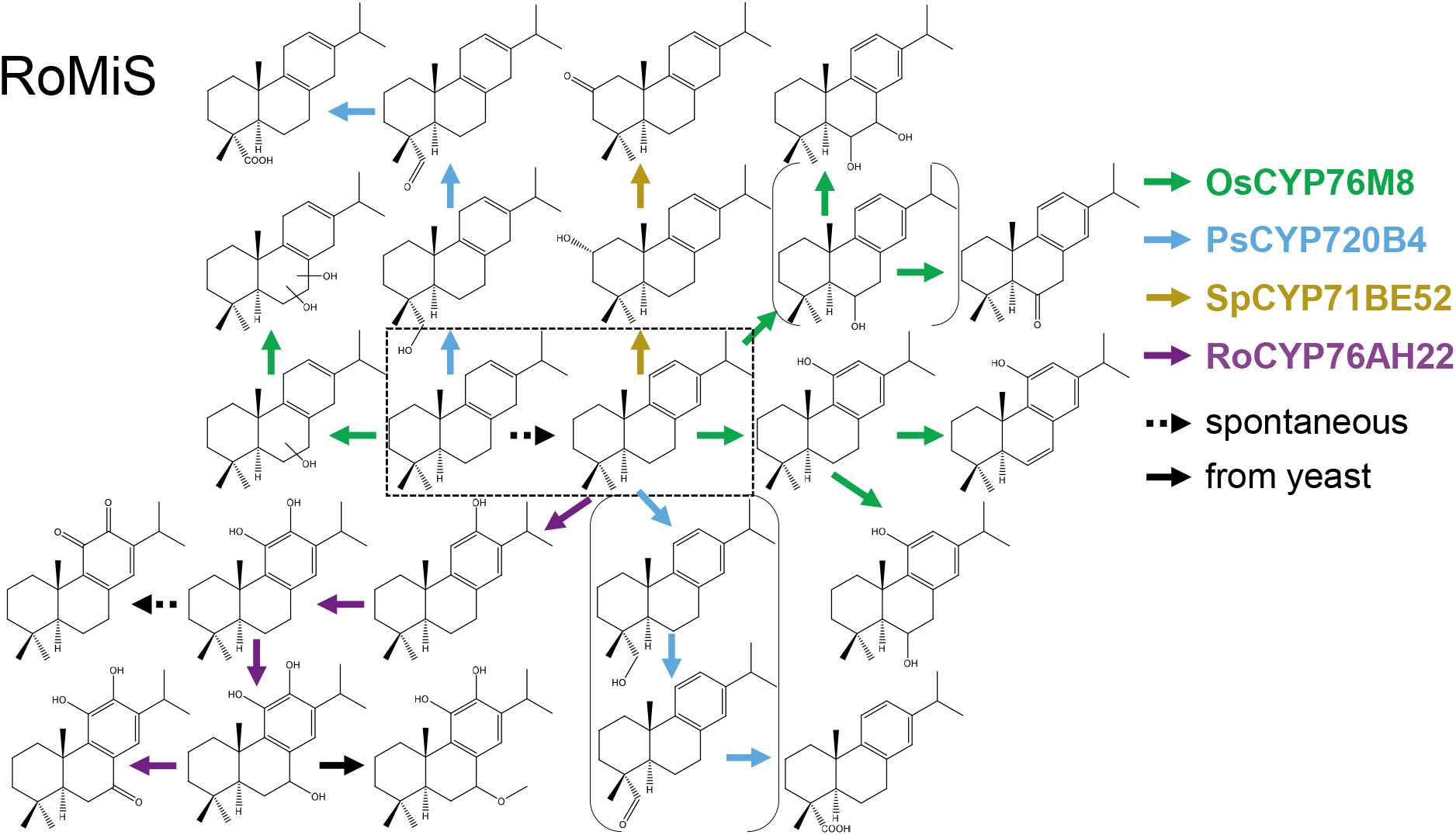
Overview of the network of products derived from miltiradiene (produced by RoMiS). Miltiradiene is on the left side of the box with dashed lines. Abietatriene, on the right side of that box, is a spontaneous oxidation product of miltiradiene. Reactions catalyzed by the CYP enzymes are indicated by the color code on the right side of the figure.

**Fig. S37b.**
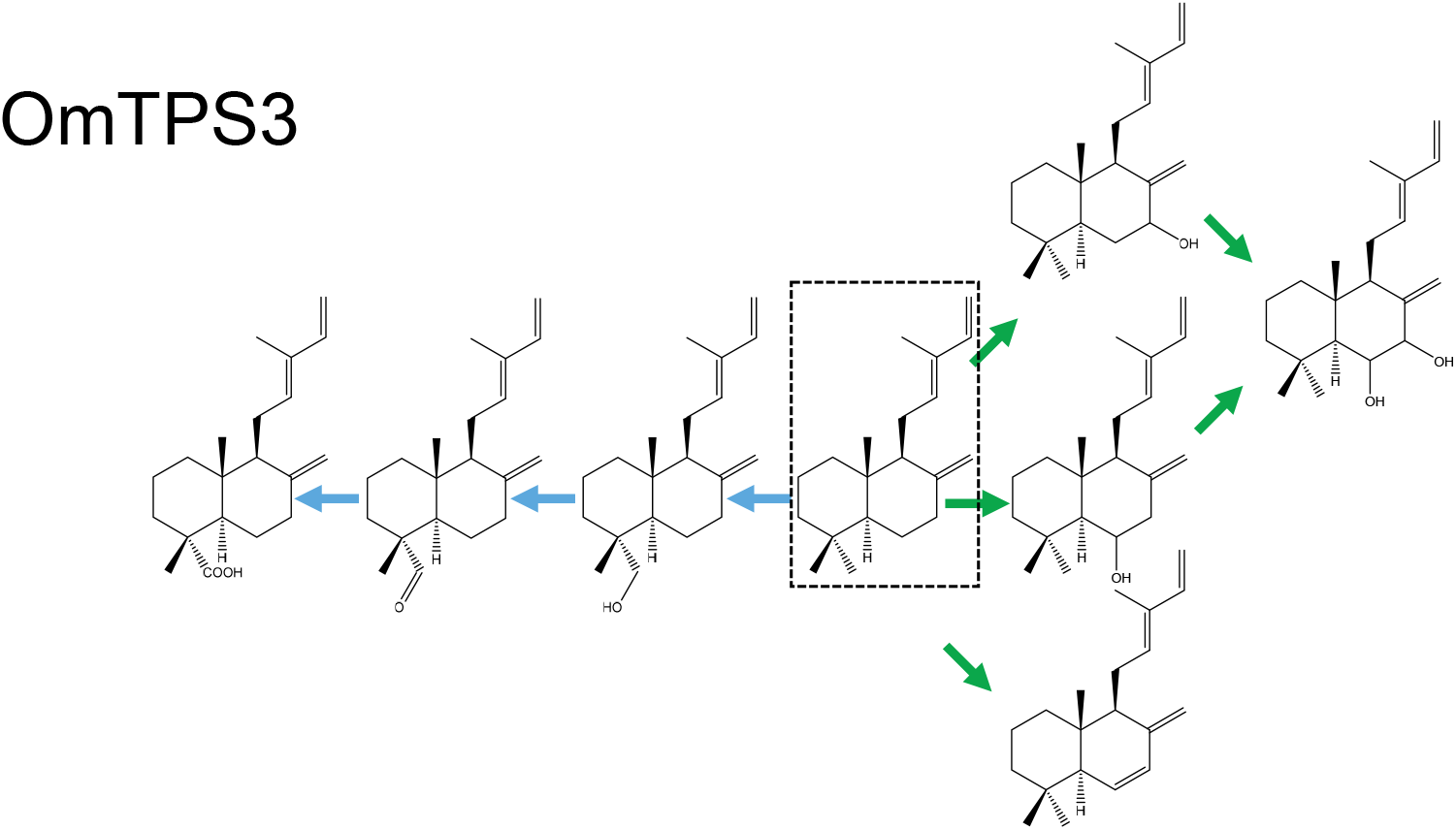
Overview of the network of products derived from trans-biformene (produced by OmTPS3, in the box with dashed lines). Reactions catalyzed by the CYP enzymes are indicated by the color code as in Fig. S37a.

**Fig. S37c.**
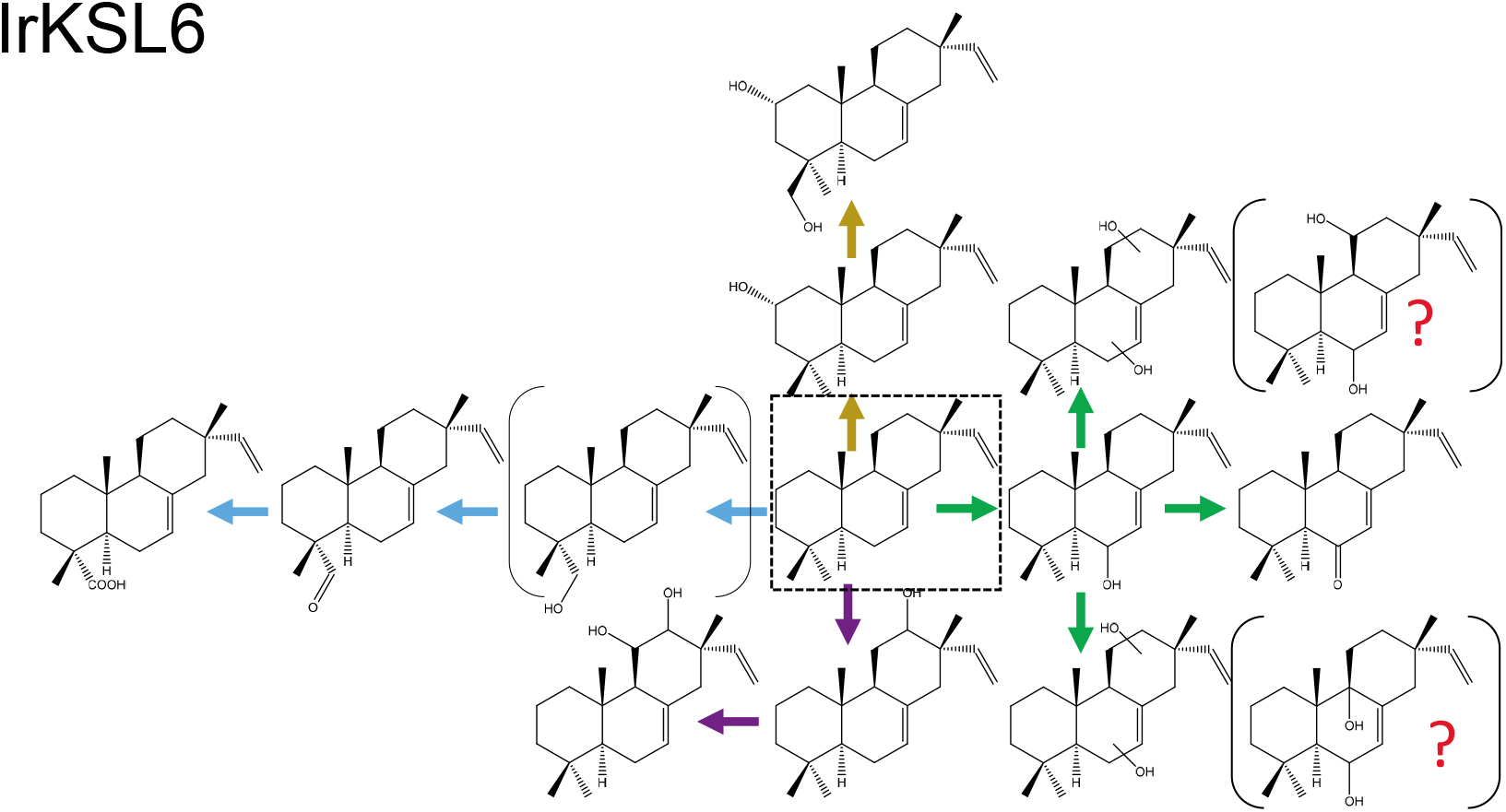
Overview of the network of products derived from isopimara-7,15-diene (produced by IrKSL6, in the box with dashed lines). Reactions catalyzed by the CYP enzymes are indicated by the color code as in Fig. S37a. Question marks indicate hypothetical structures for the compounds on located on their left.

**Fig. S37d.**
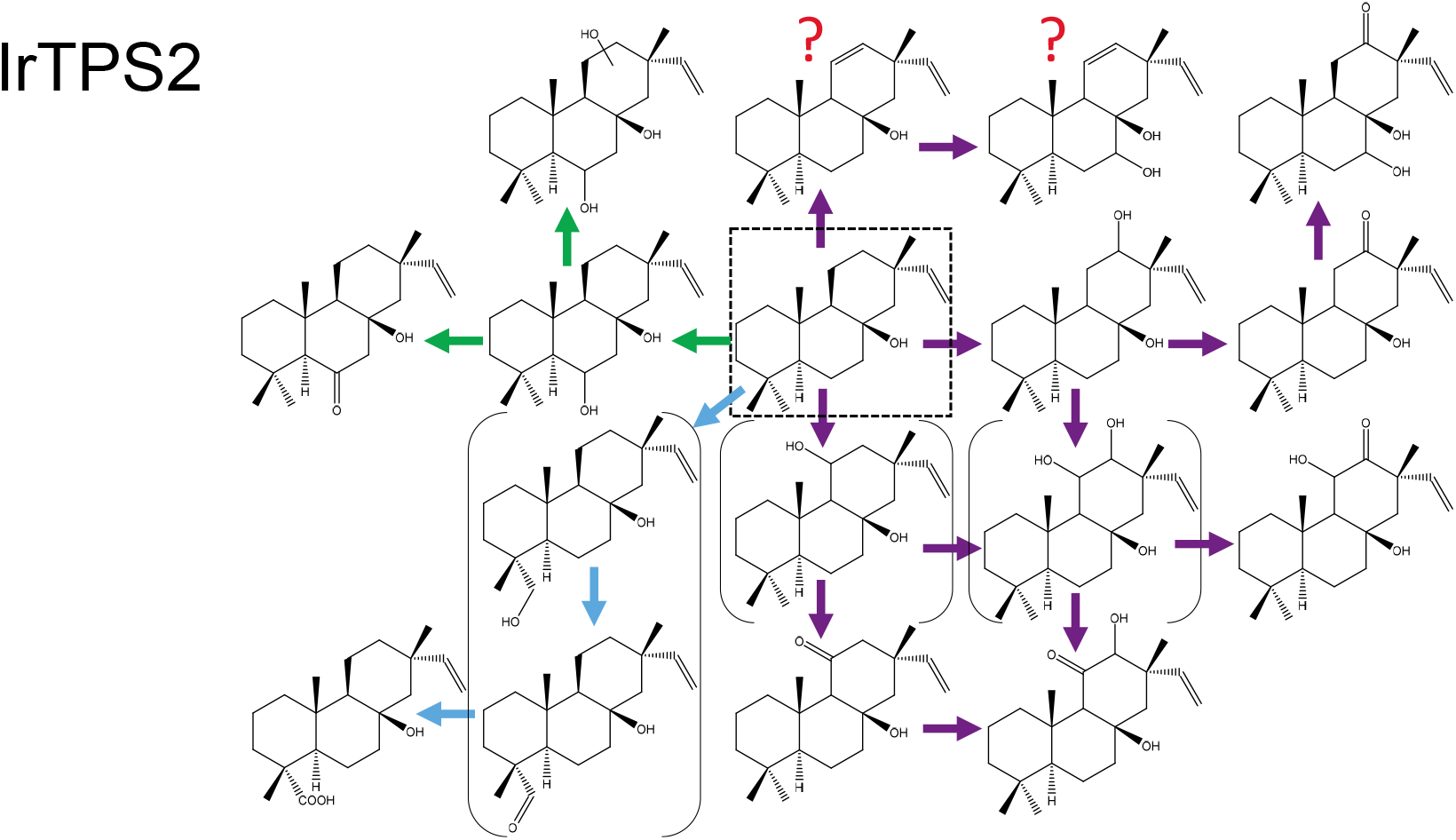
Overview of the network of products derived from nezukol (produced by IrTPS2, in the box with dashed lines). Reactions catalyzed by the CYP enzymes are indicated by the color code as in Fig. S37a. Question marks indicate hypothetical structures.

**Fig. S37e.**
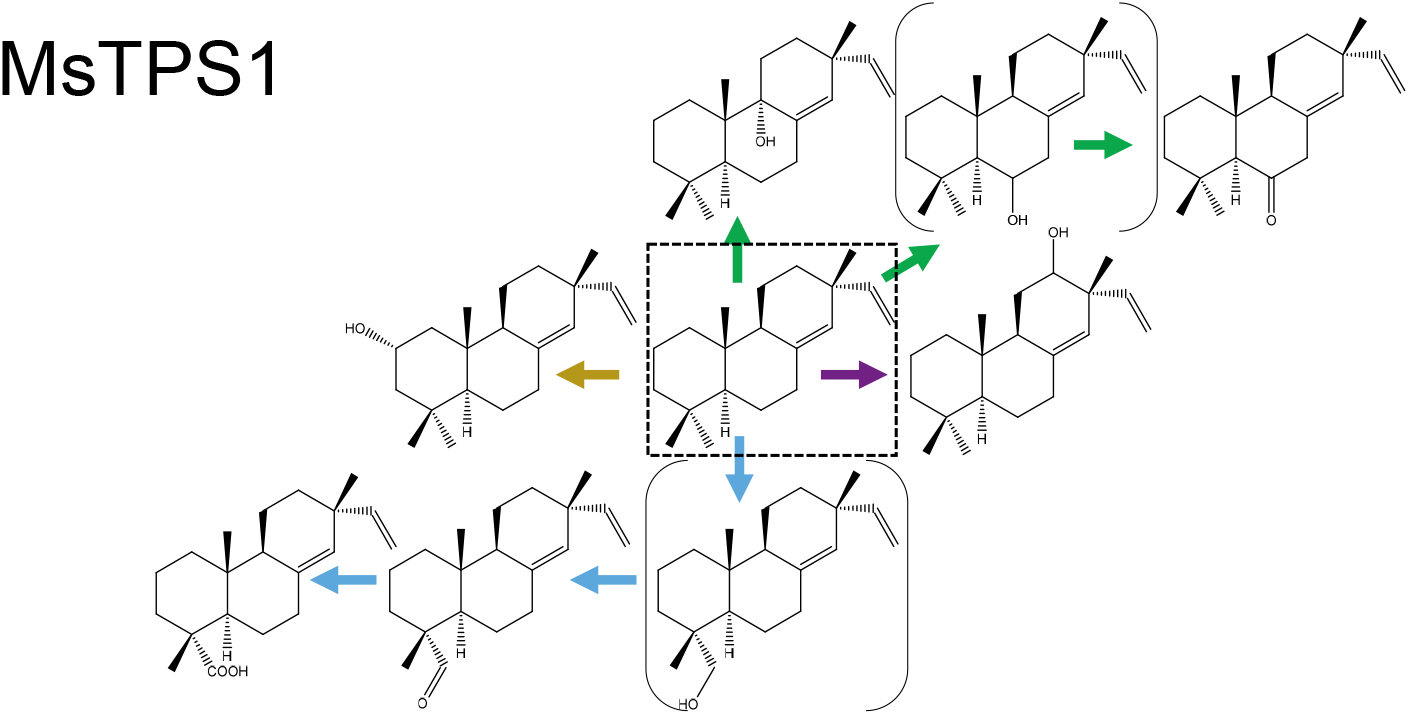
Overview of the network of products derived from sandaracopimara-8(14),15-diene (produced by MsTPS1, in the box with dashed lines). Reactions catalyzed by the CYP enzymes are indicated by the color code as in Fig. S37a.

**Fig. S37f.**
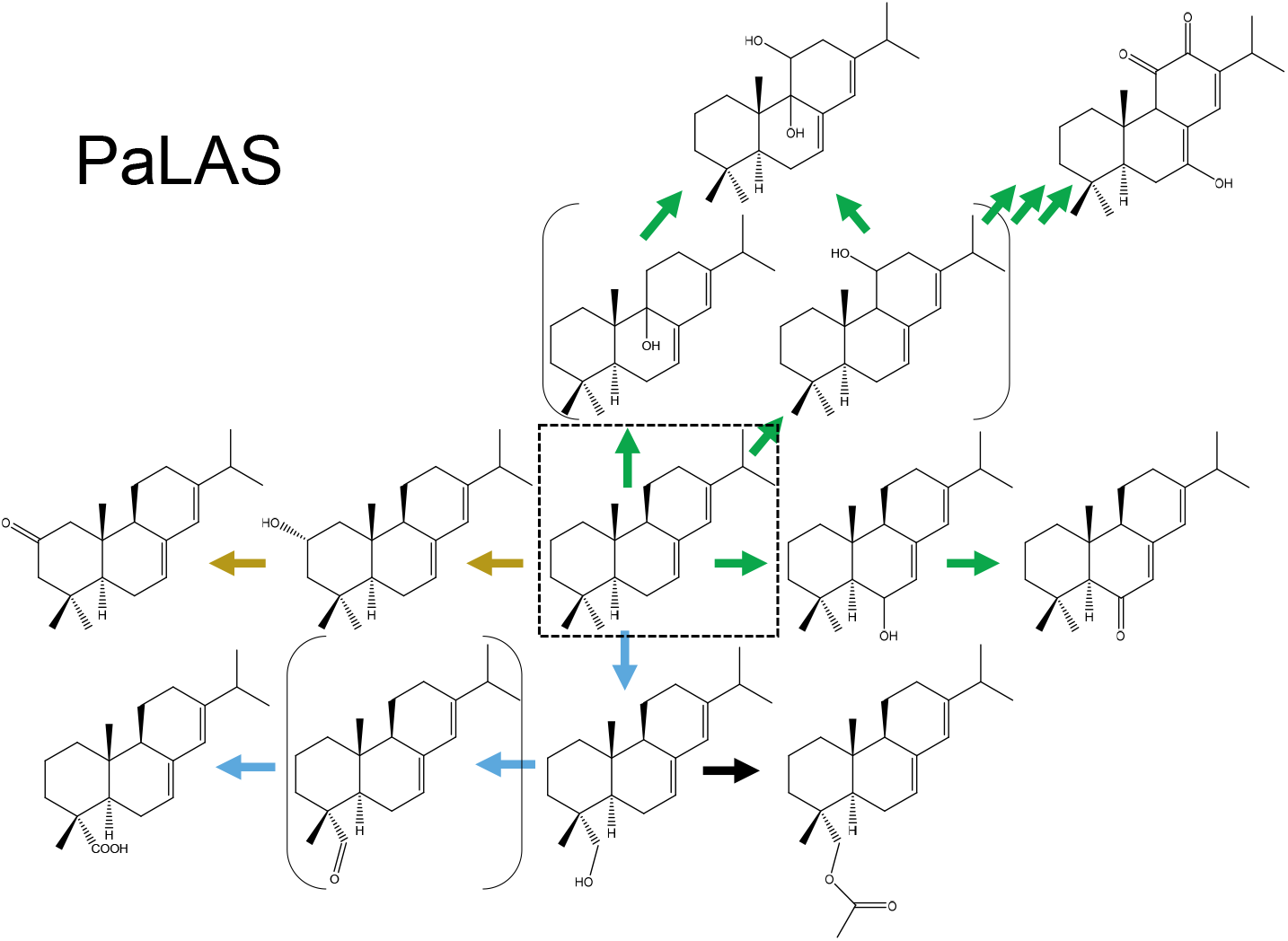
Overview of the network of products derived from abieta7(8),13-diene (produced by PaLAS, in the box with dashed lines). Reactions catalyzed by the CYP enzymes are indicated by the color code as in Fig. S37a.

**Fig. S37g.**
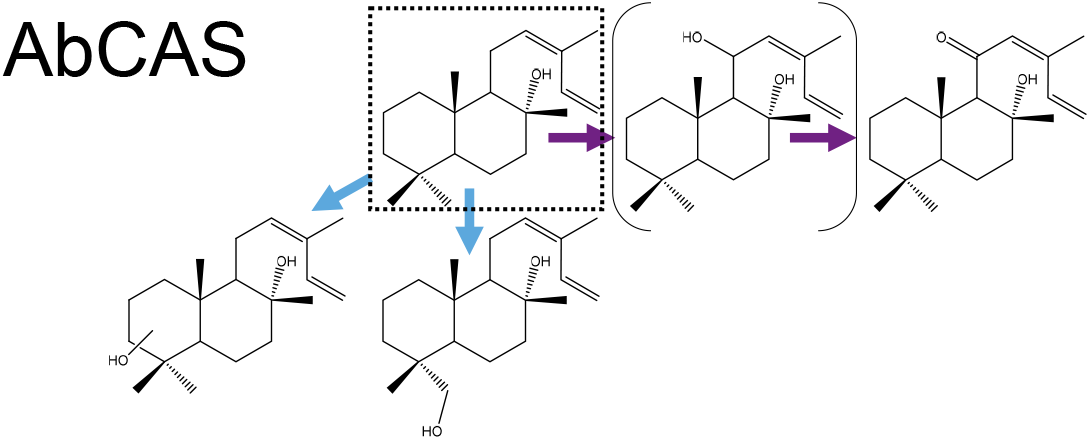
Overview of the network of products derived from cis-abienol (produced by AbCAS, in the box with dashed lines). Reactions catalyzed by the CYP enzymes are indicated by the color code as in Fig. S37a.

**Fig. S37h.**
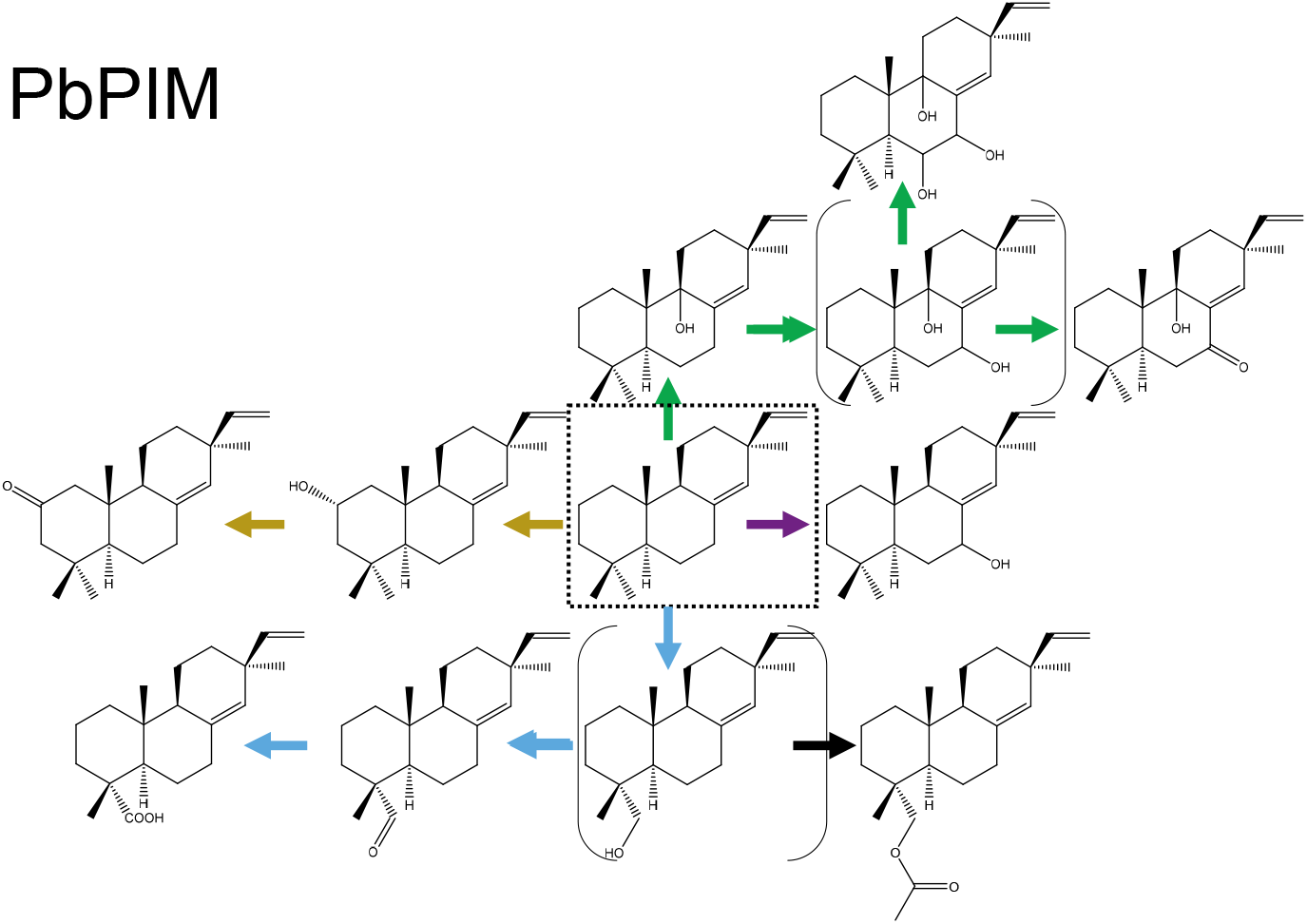
Overview of the network of products derived from pimara-8(14),15-diene (produced by PbPIM, in the box with dashed lines). Reactions catalyzed by the CYP enzymes are indicated by the color code as in Fig. S37a.

**Figure S37i.**
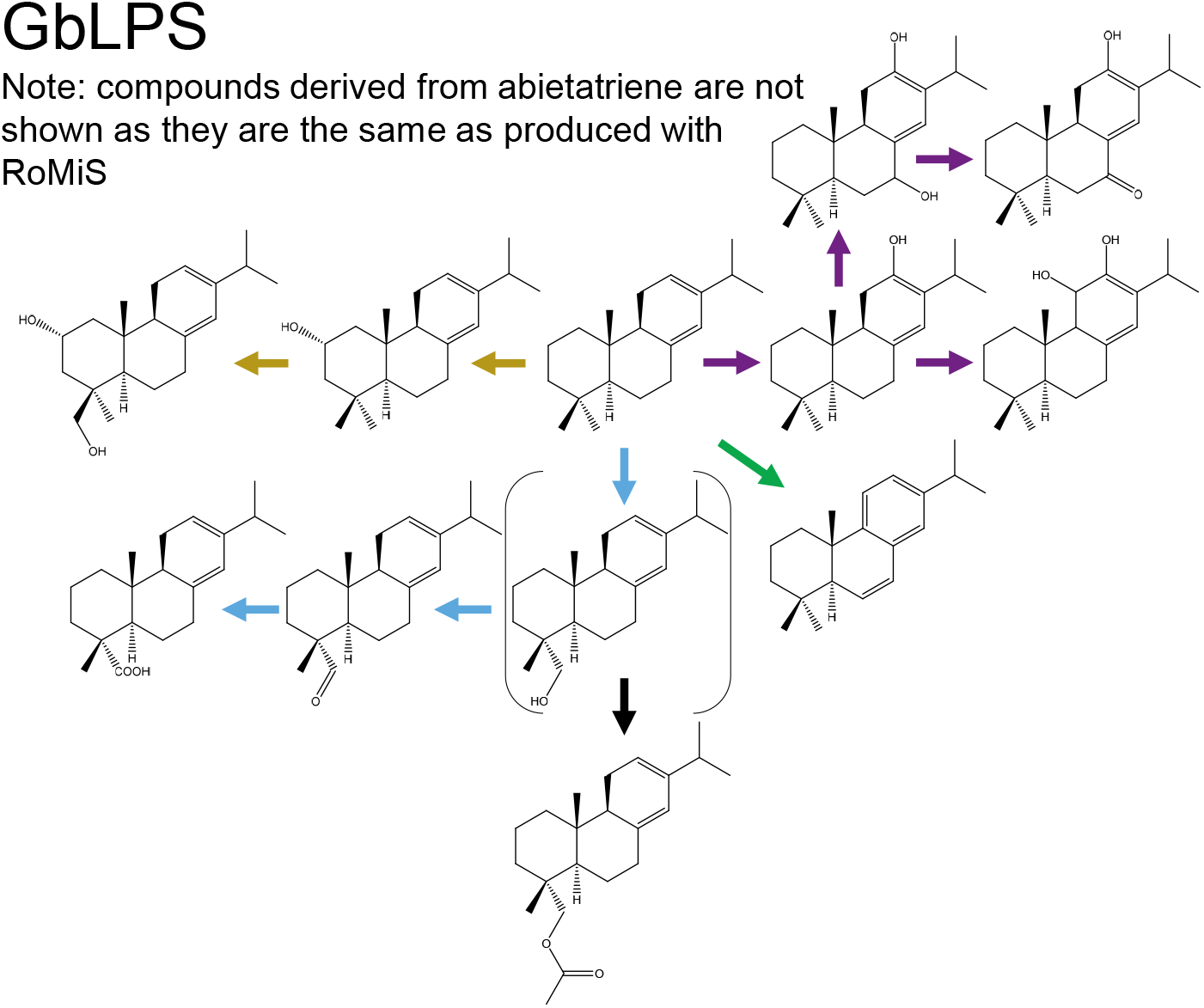
Overview of the network of products derived from levopimaradiene (produced by GbLPS, in the box with dashed lines). Reactions catalyzed by the CYP enzymes are indicated by the color code as in Fig. S37a.

**Table S1.** Genes used in this study.

| **Gene** | **Organism** | **Accession no.** | **Annotation** | **Ref.** |
| --- | --- | --- | --- | --- |
| NtGGPPS | *Nicotiana tabacum* | NP_001313025 | full length | [17] |
| RoCPS | *Rosmarinus officinalis* | AHL67261 | full length | [4d] |
| PbPIM | *Pinus banksiana* | AFU73868 | deleted transit peptide (-52 aa), yeast codon optimized | [18] |
| RoMiS | *Rosmarinus officinalis* | AHL67263 | yeast codon optimized, deleted transit peptide (-45 aa) | [4d] |
| OmTPS3 | *Origanum majorana* | AZB50371 | full length, yeast codon optimized | [19] |
| IrKSL6 | *Isodon rubescens* | ASC55318 | full length, yeast codon optimized | [20] |
| IrTPS2 | *Isodon rubescens* | ARO38140 | full length, yeast codon optimized | [21] |
| MsTPS1 | *Mentha spicata* | AZB50369 | full length, yeast codon optimized | [19] |
| GbLPS | *Ginkgo biloba* | Q947C4 | deleted transit peptide (-55 aa), yeast codon optimized | [22] |
| PaLAS | *Picea abies* | AAS47691 | full length, yeast codon optimized | [23] |
| SmCPSKSL1 | *Selaginella moellendorffii* | AEK75338 | full length, yeast codon optimized | [24] |
| AbCAS | *Abies balsamea* | XP_002513340 | full length, yeast codon optimized | [25] |
| ATR1 | *Arabidopsis thaliana* | NP_194183 | - | [4d] |
| RoCYP76AH22 | *Rosmarinus officinalis* | KP091843 | - | [4d] |
| SpCYP71BE52 | *Salvia pomifera* | KT157042 | yeast codon optimized | [4c] |
| OsCYP76M8 | *Oryza sativa* | BAG91560 | yeast codon optimized | [10a] |
| PsCYP720B4 | *Picea sitchensis* | HM245403 | yeast codon optimized | [12] |

**Table S2.** Constructs and corresponding yeast strains used in this study.

| Yeast strain | Core module | diTPS | CYP |
| --- | --- | --- | --- |
| INVSc1 | GGPPS:RoCPS:ATR1 | PbPIM | - |
|  |  |  | RoCYP76AH22 |
|  |  |  | SpCYP71BE52 |
|  |  |  | OsCYP76M8 |
|  |  |  | PsCYP720B4 |
|  |  | RoMiS | - |
|  |  |  | RoCYP76AH22 |
|  |  |  | SpCYP71BE52 |
|  |  |  | OsCYP76M8 |
|  |  |  | PsCYP720B4 |
|  |  | OmTPS3 | - |
|  |  |  | RoCYP76AH22 |
|  |  |  | SpCYP71BE52 |
|  |  |  | OsCYP76M8 |
|  |  |  | PsCYP720B4 |
|  |  | IrKSL6 | - |
|  |  |  | RoCYP76AH22 |
|  |  |  | SpCYP71BE52 |
|  |  |  | OsCYP76M8 |
|  |  |  | PsCYP720B4 |
|  |  | IrTPS2 | - |
|  |  |  | RoCYP76AH22 |
|  |  |  | SpCYP71BE52 |
|  |  |  | OsCYP76M8 |
|  |  |  | PsCYP720B4 |
|  |  | MsTPS1 | - |
|  |  |  | RoCYP76AH22 |
|  |  |  | SpCYP71BE52 |
|  |  |  | OsCYP76M8 |
|  |  |  | PsCYP720B4 |
|  | GGPPS:ATR1 | GbLPS | - |
|  |  |  | RoCYP76AH22 |
|  |  |  | SpCYP71BE52 |
|  |  |  | OsCYP76M8 |
|  |  |  | PsCYP720B4 |
|  |  | PaLAS | - |
|  |  |  | RoCYP76AH22 |
|  |  |  | SpCYP71BE52 |
|  |  |  | OsCYP76M8 |
|  |  |  | PsCYP720B4 |
|  |  | SmCPSKSL1 | - |
|  |  |  | RoCYP76AH22 |
|  |  |  | SpCYP71BE52 |
|  |  |  | OsCYP76M8 |
|  |  |  | PsCYP720B4 |
|  |  | AbCAS | RoCYP76AH22 |
|  |  |  | SpCYP71BE52 |
|  |  |  | OsCYP76M8 |
|  |  |  | PsCYP720B4 |

**Table S3.** See separate file.

References cited in Table S3:

Scheler et al, Tissier, 2016, Nat Comm.^[4d]^

Huffmann, Gibbs, 1974, J. Org. Chem^[26]^

Zhao et al., Zhou, 2016, Org. Biomol. Chem.^[27]^

Forman, et al, Hamberger, 2017, Molecules^[28]^

Noma et al, Kawashima, 1982, Phytochemistry^[29]^

Xu et al., Ma, 2011, Fitoterapia^[30]^

Popova, et al, Bankova, 2010, J Agr. F Chem^[31]^

Jin et al., Huang, 2017, Plant Phys.^[32]^

Mafu et al, Peters, 2016, PNAS^[10c]^

Pateraki et al., Hamberger, 2017, eLife^[4b]^

Wang et al, Peters, 2012, JBC^[33]^

Chernenko et al, Gatilov, 1991, Chemistry of Natural Compounds^[34]^

Piovano et al, Pascard, 1988, Phytochemistry^[35]^

Pelot, et al, Zerbe, 2017, PLoS ONE^[36]^

Bohlmann, Zdero, 1982, Phytochemistry^[37]^

Bohlmann et al, Zdero, 1973, Chemische Berichte^[38]^

Tasnim, et al, Mattsson, 2020, Plants^[39]^

Wang et a., Peters, 2012, J. Biol. Chem.^[10a]^

Nagashima, et al, Asakawa, 2003, Phytochemistry^[40]^

Ro, Bohlmann, 2006, Phytochemistry^[41]^

Mafu et al, Peters, 2011, Chembiochem^[24]^

Sallaud et al, Tissier, 2012, Plant J^[42]^

Zerbe et al, Bohlmann, 2012, J Biol Chem^[25]^

Hall et al, Bohlmann, 2013, Plant Physiol^[18]^

Chang et al, Kuo, 2005, Chem Pharm Bull^[43]^

Ulubelen and Topcu, 1992, Phytochemistry^[44]^

Geiwiz and Haslinger, 1995, Helvetica Chimica Acta^[45]^

Hamberger et al., Bohlmann, 2011, Plant Physiol.^[12]^

**Table S4.** See separate file.

References cited in Table S4:

Scheler et al, Tissier, 2019, J. Agr. Food Chem.^[10b]^

**Table S5.** Assignment of ^1^H-NMR Signals (600 MHz, CDCl_3_, 298 K) of 38.

| H at relevant<br>C atom<sup>a</sup> | δ [ppm]<sup>b</sup> | I<sup>c</sup> | M<sup>c</sup> | J [Hz]<sup>c</sup> | gs-COSY<sup>d</sup> |
| --- | --- | --- | --- | --- | --- |
| H-C(1<sub>α</sub>) | 0.74-0.81 | 1 | m | 3.4/12.9 | H-C(1<sub>β</sub>) |
| H-C(5) | 0.78-0.83 | 1 | m | - | H-C(6) |
| CH<sub>3</sub>(18) | 0.88 | 3 | s | - | - |
| CH<sub>3</sub>(19) | 0.88 | 3 | s | - | - |
| H-C(3<sub>α</sub>) | 1.11-1.18 | 1 | ddd | 4.0/13.5/17.4 | H-C(3<sub>β</sub>) |
| CH<sub>3</sub>(17) | 1.26 | 3 | s | - | - |
| CH<sub>3</sub>(20) | 1.38 | 3 | s | - | - |
| H-C(2<sub>α</sub>) | 1.36-1.42 | 1 | m | - | H-C(2<sub>β</sub>) |
| H-C(3<sub>β</sub>) | 1.39-1.45 | 1 | m | - | H-C(3<sub>α</sub>) |
| H-C(6) | 1.55-1.61 | 2 | m | - | H-C(5) |
| H-C(7<sub>α</sub>) | 1.57-1.63 | 1 | m | - | H-C(7<sub>β</sub>) |
| H-C(2<sub>β</sub>) | 1.65-1.74 | 1 | m | - | H-C(2<sub>α</sub>) |
| H-C(14<sub>α</sub>) | 1.69-1.74 | 1 | m | - | H-C(14<sub>β</sub>) |
| H-C(14<sub>β</sub>) | 1.82-1.87 | 1 | m | - | H-C(14<sub>α</sub>) |
| H-C(7<sub>β</sub>) | 1.83-1.88 | 1 | m | - | H-C(7<sub>α</sub>) |
| H-C(9) | 2.17 | 1 | s | - | - |
| H-C(12<sub>α</sub>) | 2.16-2.21 | 1 | m | - | H-C(12<sub>β</sub>) |
| H-C(1<sub>β</sub>) | 2.23-2.28 | 1 | m | 3.4/4.7/12.9 | H-C(1<sub>α</sub>) |
| H-C(12<sub>β</sub>) | 2.46 | 1 | d | 12.6 | H-C(12<sub>α</sub>) |
| H-C(15) | 4.93 | 2 | dd | 10.7/17.3 | H-C(16) |
| H-C(16) | 5.80 | 1 | ddd | 1.3/10.7/17.3 | H-C(15) |

<sup>a</sup>Numbering of carbon atoms refers to formula shown above. <sup>b</sup>The chemical shifts (ppm) are given in relation to DMSO-d<sub>6</sub>. <sup>c</sup>Determined from 1D spectrum. Observed homonuclear H,H correlations.

**Table S6.** Assignment of ^13^C-NMR Signals (150 MHz, CDCl_3_, 298 K) of 38.

| C-Atom<sup>a</sup> | $\delta$ [ppm]<sup>b</sup> | 135°-DEPT<sup>c</sup> | HSQC [$^1J_{\text{H,C}}$]<sup>d</sup> | HMBC [$^{2,3,4}J_{\text{H,C}}$]<sup>d</sup> |
| --- | --- | --- | --- | --- |
| C(20) | 15.6 | $\text{CH}_3$ | H-C(20) | H-C(1), H-C(5), H-C(9) |
| C(6) | 17.6 | $\text{CH}_2$ | H-C(6) | H-C(5), H-C(7) |
| C(2) | 18.0 | $\text{CH}_2$ | H-C(2) | H-C(1), H-C(3), H-C(18), H-C(19) |
| C(18) | 21.8 | $\text{CH}_3$ | H-C(18) | H-C(3), H-C(5), H-C(19), H-C(20)* |
| C(17) | 25.4 | $\text{CH}_3$ | H-C(17) | H-C(12), H-C(14), H-C(15) |
| C(4) | 33.2 | C |  | H-C(2), H-C(3), H-C(5), H-C(18),<br>H-C(19) |
| C(19) | 33.8 | $\text{CH}_3$ | H-C(19) | - |
| C(10) | 36.8 | C |  | H-C(1), H-C(6), H-C(9), H-C(15),<br>H-C(20) |
| C(1) | 38.8 | $\text{CH}_2$ | H-C(1) | H-C(2), H-C(3), H-C(9), H-C(20) |
| C(13) | 41.3 | C |  | H-C(12), H-C(14), H-C(15), H-C(16), H-C(17) |
| C(3) | 42.0 | $\text{CH}_2$ | H-C(3) | H-C(1), H-C(2), H-C(5), H-C(18),<br>H-C(19) |
| C(7) | 43.3 | $\text{CH}_2$ | H-C(7) | H-C(5), H-C(6), H-C(14) |
| C(14) | 50.7 | $\text{CH}_2$ | H-C(14) | H-C(9), H-C(12), H-C(15), H-C(17) |
| C(12) | 54.2 | $\text{CH}_2$ | H-C(12) | H-C(14), H-C(15), H-C(17) |
| C(5) | 55.9 | CH | HC(5) | H-C(3), H-C(6), H-C(7), H-C(9),<br>H-C(18), H-C(19), H-C(20) |
| C(9) | 66.6 | CH | H-C(9) | H-C(1), H-C(12), H-C(14), H-C(20) |
| C(8) | 78.1 | C |  | H-C(6), H-C(7), H-C(9), H-C(14) |
| C(16) | 109.8 | $\text{CH}_2$ | H-C(16) | |
| C(15) | 148.7 | CH | H-C(15) | H-C(12), H-C(14), H-C(17) |
| C(11) | 209.6 | C |  | H-C(9), H-C(12) |

<sup>a</sup>Numbering of carbon atoms refers to formula shown above. <sup>b</sup>The chemical shifts (ppm) are given in relation to $\text{CDCl}_3$ . <sup>c</sup>Chemical shifts of the quaternary carbon atoms are taken from the difference of the $^{13}\text{C}$ spectrum and the 135°-DEPT spectrum. <sup>d</sup>Observed heteronuclear H,C correlations.

**Table S7.** Assignment of ^1^H-NMR Signals (600 MHz, CDCl_3_, 298 K) of 120.

| H at relevant C atom<sup>a</sup> | $\delta$ [ppm]<sup>b</sup> | I<sup>c</sup> | M<sup>c</sup> | $J$ [Hz]<sup>c</sup> | gs-COSY<sup>d,e</sup> |
| --- | --- | --- | --- | --- | --- |
| H-C(1<sub>α</sub>) | 0.94-1.01 | 1 | m | - | H-C(1<sub>β</sub>), H-C(2<sub>β</sub>) |
| H-C(9) | 0.96-1.00 | 1 | m | - | - |
| CH<sub>3</sub>(20) | 1.05 | 3 | s | - | - |
| H-C(6<sub>α</sub>) | 1.12-1.17 | 1 | m | - | H-C(6<sub>β</sub>), H-C(7<sub>α</sub>) |
| CH<sub>3</sub>(19) | 1.22 | 3 | s | - | - |
| CH<sub>3</sub>(17) | 1.24 | 3 | s | - | - |
| H-C(12<sub>α</sub>) | 1.31-1.38 | 1 | m | - | - |
| H-C(14) | 1.35-1.38 | 2 | m | - | - |
| H-C(7<sub>α</sub>) | 1.43-1.50 | 1 | m | - | H-C(6<sub>α</sub>) |
| H-C(11<sub>α</sub>) | 1.45-1.53 | 1 | m | - | - |
| H-C(2<sub>α</sub>) | 1.53-1.59 | 1 | m | - | - |
| H-C(12<sub>β</sub>) | 1.56-1.61 | 1 | m | - | - |
| H-C(7<sub>β</sub>) | 1.61-1.66 | 1 | m | - | - |
| H-C(3<sub>α</sub>) | 1.62-1.67 | 1 | m | - | - |
| H-C(11<sub>β</sub>) | 1.65-1.69 | 1 | m | - | - |
| H-C(2<sub>β</sub>) | 1.66-1.70 | 1 | m | - | H-C(1<sub>α</sub>) |
| H-C(6<sub>β</sub>) | 1.70-1.79 | 1 | m | - | H-C(6<sub>α</sub>) |
| H-C(5) | 1.73-1.77 | 1 | m | - | - |
| H-C(1<sub>β</sub>) | 1.74-1.80 | 1 | m | - | H-C(1<sub>α</sub>) |
| H-C(3<sub>β</sub>) | 1.74-1.83 | 1 | m | - | - |
| H-C(16) | 4.86 | 2 | ddd | 1.2/10.8/17.5 | H-C(12), H-C(15) |
| H-C(15) | 5.74 | 1 | dd | 10.8/17.5 | H-C(16) |

<sup>a</sup>Numbering of carbon atoms refers to formula shown above. <sup>b</sup>The chemical shifts (ppm) are given in relation to DMSO-d<sub>6</sub>. <sup>c</sup>Determined from 1D spectrum. Observed homonuclear H,H correlations. <sup>d</sup>Due to the strong overlap of the signals, assignment of correlations by COSY is not possible.

**Table S8.** Assignment of ^13^C-NMR Signals (150 MHz, CDCl_3_, 298 K) of 120.

| C-Atom<sup>a</sup> | $\delta$ [ppm]<sup>b</sup> | 135°-DEPT<sup>c</sup> | HSQC [$^1J_{\text{H,C}}$]<sup>d</sup> | HMBC [$^{2,3,4}J_{\text{H,C}}$]<sup>d</sup> |
| --- | --- | --- | --- | --- |
| C(20) | 15.9 | $\text{CH}_3$ | H-C(20) | H-C(1), H-C(5), H-C(20) |
| C(19) | 16.2 | $\text{CH}_3$ | H-C(19) | H-C(3), H-C(5) |
| C(11) | 17.0 | $\text{CH}_2$ | H-C(11) | H-C(9), H-C(12) |
| C(2) | 17.6 | $\text{CH}_2$ | H-C(2) | H-C(1), H-C(3), H-C(19) |
| C(6) | 20.8 | $\text{CH}_2$ | H-C(6) | H-C(5), H-C(7) |
| C(17) | 24.2 | $\text{CH}_3$ | H-C(17) | H-C(14), H-C(15) |
| C(13) | 36.5 | C | - | H-C(11), H-C(12), H-C(14), H-C(15), H-C(16), H-C(17) |
| C(10) | 36.6 | C | - | H-C(1), H-C(5), H-C(9), H-C(20) |
| C(3) | 37.0 | $\text{CH}_2$ | H-C(3) | H-C(1), H-C(19) |
| C(12) | 38.0 | $\text{CH}_2$ | H-C(12) | H-C(14), H-C(15), H-C(17) |
| C(1) | 38.5 | $\text{CH}_2$ | H-C(1) | H-C(2), H-C(3), H-C(9), H-C(20) |
| C(7) | 43.1 | $\text{CH}_2$ | H-C(7) | H-C(5), H-C(6), H-C(14) |
| C(4) | 47.4 | C | - | H-C(2), H-C(3), H-C(5), H-C(6), H-C(19) |
| C(5) | 50.5 | CH | H-C(5) | H-C(1), H-C(3), H-C(6), H-C(7), H-C(9), H-C(19), H-C(20) |
| C(14) | 51.6 | $\text{CH}_2$ | H-C(14) | H-C(15), H-C(17) |
| C(9) | 57.0 | CH | H-C(9) | H-C(1), H-C(5), H-C(11), H-C(12), H-C(14), H-C(20) |
| C(8) | 72.8 | C | - | H-C(6), H-C(7), H-C(9), H-C(11), H-C(14) |
| C(16) | 108.7 | $\text{CH}_2$ | H-C(16) | H-C(17) |
| C(15) | 151.5 | CH | H-C(15) | H-C(12), H-C(14) |
| C(18) | 181.9 | C | - | H-C(3), H-C(5), H-C(19) |

<sup>a</sup>Numbering of carbon atoms refers to formula shown above. <sup>b</sup>The chemical shifts (ppm) are given in relation to $\text{CDCl}_3$ . <sup>c</sup>Chemical shifts of the quaternary carbon atoms are taken from the difference of the $^{13}\text{C}$ spectrum and the 135°-DEPT spectrum. <sup>d</sup>Observed heteronuclear H,C correlations.

**Table S9.**
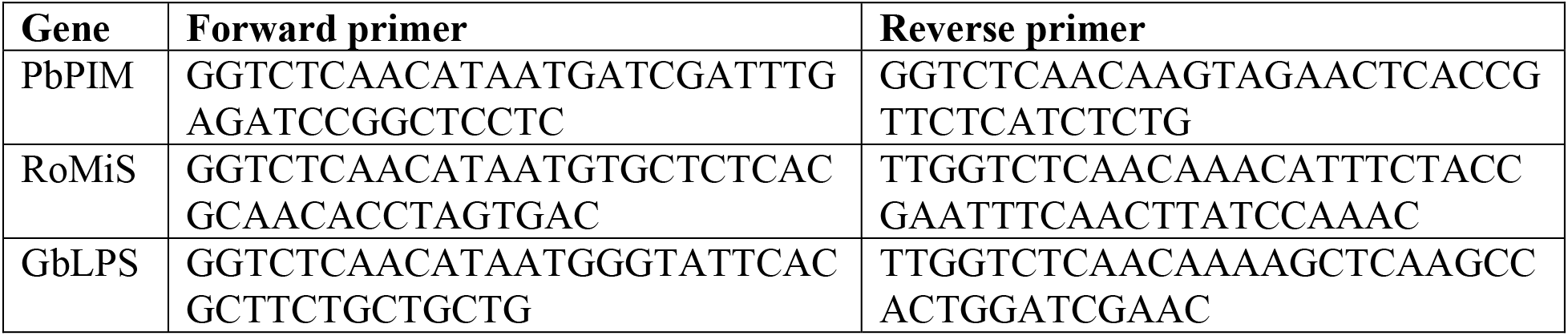
Primers used to delete transit peptides of PbPIM, RoMiS and GbLPS.

